# Brønsted-basic small molecules activate GTP hydrolysis in Ras Q61 mutants

**DOI:** 10.1101/2025.11.04.686643

**Authors:** Ye-Cheng Wang, Si-Cong Chen, Yang Wu, Yang Cao, Zhongda Shi, Michael A. Norinskiy, Celine D. Wang, Hasan Celik, Ziyang Zhang

## Abstract

The *RAS* oncogenes (*KRAS*, *HRAS*, *NRAS*) are among the most frequently mutated genes in human cancer, affecting over three million patients annually. Therapeutic development has largely focused on inhibitors for *KRAS* codon 12 mutations (G12C/D/V/S/R) which are key drivers in lung, colorectal, and pancreatic cancers. In contrast, mutant-selective inhibitors for Q61 variants remain elusive. A common mechanistic feature of G12 and Q61 mutants is the impaired hydrolysis of GTP, which traps Ras in its active, signaling-competent state. We envisioned that an alternative therapeutic strategy – reactivation of GTP hydrolysis – could address this shared oncogenic mechanism. Here we report small molecules that accelerate GTP hydrolysis in K-Ras Q61 mutants. These compounds compensate for the loss of the catalytic residue Gln61 by introducing a general base into the active site, selectively enhancing hydrolysis of K-Ras Q61X (X = H, L, K, R) mutants by up to 20-fold while sparing the wild-type protein. In mutant cancer cell lines, these compounds reduce GTP-bound Ras levels and suppress downstream signaling. We show that the chemical design principles are generalizable to other Ras isoforms. This work establishes a mechanistic foundation for small-molecule “GTPase activators” and offers a new paradigm for targeting Ras-driven cancers.

## Introduction

The *RAS* genes (*KRAS*, *HRAS*, *NRAS*) are the most frequently mutated oncogenes in human cancer, collectively affecting more than 3 million patients every year^1^. Although oncogenic *RAS* was considered “undruggable”, recent breakthroughs in mutant-selective *KRAS G12C* inhibitors have demonstrated the feasibility of direct driver-oncogene inhibition and translated to patient benefit^2–6^. With two oral KRAS G12C inhibitors in clinical use and many more in clinical trials^7–15^, current research efforts have focused on addressing other oncogenic mutants and improving the depth and duration of inhibition.

The vast majority of current efforts are directed toward *KRAS* hotspot mutations at codon 12 (*KRAS* G12C/D/V/S/R)^16–42^, which are common driver mutations in lung, colorectal and pancreatic cancers. In contrast, mutant-selective inhibitors of Q61 mutants remain elusive and represent an area of unmet need. Hotspot mutations at codon 61 (Q61H/L/K/R) are much more common in mutant *HRAS* and *NRAS*-driven cancers compared to *KRAS*-driven cancers, notably head and neck cancer and melanoma, accounting for about 60% and 85% of total cases, respectively^43^.

A common mechanistic feature of these oncogenic mutants, particularly at codons 12 and 61, is the impaired hydrolysis of GTP, which traps Ras in its active, signaling-competent state. This defect is most severe for Q61 mutants, as glutamine 61 is an indispensable component of the catalytic machinery for both intrinsic and GTPase-accelerating protein (GAP)-mediated GTP hydrolysis^44–48^. Q61 mutants are thought to be completely devoid of GTPase activity, precluding access to the GDP-bound state^49^. In addition, the Q61R mutation affects the dynamics of the Switch II region, rendering the Switch II pocket commonly targeted by inhibitors less accessible^50^.

An alternative therapeutic strategy for Ras Q61 mutants is to activate their enzymatic activity to promote their transition to the signaling-incompetent GDP state^51^. Such a strategy was first explored by Wittinghofer, Scheffzek and coworkers, who showed that a modified GTP nucleotide was hydrolyzed >200 times faster by oncogenic Ras mutants^52,53^. While unlikely to be a therapeutic approach, this finding supports the feasibility of restoring GTPase activity of Ras.

Here we report the design, synthesis and functional characterization of small molecule ligands that directly bind to Ras and accelerate the hydrolysis of GTP for Q61 mutants. Our strategy is based on the principle of chemical complementation, where an exogenous small molecule rescues a catalytically dead enzyme by providing an alternative reaction mechanism, which has been applied to aminotransferases^54^, proteases^55,56^, kinases^57^, among others^58^. We show that the loss of the catalytic residue Gln61 in oncogenic mutants can be rescued by chemically supplementing the active site with a general base. These compounds accelerate the rate of GTP hydrolysis by Ras Q61X (X = H, L, K, R) mutants to near wild-type levels (up to 20-fold increase) without affecting that of the wildtype protein and reduce the levels of GTP-bound Ras in mutant cancer cell lines.

## Results

To devise a chemical strategy to restore GTP hydrolysis in Ras Q61 mutants, we first surveyed the natural mechanisms for GTP hydrolysis in small GTPases. Among these, the translation-associated GTPases (eEF1A in humans, EF-Tu in bacteria) are unique in that they utilize a histidine residue (His87, EF-Tu numbering) in lieu of glutamine to coordinate the nucleophilic water molecule, assisted by the 5’-phosphate of A2662 on the large ribosomal subunit ^59^. We hypothesized that such a mechanism could be reconstituted chemically by using a small molecule ligand to simultaneously present a general base in a catalytically competent orientation (Fig. 1a).

**Fig. 1.**
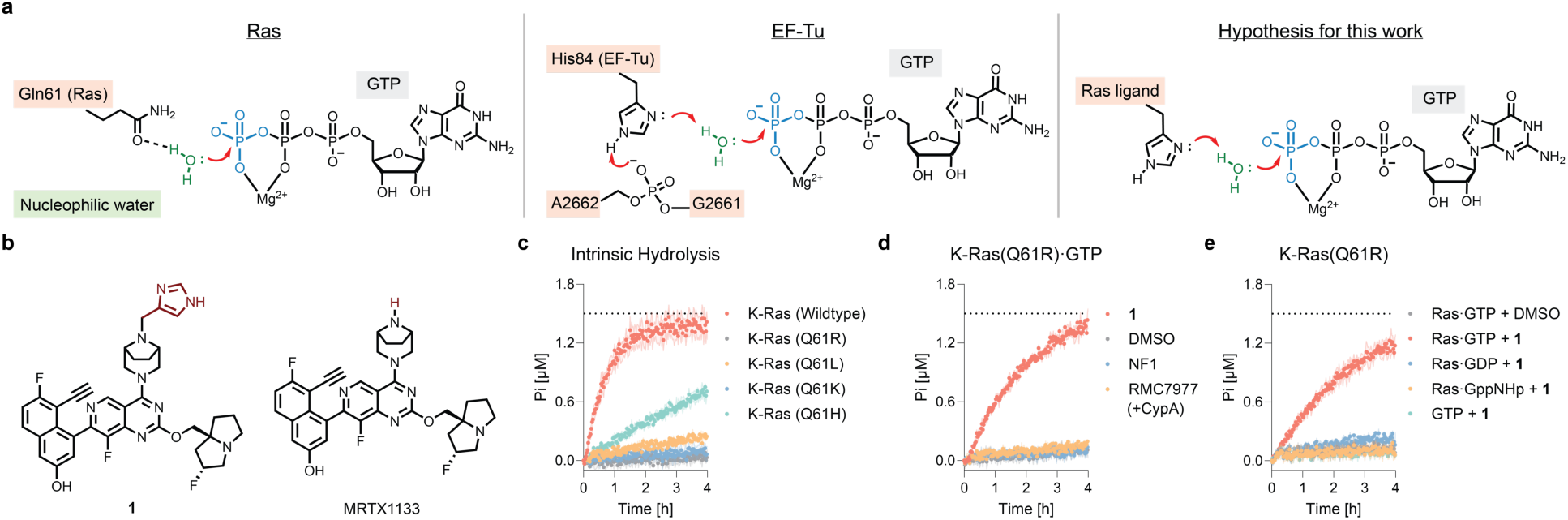
Synthetic chemical ligands activate the GTP hydrolysis of K-Ras (Q61R). **a.** Catalytic mechanisms for GTP hydrolysis of Ras family small GTPases, the translation-associated GTPase EF-Tu, and the mechanistic rationale for small molecule GTP hydrolysis potentiators in this work. **b.** Chemical structures of compound **1** and the parental compound MRTX1133. **c.** Intrinsic GTP hydrolysis rates of wildtype K-Ras and Q61 mutants. **d.** Compound **1**, but not the NF1 GRD or the ternary complex inhibitor RMC7977/cyclophilin A, accelerates GTP hydrolysis of K-Ras(Q61R). **e**. The effect of compound **1** is selective for the GTP-bound state.

To test this hypothesis, we synthesized compound **1**, which contains the chemical scaffold of MRTX 1133, which is privileged for binding the Switch II pocket in K-Ras^60^, as well as a pendent imidazole group on the piperazine ring (Fig. 1b). We chose the Switch II pocket ligands because they provide ready access to the γ-phosphate of GTP. We first measured the intrinsic single-turnover GTP hydrolysis rate of four K-Ras mutants found in cancer – Q61K, Q61H, Q61L and Q61R. Consistent with literature reports, the Q61H mutant displayed weak GTPase activity, whereas the Q61K/L/R mutants were completely inactive^49,61^ (Fig. 1c). Such an observation suggests that His61 in Q61H may play a role in supporting GTP hydrolysis, albeit much less efficiently than Gln61 in wildtype K-Ras. To assess the activity of our compounds, we first focused on K-Ras Q61R, a mutant most recalcitrant to GTP hydrolysis which exists exclusively in the hydrolysis-incompetent “State 2”^62^. Treatment of K-Ras(Q61R)•GTP with compound **1** (50 µM) initiated GTP hydrolysis (Fig. 1d), whereas neither the natural GAP protein neurofibromin 1 (NF1) nor the ternary complex inhibitor RMC7977^63,64^ had any measurable effect (Fig. 1d). The parental structure of compound **1**, MRTX1133, also did not stimulate GTP hydrolysis (Extended Data Fig. 1). The effect was specific for GTP-bound K-Ras, as the hydrolysis of GDP-or GppNHp-bound K-Ras(Q61R), as well as free GTP, were not affected by the addition of compound **1** (Fig. 1e).

To assess the perturbation of the chemical environment and directly monitor the GTP hydrolysis of K-Ras(Q61R) in the presence of compound **1**, we used ^31^P NMR to measure the chemical shift changes of the α-,β- and γ-phosphoryl groups in GTP (Fig. 2a). Addition of compound **1** to K-Ras(Q61R) complexed with the non-hydrolyzable analog GppNHp revealed small but measurable chemical shifts of all three phosphoryl nuclei (Fig. 2b). Upon binding compound **1**, the β-and α-phosphoryl signals of GppNHp were shifted downfield from −0.63 ppm to −0.06 ppm (Δδ = 0.57 ppm) and from −11.92 ppm to −11.70 ppm (Δδ = 0.22 ppm), respectively, compared with the DMSO-treated sample. The γ-phosphoryl signal of GppNHp was more strongly perturbed, shifting from −2.96 ppm to −2.16 ppm (Δδ = 0.80 ppm). These chemical shift perturbations indicate that compound **1** binds close to the nucleotide, consistent with the reported X-ray co-crystal structures of Switch-II pocket ligands bound to Ras in the GTP state (PDB ID: 7T47).

**Fig. 2.**
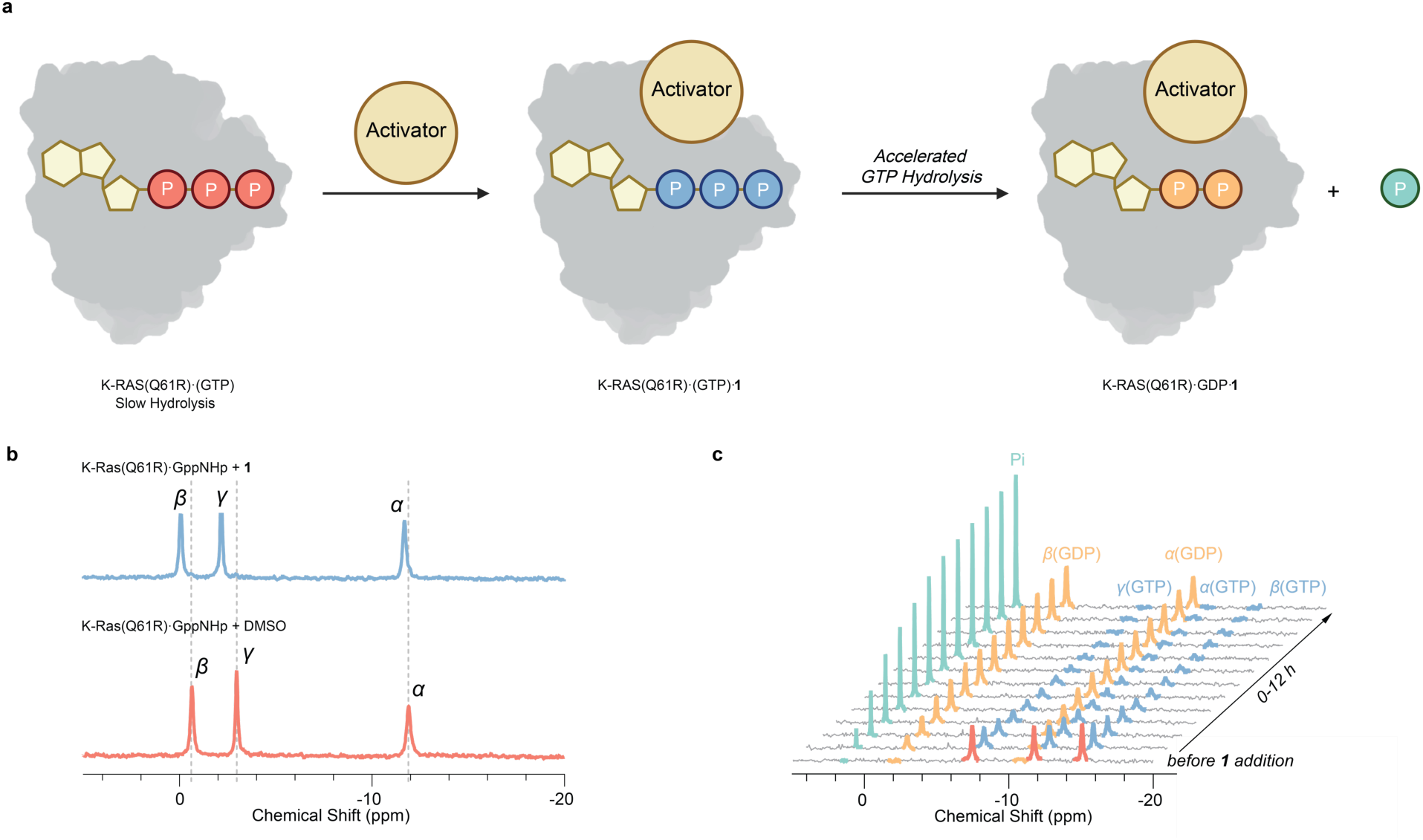
Detection of time-dependent hydrolysis of GTP by ³¹P NMR spectroscopy. **a.** Cartoon illustration of activator-induced GTP hydrolysis on K-Ras(Q61R). **b**. ^31^P NMR analysis of K-Ras(Q61R)•GppNHp treated with compound **1** (2.0 equiv.) and DMSO. **c**. Time-dependent ^31^P NMR spectra monitoring K-Ras(Q61R)•GTP hydrolysis catalyzed by compound **1** (2.0 equiv.) from 0–12 h.

We next carried out ^31^P NMR experiments with K-Ras(Q61R)•GTP and compound **1**, collecting and integrating data every hour to achieve sufficient signal-to-noise ratio. We observed a time-dependent reduction of the GTP signal and the concomitant formation of GDP and free phosphate (Fig. 2c). By contrast, DMSO-treated K-Ras(Q61R)•GTP showed no measurable GTP hydrolysis over the same time period (Extended Data Fig. 2). In the early stage (0-1 h), the α-, β-, and γ-phosphoryl signals were slightly perturbed compared with DMSO-treated condition (Fig. 2c and Extended Data Fig. 2b), consistent with our observations with the GppNHp-bound protein. Over the course of 10 h, GTP was completely hydrolyzed as indicated by the emergence of free phosphate (1.51 ppm), the new α-(−10.78 ppm) and β-(−2.00 ppm) phosphoryl signals^65^ of the K-Ras(Q61R)•GDP·**1** complex, and the disappearance of GTP peaks (Fig. 2c).

To probe the mechanistic basis of compound **1**’s activity, we synthesized 16 derivatives varying the pKa and steric bulk of the appendage (Fig. 3a). We first prepared compounds **2** and **3**, where we methylated either nitrogen on the imidazole ring, and found that while **2** retained activity, **3** was completely inactive (Fig. 3b). When we replaced the methylene linker with a carbonyl group, the resulting compound **4** was also inactive. These results led us to conclude that an appropriately positioned basic nitrogen is required for compound activity. As the lone pair of electrons of imidazole can act both as a nucleophile (as in phosphodiesterases) or a general base (as in EF-Tu) (Extended Data Fig. 3), then we prepared tetrazole **5**, thiazole **6** and oxazole **7**, which retained the p-lone pair on the nitrogen but have reduced basicity (Fig. 3c). The resulting compounds showed diminished activity compared to imidazole **1** as the p*K*a of their conjugate acid decreases. In addition, we found that substituting the imidazole group for other basic functions such as a pyridine (**8**) and a carboxylic acid (**9**) also yielded active compounds, consistent with the hypothesis that our compounds act through a general base mechanism (Fig. 3d). Similar to the imidazole series (**1-4**), we observed that the positioning of the basic nitrogen was critical for activity, as neither the *para-* nor *ortho-*isomers of **8** were active (Extended Data Fig. 4). Interestingly, the addition of steric bulk (methyl, **10**) or an electron-withdrawing group (nitro, **11**) on the imidazole ring did not significantly change the activity (Fig. 3e). We plotted the kinetic constant of compound-mediated GTP hydrolysis against the calculated p*K*_a_^66^ of the pendent R group (Fig. 3f, Supplemental Fig. S1 and Supplemental Table S1). This analysis revealed a trend across all tested compounds with an optimal p*K*_a_ value of 7, regardless of the chemotype. While we cannot rule out the possibility of a mechanism involving direct nucleophilic attack on the γ-phosphate of GTP, the observation that poorly nucleophilic compounds such as carboxylic acid **9** and triazole **12** retain activity suggests general acid-base catalysis is more likely.

**Fig. 3.**
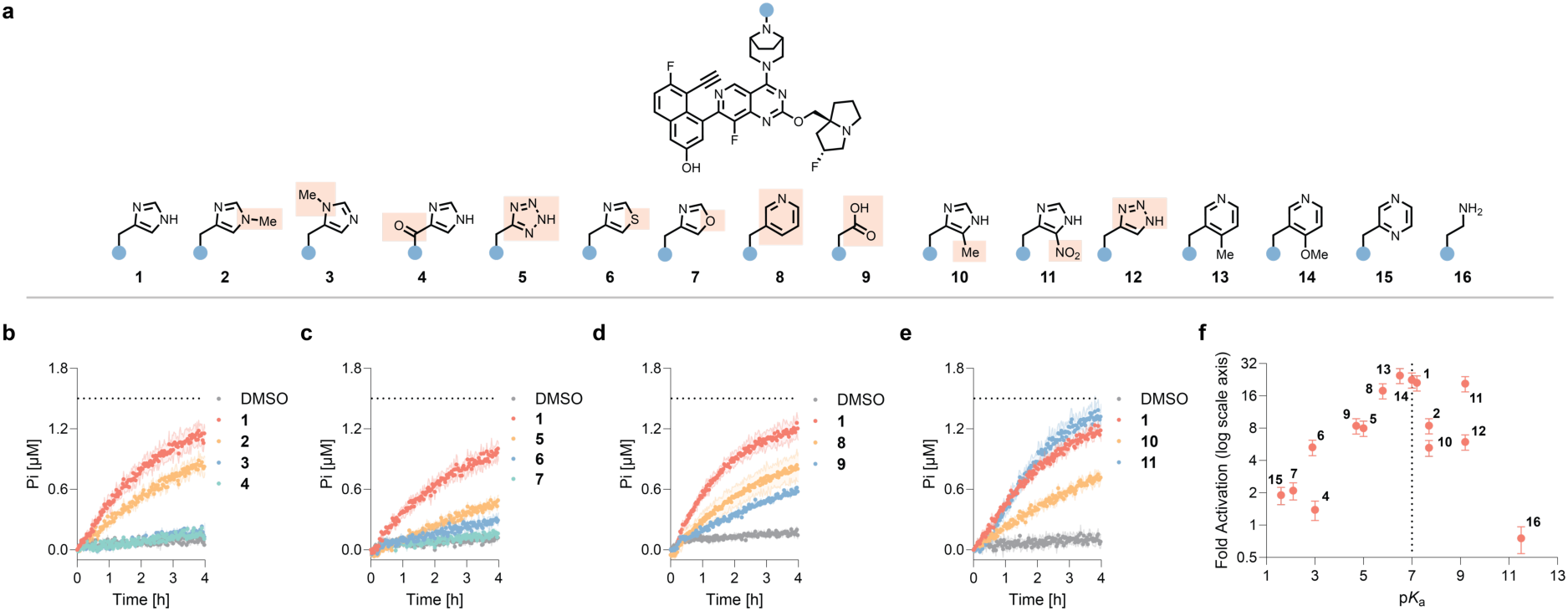
Activation of GTP hydrolysis requires a properly positioned Brøsted base with optimal pKa. **a.** Chemical structures of derivatives of compound **1**. **b-e.** GTP hydrolysis of K-Ras(Q61R)•GTP treated with compounds (50 µM) shown above. **f**. Relationship between fold activation (calculated as the increase in the apparent first-order kinetic constant, see Methods for details) of compound-mediated GTP hydrolysis and the calculated p*K*_a_ of the pendant group. **b**-**e**, Data is representative of three experiments and shown as mean ± SD of three technical replicates. **f**, Data is representative of three experiments and shown as mean ± SEM of three technical replicates.

We next asked whether any of the compounds were active against other K-Ras Q61 hotspot mutants. Compound **1** accelerated the GTP hydrolysis of all four Q61 mutants examined (Q61R, Q61L, Q61K, Q61H) in a concentration-dependent manner but did not significantly alter the intrinsic hydrolysis rate of wildtype or G12D (Fig. 4a-4b, Supplemental Fig. 2 and Supplemental Tables S2-S3). The lack of activity against wildtype K-Ras was not because the compounds fail to bind, as we were able to detect binding at 10 µM using a fluorescence polarization assay (Extended Data Fig. 5 and Supplemental Tables S4-S6). This finding suggests that small molecule-induced acceleration of GTP hydrolysis operates on when the native hydrolysis machinery is lost. Compound **1** did not rescue the hydrolysis of H-Ras(Q61R) or N-Ras(Q61R) (Figure 4D), as this Switch II ligand scaffold is known to be selective for K-Ras due to a critical hydrogen bonding interaction with His95, which is absent in H- and N-Ras^67,68^ (Fig. 4c). However, when we mutated Q95 (H-Ras) or L95 (N-Ras) to a histidine to confer compound sensitivity, the resulting proteins showed markedly accelerated GTP hydrolysis upon compound **1** treatment (Fig. 4d). These results demonstrate that while compound **1** does not directly engage H- and N-Ras in their native form, the chemical mechanism to promote GTP hydrolysis is likely applicable to a variety of oncogenic Q61X mutants once suitable ligand scaffolds are identified.

**Fig. 4.**
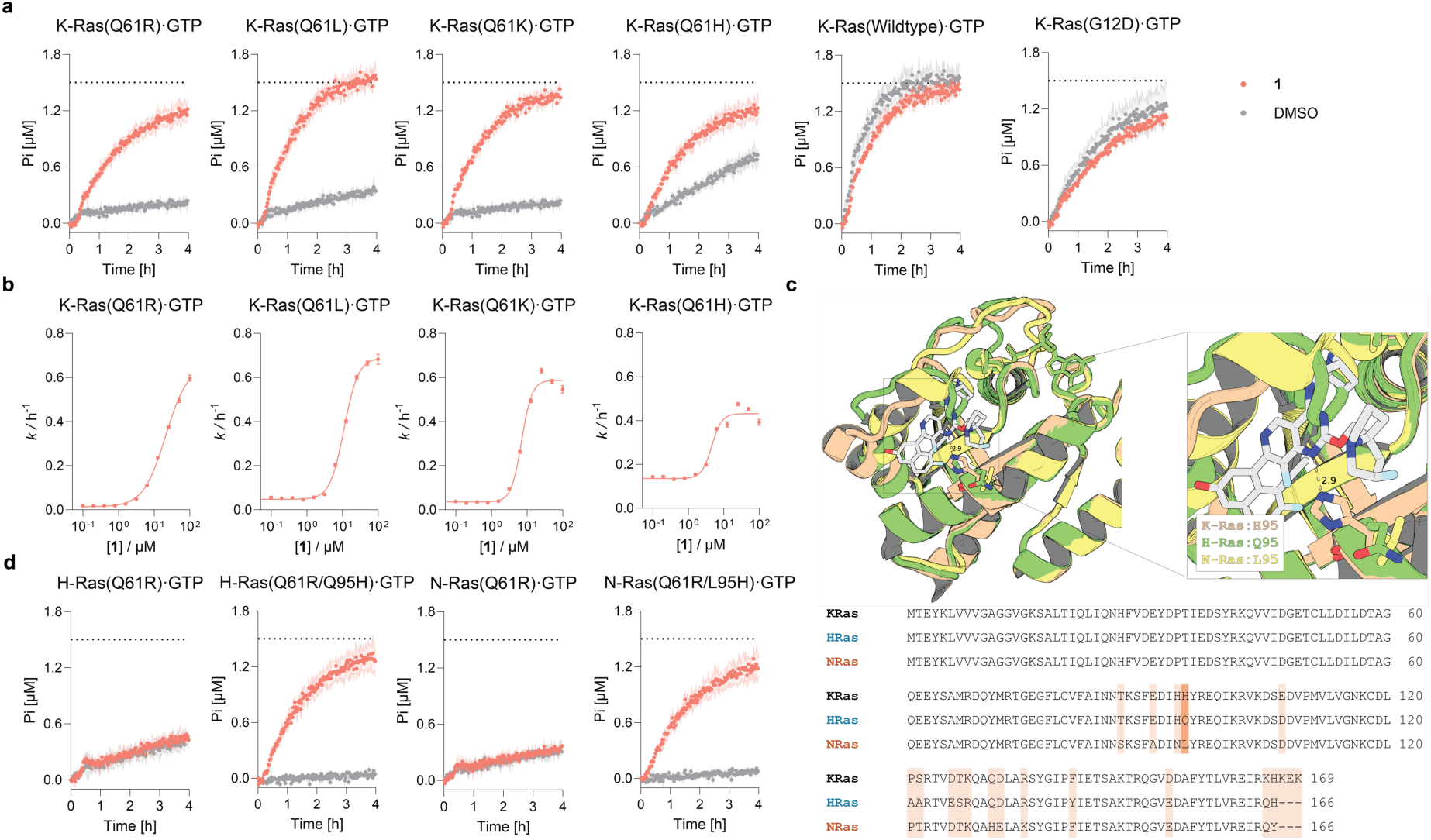
Compound 1 selectively accelerates GTP hydrolysis of Q61 mutants. **a**. Compound **1** accelerates the hydrolysis of K-Ras Q61 mutants, but not wildtype or G12D. **b**. Concentration-dependent acceleration of GTP hydrolysis for K-Ras Q61 mutants. **c**. Structures were obtained from K-Ras(G12D)•GppCp and MRTX1133 (PDB ID: 7T47), H-Ras(wildtype)•GppNHp (PDB ID: 5UHV), and N-Ras(wildtype)•GppNHp (PDB ID: 4EFL). The 95th residue is shown in stick representation, corresponding to H95 in K-Ras (wheat), Q95 in H-Ras (green), and L95 in N-Ras (yellow). Ligands and interacting residues are displayed for orientation. Sequence alignment of K-Ras, H-Ras and N-Ras, highlighting the sequence variation at histidine 95. **d.** Compound **1** rescue the hydrolysis of H-Ras and N-Ras mutants with histidine 95 substitutions. **a**, **b**, **d**, Data is representative of three experiments and shown as mean ± SD of three technical replicates.

We investigated whether our compounds allow targeted GTPase acceleration of K-Ras Q61X in genetically characterized cell lines. We chose to first evaluate our compounds in mouse embryonic fibroblasts in which the endogenous *KRAS, HRAS* and *NRAS* genes have been deleted and only a single driver oncogene is expressed (Rasless MEFs)^69^, as they provide a well-defined isogenic system to study a single Ras isoform. Compound **2**, which shows better solubility in cell culture media was tested on Rasless MEFs expressing K-Ras Q61R. We observed concentration- and time-dependent reduction of Ras•GTP levels in this cell line (Fig. 5a-5b), with the concomitant suppression of ERK phosphorylation (T202/Y204). At 4 h, compound **2** exhibited an apparent half-maximum inhibitory concentration (*IC*_50_) of 10 µM under these treatment conditions. We next examined a range of cell lines with characterized Ras Q61 mutations, including Rasless MEF K-Ras(Q61L), SW948 (K-Ras(Q61L/WT)), Calu-6 (K-Ras(Q61K/WT)), and NCI-H460 (K-Ras(Q61H/Q61H)). In each of these cell lines, treatment with 30 µM compound **2** for 4 h led to a reduction of Ras•GTP and p-ERK levels (Fig. 5c-5d). Meanwhile, compound **2** did not show any effect in Rasless MEF B-Raf(V600E) or A375 (K-Ras WT) cells, which both harbor B-Raf V600E mutations downstream of K-Ras (Fig. 5e). Together, these data indicate that compound **2** is active in cells and functions as a K-Ras(Q61X) accelerator by promoting GTP hydrolysis, thereby suppressing downstream ERK signaling in a Ras-dependent manner.

**Fig. 5.**
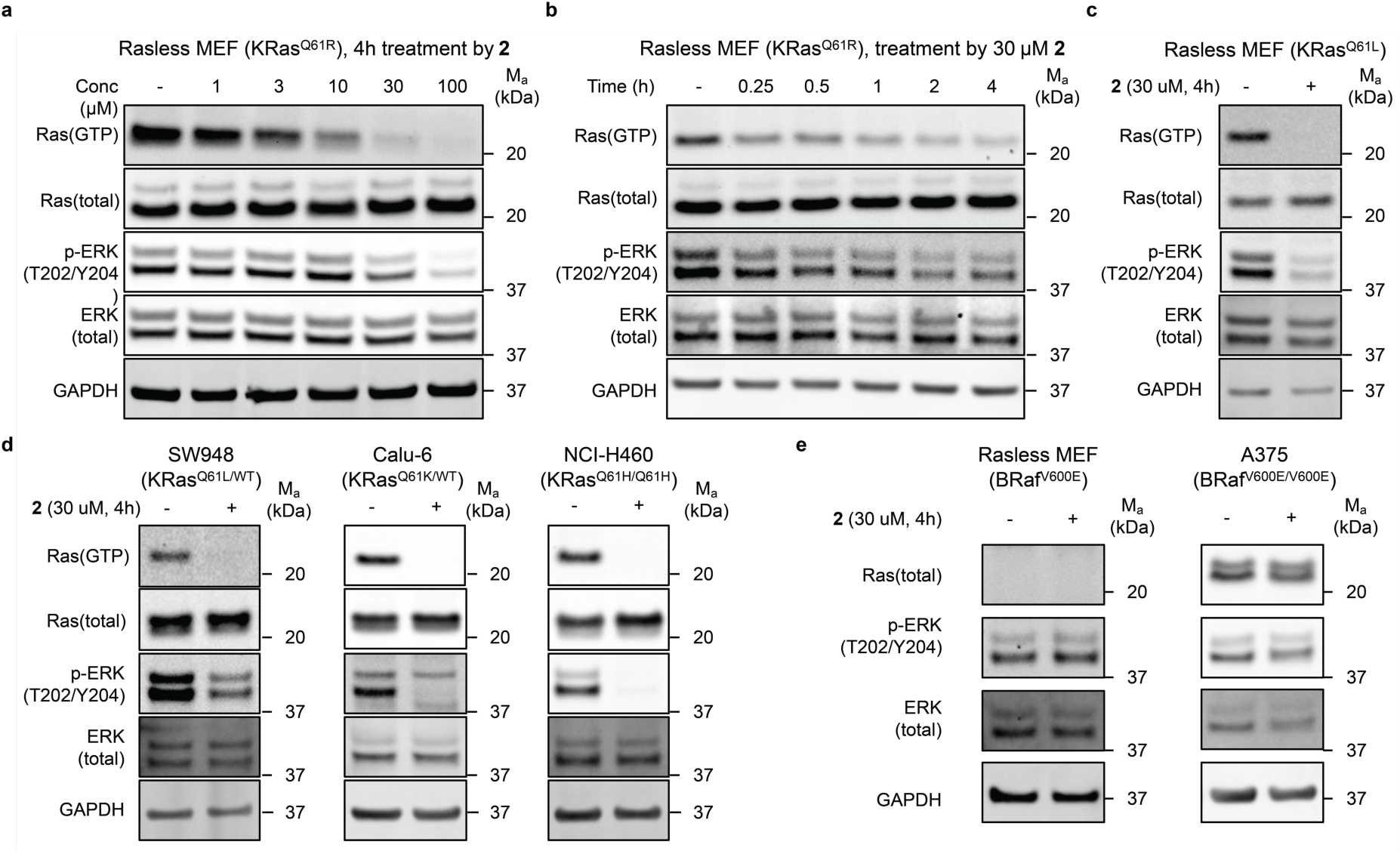
GTPase accelerators suppress MAPK signaling in K-Ras Q61 mutant cells. **a**. Rasless MEF K-Ras(Q61R) cells were treated with compound **2** at the indicated concentrations (0, 1, 3, 10, 30, and 100 μM) for 4 h. **b**. Rasless MEF K-Ras(Q61R) cells were treated with 30 μM compound **2** for the indicated times (0, 0.25, 0.5, 1, 2, and 4 h). **c**. Rasless MEF K-Ras(Q61L) cells were treated with 30 μM compound **2** for 4 h. **d**. Cancer cell lines harboring KRAS mutations were treated with 30 μM compound **2** for 4 h: SW948 K-Ras(Q61L/WT), Calu-6 K-Ras(Q61K/WT), NCI-H460 K-Ras(Q61H/Q61H). **e**. Cancer cell lines harboring BRAF mutations were treated with 30 μM compound **2** for 4 h: Rasless MEF B-Raf(V600E) cells and A375 B-Raf(V600E/V600E). Ras•GTP was detected by RAF-RBD pull-down and immunoblotting with anti-Ras antibody. Phosphorylated ERK (T202/Y204), total ERK, and GAPDH were analyzed by immunoblotting as indicated.

## Discussion

While K-Ras G12C inhibitors have demonstrated that direct targeting of mutant Ras can yield meaningful clinical benefit, no approved therapies yet exist for other oncogenic *RAS* mutants. Most *RAS* hotspot mutations, particularly those at G12 and Q61, disrupt both the intrinsic and GAP-mediated hydrolysis of GTP, resulting in persistent accumulation of the active, GTP-bound Ras that drives mitogenic signaling. This mechanistic insight suggests an alternative therapeutic strategy: rather than blocking Ras signaling, restoring GTP hydrolysis could inactivate mutant Ras while sparing the wild-type protein. Such a strategy offers broadly applicable route to target multiple RAS mutants with a mechanism-driven therapeutic index.

The ability of imidazole to act as a general acid/base (sometimes referred to as a “proton shuttle”) is crucial in many enzymatic and non-enzymatic mechanisms^70–74^. Imidazole, both as a free small molecule or as part of a substrate, has been shown to rescue the activity of catalytically-dead mutant kinases^57^ and proteases^56^. A method to introduce the imidazole moiety to the proximity of phosphodiester bonds using click chemistry has been developed to allow targeted cleavage of RNA, a mechanism that is partially dependent on the coordination of metal ions^75^. However, to the best of our knowledge, no chemical approaches to directly stimulate nucleotide hydrolysis have been reported. Our work demonstrates that stimulated nucleotide hydrolysis can be achieved by the precise delivery of a general base, such as imidazole, pyridine, and carboxylate groups, to allow the deprotonation of a water molecule and subsequent nucleophilic attack on the γ-phosphate of GTP.

Recently, Ras inhibitors that promote the formation of ternary complexes with cyclophilin A were found to accelerate GTP hydrolysis of G12 mutants, but not Q61 mutants, by stabilizing the transition state of intrinsic GTP hydrolysis^64,76^. Our work complements this approach by directly supplying a Brønsted base to rescue the broken catalytic machinery due to the loss of Gln61. Although we focused on K-Ras(Q61X) mutants in this study, we show that H-Ras and N-Ras mutants are similarly amenable to GTP hydrolysis acceleration using chemical genetics methods.

One limitation of the current work is that a high concentration of ligands is required for effective GTP hydrolysis. This is because currently available Switch II ligands have low affinity for the GTP-bound state and prefer the GDP-bound state of Ras^50^. We anticipate such limitations can be overcome with advances in ligand discovery^77–79^ and optimization. Overall, our findings lay a conceptual and mechanistic foundation for the development of small molecule GTPase accelerators for Ras-driven cancer. We are optimistic that our design principles may have broader utility for other small GTPases where defective GTP hydrolysis underlies disease mechanism.

## Acknowledgement

We thank the Pines Magnetic Resonance Center’s Core NMR Facility (PMRC Core) for spectroscopic assistance. The instrument used in this work was in part supported by NIH S10OD024998. This work is supported by the Damon Runyon Cancer Research Foundation (Damon Runyon–Rachleff Innovator program) and the Pew Charitable Trusts (Pew–Stewart Scholar program).

## Author contributions

Z.Z. and Y.-C.W. conceived the project, designed the study and wrote the manuscript. Y.-C.W., S.-C.C. and Z. S. synthesized the chemical ligands. Y.-C.W., Y.C., Y.W., Z.S., M.A.N and C.D.W. expressed the proteins and performed biochemical assays. S.-C.C. and Y.-C.W. performed the cellular assays. Y.-C.W. and H.C. acquired ^31^P NMR data. Y.-C.W., S.-C.C., Y.C., Y.W., Z.S., M.A.N., and C.D.W. performed analysis. All authors edited and approved the manuscript.

## Competing interests

Z.Z. and Y.-C.W. are inventors of a patent application related to this work owned by the University of California. Z.Z. receives stock and/or cash compensation from X-biotix Inc. and Montara Therapeutics.

## Data availability

Numerical data underlying the main and Supplementary Figures are included in the Source Data files. Uncropped western blots are included in the Source Data files. All data that support the findings of this study are available from the corresponding author upon reasonable request.

## Methods

### Recombinant protein expression and purification

*K-Ras (wild-type), K-Ras (G12D), K-Ras (Q61R), K-Ras(Q61L), K-Ras(Q61K), K-Ras(Q61H), H-Ras(Q61R), H-Ras(Q61R/Q95H), N-Ras(Q61R), N-Ras(Q61R/L95H), NF1-GRD, CypA, GST-RBD.* DNA sequences encoding human K-Ras (wild-type, amino acids 1–169), human K-Ras (G12D, amino acids 1–169), human K-Ras Q61R (amino acids 1–169), human K-Ras Q61L (amino acids 1–169), human K-Ras Q61K (amino acids 1–169)), human H-Ras Q61R (amino acids 1–166), human N-Ras Q61R (amino acids 1–166), human NF1-GRD (amino acids 1,203– 1,530) were codon optimized, synthesized by Twist Biosciences and cloned into pJExpress411 vector or noted below using the Gibson assembly method. Human K-Ras Q61H (amino acids 1– 169), human H-Ras Q61R/Q95H (amino acids 1–166), human N-Ras Q61R/L95H (amino acids 1–166) were constructed from QuikChange site-directed mutagenesis using the primers listed below. The resulting constructs contain a N-terminal 6× His tag and a TEV cleavage site (ENLYFQG). The proteins were expressed and purified following previously reported protocols, summarized below^2^.

*K-Ras (wild-type), K-Ras (G12D), K-Ras (Q61R), K-Ras(Q61L), K-Ras(Q61K), K-Ras(Q61H), H-Ras(Q61R), H-Ras(Q61R/Q95H), N-Ras(Q61R), N-Ras(Q61R/L95H).* Briefly, chemically competent BL21(DE3) cells (Invitrogen #C600003, propagated in house and chemically treated with Zymo Mix & Go Transformation Kit #T3002) were transformed with the corresponding plasmid and grown on LB agar plates containing 50 µg ml^−1^ kanamycin. A single colony was used to inoculate a culture at 37 °C, 220 r.p.m. in terrific broth containing 50 µg ml^−1^ kanamycin. When the optical density at 600 nm reached 0.6, the culture temperature was reduced to 20 °C, and protein expression was induced by the addition of IPTG (GoldBio, #I2481C50) to 0.2 mM. After 16 h at 20 °C, cells were pelleted by centrifugation (6,500 *g*, 10 min) and lysed in lysis buffer (20 mM Tris 8.0, 500 mM NaCl, 5 mM imidazole) by sonication. The lysate was clarified by high-speed centrifugation (19,000 *g*, 25 min) and the supernatant was used in subsequent purification by immobilized metal-affinity chromatography. His-TEV tagged protein was captured with Co-TALON resin (Clontech, Takara Bio #635503; 2 ml slurry per liter of culture) at 4 °C for 1 h with constant end-to-end mixing. The loaded beads were then washed with lysis buffer (50 ml per liter of culture) and the protein was eluted with elution buffer (20 mM Tris 8.0, 300 mM NaCl, 300 mM imidazole). To this protein solution was added His-tagged TEV protease (0.15 mg TEV per mg of Ras protein) and GDP (Abcam #ab146529, 1 mg per mg of Ras protein), and the mixture was dialyzed against TEV cleavage buffer (20 mM Tris 8.0, 300 mM NaCl, 1 mM EDTA) at 4 °C using a 10K molecular weight cutoff dialysis cassette (Thermo Scientific #66810) until LC–MS analysis showed full cleavage (typically 16–24 h). MgCl_2_ was added to a final concentration of 5 mM, and the mixture was incubated with 1 ml of Ni-NTA (Thermo Scientific #88222) beads at 4 °C for 1 h to remove TEV protease, any residual His-tagged proteins and peptides. The protein solution was diluted 1:10 v/v with 20 mM Tris 8.0 and further purified using anion-exchange chromatography (HiTrapQ column, 5 mL, Cytiva #17115401 0–30%B over 20 C.V.) (Buffer A: 20 mM Tris, pH 8.0, 50 mM NaCl, Buffer B: 20 mM Tris, pH 8.0, 500 mM NaCl). The protein was concentrated using a 10K molecular weight cutoff centrifugal concentrator (Amicon-15, Millipore #UFC901024) to 20 mg ml^−1^ and purified by size-exclusion chromatography on a Superdex 75 10/300 GL column (GE Healthcare Life Sciences #29-1487-21). Fractions containing pure biotinylated Ras protein were pooled, concentrated to 20 mg ml^−1^ and stored at –78 °C. In our hands, this protocol gives a typical yield of 5–15 mg per liter of culture.

*NF1-GRD.* Protein was expressed and purified using the identical protocol as for the Ras proteins, except that TEV cleavage, the ion exchange and nucleotide loading steps were omitted.

*CypA.* DNA sequences encoding human cyclophilin A (CypA) were synthesized by Twist Biosciences and cloned into pET47b vector using standard molecular biology techniques. Chemically competent BL21(DE3) cells were transformed with the pET47b-CypA plasmid and grown on LB agar plates containing 50 µg ml^−1^ kanamycin. A single colony was used to inoculate a culture at 37 °C, 220 r.p.m. in terrific broth containing 50 µg ml^−1^ kanamycin. When the optical density reached 0.6, the culture temperature was reduced to 20 °C, and protein expression was induced by the addition of IPTG to 1 mM. After 16 h at 20 °C, cells were pelleted by centrifugation (6,500g, 10 min) and lysed in lysis buffer (50 mM Tris 7.5, 150 mM NaCl, 30 mM imidazole) with sonication. Cleared lysates were subjected to affinity purification using Ni-NTA column (Thermo Scientific #88222). His-tagged CypA was eluted in 20mM Tris 7.5, 150mM NaCl, 400mM imidazole. Eluted fractions were subjected to buffer exchange (50mM Tris 8.0, 150mM KCl, 1mM DTT) by Amicon centrifugal filter (10kDa MWCO, Millipore #UFC901024). To this protein solution was added His-tagged HRV 3C protease (1mg protease per 100mg protein of interest). After overnight incubation at 4°C, LC-MS and SDS-PAGE confirmed complete cleavage. The protein solution was incubated with Ni-NTA resin under 4°C for 1h and eluted to afford pure full length CypA (confirmed by SDS-PAGE and LC-MS).

*GST-Raf1-RBD.* The GST-tagged RBD domain of Raf1 (residues 1-149, GST-Raf1-RBD) was expressed and purified following a published protocol.^80,81^

#### GTP and GppNHp loading for phosphate sensor assay

For Ras•GTP preparation, the Ras was diluted in EDTA Buffer (25 mM HEPES 7.5, 150 mM NaCl, 5 mM EDTA, 1 mM DTT) to 50 µM supplemented with 5 mM GTP at 0 °C to 250 µM. After incubation for 2 h on ice, the protein solutions were exchanged into buffer (50 mM HEPES 7.3, 150 mM KCl) using a PD-10 column (Cytiva #17085101) following the manufacturer’s instructions. The mixture was concentrated using an Amicon-4 concentrator (10,000 MWCO, Millipore #UFC801024), aliquoted by liquid N_2_ and stored at –78 °C.

### ^31^P NMR sample preparation and data acquisition

For K-Ras(Q61R)•GppNHp preparation, the K-Ras(Q61R) after anion exchange column as described above was diluted in reaction buffer (5 mM HEPES, pH 7.5, 150 mM NaCl, 5 mM EDTA, 1 mM DTT) to a final concentration of 315 µM K-Ras(Q61R), 21.3 mg GppNHp (final concentration 7.78 mM) was added. EDTA (0.5 M, pH 8.0, 200 µL added to afford final 25 mM EDTA). The total volume was 5.00 mL. They were incubated at 4 °C for 2.0 hours. The reaction buffer was exchanged to Phosphatase Buffer (32 mM Tris, pH 8.0, 20 mM Ammonium Sulfate, 0.1 mM ZnCl_2_) using a PD-10 column (Cytiva #17085101) following the manufacturer’s instructions. Then, 100 U CIP was added to the protein solution along with 20 mg of GppNHp, which was incubated for 12h at 4 °C. MgCl_2_ was added to a final concentration of 30 mM and incubated with protein solution for 15min on ice. The mixture was concentrated using an Amicon-4 concentrator (10,000 MWCO) and purified by SEC column (Superdex 75 10/300 GL, buffer: 22.2 mM HEPES, pH 7.3, 167 mM NaCl, 2.2 mM MgCl_2_). The GppNHp loading ratio was checked by AE column test. Standard GDP, GTP, GppNHp and K-Ras(Q61R)•GppNHp were prepared in 20 mM Tris, pH 8.0 to 600 µL x 25 µM. EDTA was added to 1 mM. The mixture was heated at 95 °C for 10 min and the insoluble material was pelleted by centrifugation (21,000 x g, 5 min). The supernatant was analyzed by ion exchange chromatography (HiTrapQ, 1 mL, Cytiva, 0-60% B over 20 C.V.): Prepare HiTrap Q 1 mL column with pump wash, 5 C.V. of Buffer A-Buffer B-Buffer A, inject sample to sample loop and start run method, which washes unbound with 5 C.V. of Buffer A, load the sample loop with 2 mL Buffer A, and start the gradient elution. Our protocol gives pure K-Ras(Q61R)•GppNHp without detectable GTP and GDP impurity. 20 µL compound **1** (20 mM) or DMSO was mixed with 86.6 µL Ras NMR buffer (22.2 mM HEPES, pH 7.3, 167 mM NaCl, 2.2 mM MgCl_2_) and 50.0 µL D_2_O first. Then 343 µL K-Ras(Q61R)•GppNHp (582 µM) was added and gently mixed. The sample was transferred into a 5 mm NMR tube for ^31^P NMR collection. The final NMR sample contains 400 µM K-Ras(Q61R)•GppNHp (with or without 800 µM compound **1**, 4% DMSO) of 500 µL 10% D_2_O in Ras NMR buffer.

For K-Ras(Q61R)•GTP preparation, the K-Ras(Q61R) after AE column as described above were diluted in EDTA Buffer (25 mM HEPES 7.5, 150 mM NaCl, 5 mM EDTA, 1 mM DTT) supplemented with 5 mM GTP at 0 °C to 250 µM. For this experiment, 391 µL K-Ras(Q61R) was diluted by EDTA buffer 4.61 mL, 14.7 mg GTP was added. After incubation for 2 h on ice, the protein solutions were exchanged into Ras NMR Buffer (22.2 mM HEPES, pH 7.3, 167 mM NaCl, 2.2 mM MgCl_2_) using a PD-10 column (Cytiva #17085101) following the manufacturer’s instructions. The mixture was concentrated using an Amicon-4 concentrator (10,000 MWCO, Millipore #UFC801024) to about 700 µL and purified by SEC column (Superdex 75 10/300 GL #29-1487-21, buffer: 22.2 mM HEPES, pH 7.3, 167 mM NaCl, 2.2 mM MgCl_2_). After that, 1506 µM K-Ras(Q61R)•GTP were aliquoted into three 255-µL aliquots and freeze by liquid N_2_ and then stored at –80 °C. It took 3-4 h in total after GTP loading. 39.6 µL compound **1** (20 mM) or DMSO was mixed with 248 µL Ras NMR buffer (22.2 mM HEPES, pH 7.3, 167 mM NaCl, 2.2 mM MgCl_2_) and 58.0 µL D_2_O first. Then 255 µL K-Ras(Q61R)•GTP (1506 µM) was added and gently mixed. The sample was transferred into a normal 5 mm NMR tube for ^31^P NMR collection. Ras concentration was increased to 662 µM here for better sensitivity of ^31^P NMR data in a single-hour long acquisition, allowing us to monitor the reaction with sufficient temporal resolution. The DMSO ratio was increased to 6% due to the solubility issue of compound **1**. The total volume of 10% D_2_O in Ras NMR buffer is increased to 600 µL for better shimming. The final sample contains 662 µM K-Ras(Q61R)•GTP (with or without 1324 µM compound **1**, 6% DMSO) in 600 µL 10% D_2_O in Ras NMR buffer.

^31^P NMR data was acquired on a 600 MHz Bruker Avance III spectrometer (^1^H at 600.13 and ^31^P 242.94 MHz) equipped with broadband observe Prodigy Cryoprobe. The sample temperature was maintained at 298 K and the experiments were conducted under N_2_ environment. The ^31^P NMR data were acquired using TopSpin v. 3.6.2 and a single pulse experiment, a recycle delay (d1) of 5s, a flip angle of 30° an acquisition time (aq) of 85 ms, a decoupling scheme of WALTZ-16.

^31^P NMR data for the K-Ras(Q61R)•GppNHp sample in static condition and after treatment with or without compound **1** were each time averaged with 8192 scans resulting in 11 hour 43 minutes long experimental time that yielded sufficient signal to noise ratio to analyze the perturbations in chemical shifts (Fig. 2b). ^31^P NMR data to characterize K-Ras(Q61R)•GTP hydrolysis catalyzed by compound **1** was acquired using 696 scans resulting in an experimental time of 60 minutes. The data acquisition was started immediately after 10 min and it was repeated 12 times to monitor the reaction dynamics over a 12-hour long period.

#### GppNHp loading for fluorescence polarization assay

The K-Ras(Q61R) and K-Ras(Q61L) after AE column as described above was loaded with GppNHp using a similar protocol as ^31^P NMR sample preparation. Our protocol gives pure K-Ras(Q61R)•GppNHp and K-Ras(Q61L)•GppNHp without GTP and GDP impurity. The mixture was concentrated using an Amicon-4 concentrator (10,000 MWCO, Millipore #UFC801024), aliquoted by liquid N_2_ and stored at –78 °C.

#### Phosphate sensor assay^82^

1.5 μM GTP/GDP/GppNHp loaded Ras protein or free GTP, 3.0 μM phosphate sensor (Thermo Fisher Scientific, #PV4407) in phosphate sensor assay buffer (50 mM HEPES 7.3, 150 mM KCl, 1.5 mM MgCl_2_) was incubated at 37 °C and treated with different concentration compounds or blank (1% DMSO in total) in Low Volume 384-well black/clear flat bottom polystyrene Not treated microplate (Corning #3540). The total volume of solution is 10 μL for each condition and each condition was repeated three times. Negative control is the phosphate sensor assay buffer and positive control is 2 µM Phosphate (diluted from 50 mM KH_2_PO_4_ with 2 mM NaN_3_ (Thermos Fisher Scientific #E6646,) The plate reader parameter is Excitation 430, Emission 450, Interval 1 min 30 s, Run time 4 h. For Figure 1D, NF1 was used as 1μM. RMC7977 was used as 10 μM, CypA was used as 5 μM. The phosphate release was measured using a phosphate sensor assay as described above. The hydrolysis rate was fitted using data from 0–2 h, except for wildtype where data from 0–1 h was used (due to fast reaction), according to the first-order reaction equation:

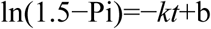

Where *k* is the apparent first-order kinetic constant. The half-life (*t*_₁/₂_) was calculated as (ln 2) / *k*.

#### Fluorescence polarization assay

Compound binding affinity was determined using a competition fluorescence polarization assay in phosphate sensor assay buffer. The *K*_d_ of the tracer molecules were first determined by measuring fluorescence polarization (excitation 485 nm, emission 535 nm) at various protein concentrations (40 µM–39 nM of K-Ras(Q61R)•GppNHp and 5 µM–4.9 nM of K-Ras(Q61L)•GppNHp) and fitting the curve to a quadratic binding model. To measure compound binding affinity, mixtures with the following composition were prepared in triplicate in low volume 384-well black plates (Corning #4511): 20 nM MRTX1133–FITC (**20**), 1000 nM K-Ras(Q61R)•GppNHp or 200 nM K-Ras(Q61L)•GppNHp, 1% DMSO, 100 µM–1.5 nM of test compound, 20 µL total volume. Data were fitted to a three-parameter sigmoidal curve to derive *IC*_50_ values. *K*_d_ of the compounds were calculated using a tool provided by Dr. Shaomeng Wang’s lab.

https://websites.umich.edu/~shaomengwanglab/software/calc_ki/index.html

#### p*K*_a_ prediction

For compounds **1**-**16** except compound **3**, which lost activity due to the change of N atom position, their predicted p*K*_a_ was calculated from a web server (https://xundrug.cn/molgpka) for small molecule p*K*_a_ prediction by graph-convolutional neural network.^66^ The structures for calculation were simplified as below with predicted p*K*_a_ and all structures were listed as base type.

#### Cell culture

Rasless MEF cells were obtained from Fredrick National Laboratory for Cancer Research and were maintained in DMEM (Gibco #11995073) supplemented with 10% FBS (Avantor #76419-584). SW948, H460 were obtained from Berkeley Cell Culture Facility and maintained in DMEM (Gibco #11995073) and 10% FBS (Avantor #76419-584). Calu-6, A375 human cancer cells were obtained from ATCC and maintained in DMEM (Gibco #11995073) and 10% FBS (Avantor #76419-584). Cells were passed for at least two generations after cryorecovery before they were used in the assays. When indicated, cells were treated with drugs at 40%–60% confluency at a final DMSO concentration of 0.3% except 1% for dose-dependent assay. At the end of the treatment period, cells were placed on ice. Unless otherwise indicated, adherent cells were washed once with ice-cold PBS (1 mL), scraped with a spatula and pelleted by centrifugation (500 *g*, 5 min). Cells were lysed in Co-IP lysis buffer (25 mM Tris, 150 mM NaCl, 5 mM MgCl_2_, 1% NP-40 and 5% glycerol) with Protease Inhibitor Cocktail (Roche #11836170001) and PhosSTOP (Roche #4906845001). Lysates were clarified by high-speed centrifugation (19,000 *g*, 10 min). Lysate concentrations were determined with a BCA protein assay (Thermo Fisher Scientific #A55862) and adjusted to 2 mg mL^−1^ with additional Co-IP lysis buffer. Samples were mixed with 5× SDS loading dye and heated at 95 °C for 5 min.

#### Analysis of GTP-bound Ras by Raf-RBD pulldown

Ras·GTP pulldown was performed with glutathione S-transferase-tagged Ras binding domain (GST-Raf1-RBD). Lysate (50 µL, 2 mg mL^−1^) was mixed with glutathione high-capacity magnetic agarose beads (Sigma, catalogue number G0924) and GST-RBD (30 µg), and the suspension was incubated at 4 °C for 1 h with constant end-to-end rotation. The beads were then washed with 2 × 500 µL of ice-cold Co-IP lysis buffer. Bound protein was eluted with 20 µL of 2× SDS loading buffer.

#### Gel electrophoresis and immunoblot

Unless otherwise noted, SDS–PAGE was run with SurePAGE 4–20% Bis-Tris gel (Genscript #M00654) in MES running buffer (Invitrogen) at 200 V for 20 min following the manufacturer’s instructions. Protein bands were transferred to 0.2-µm nitrocellulose membranes (Bio-Rad) using wet-tank transfer apparatus (Bio-Rad Criterion Blotter) in 1× TOWBIN buffer (25 mM Tris, 192 mM Glycine, pH 8.3) with 10% methanol at 75 V for 45 min. Membranes were blocked in 5% Bovine serum albumin (BSA) in Tris-buffered saline supplemented with Tween-20 (20 mM Tris, 150 mM NaCl, 0.1% Tween-20, pH 7.6, hereafter referred to as TBST) for 1 h at 23 °C. Primary antibody binding was performed with the indicated antibodies diluted in 5% BSA TBST at 4 °C for at least 16 h. After washing the membrane three times with TBST (5 min each wash), secondary antibodies (Li-COR IRDye 800CW goat anti-rabbit IgG, 1:10,000; and Li-COR IRDye 680RD goat anti-mouse IgG, 1:10,000) were added as solutions in 5% BSA–TBST at the dilutions recommended by the manufacturer. Secondary antibody binding was allowed to proceed for 1 h at 23 °C. The membrane was washed three times with TBST (5 min each wash) and imaged on a Bio-Rad ChemiDoc MP Imaging System.

#### Chemical synthesis

See Supplementary Information for details on chemical synthesis and ^1^H, ^19^F, and ^13^C NMR spectra.

---

**Extended Data Fig.1.**
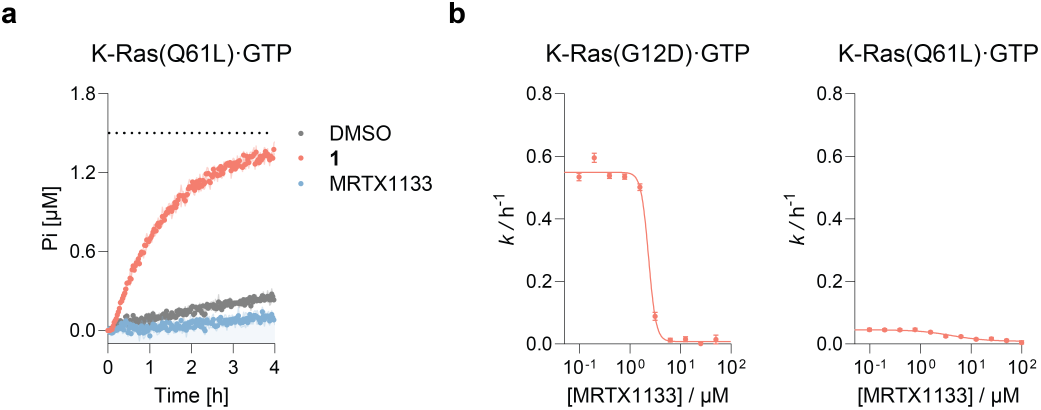
MRTX1133 inhibits GTP hydrolysis of K-Ras G12D and Q61L mutants. **A.** Compound MRTX1133 suppress the hydrolysis of K-Ras Q61L mutants. **B**. Dose-dependent inhibition of GTP hydrolysis for K-Ras G12D and Q61L mutants by MRTX1133. Data is representative of three experiments and shown as mean ± SD of three technical replicates.

**Extended Data Fig. 2.**
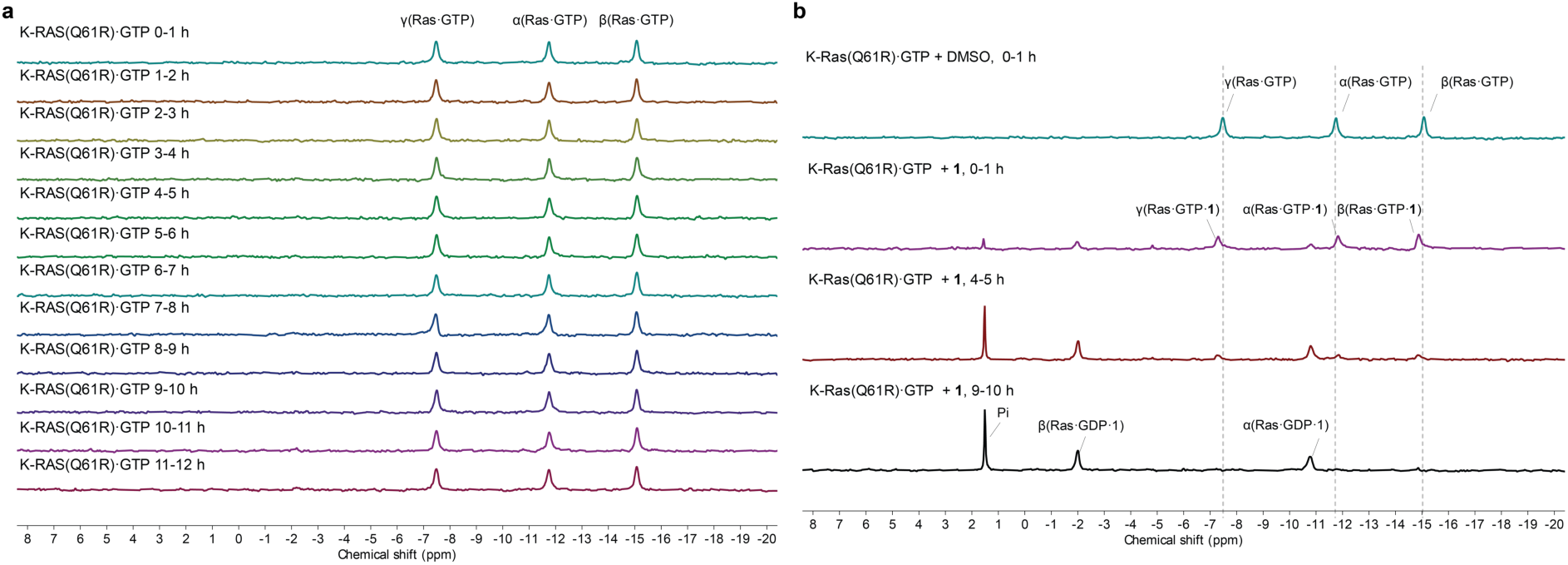
Time-dependent ^31^P NMR spectra monitoring 662 µM K-Ras(Q61R)•GTP hydrolysis in 6% DMSO. **a**. K-Ras(Q61R)·GTP recorded every hour showing slow intrinsic hydrolysis. **b**. Comparison of spectra at representative time points for K-Ras(Q61R)·GTP treated with DMSO or compound **1**.

**Extended Data Fig. 3.**
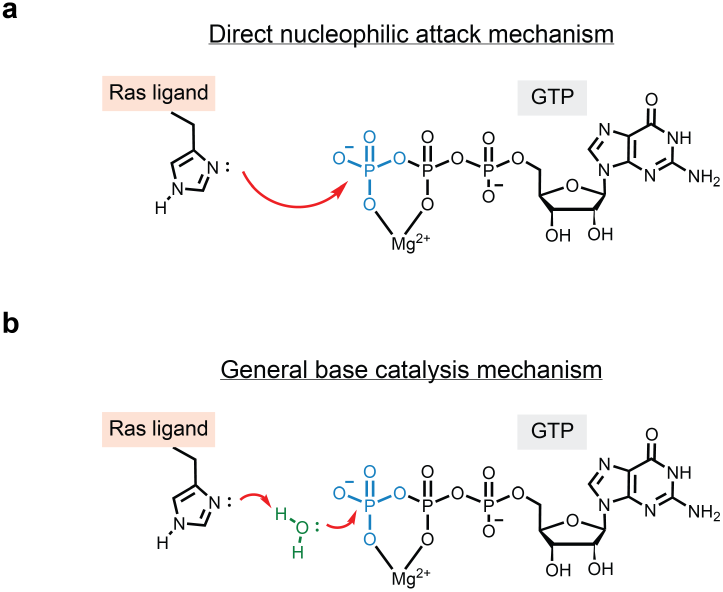
Proposed mechanisms of imidazole-catalyzed GTP hydrolysis. A. direct nucleophilic attack pathway, **B.** general base catalysis pathway.

**Extended Data Fig. 4.**
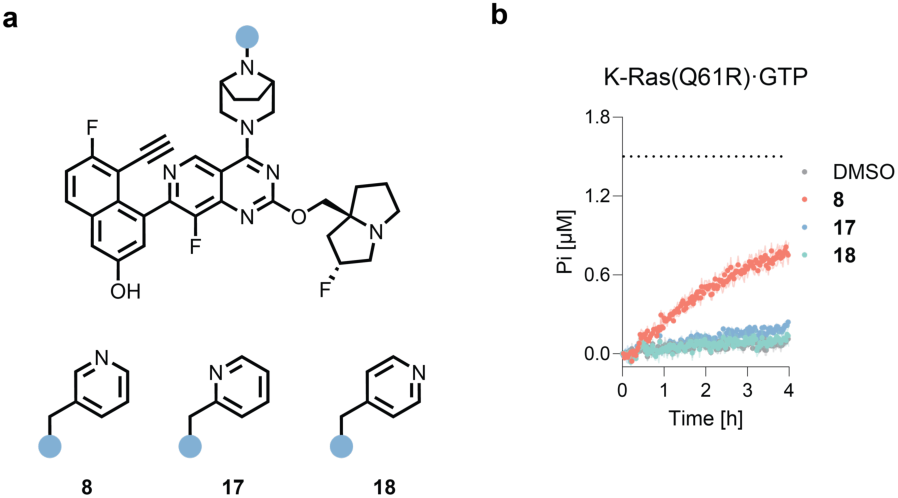
Effect of pyridine-containing derivatives on GTP hydrolysis of K-Ras(Q61R). **A.** Chemical structures of pyridine-containing compounds **8**, **17**, **18**. **B.** GTP hydrolysis of K-Ras(Q61R)•GTP treated with pyridine derivatives. Data is representative of three experiments and shown as mean ± SD of three technical replicates.

**Extended Data Fig. 5.**
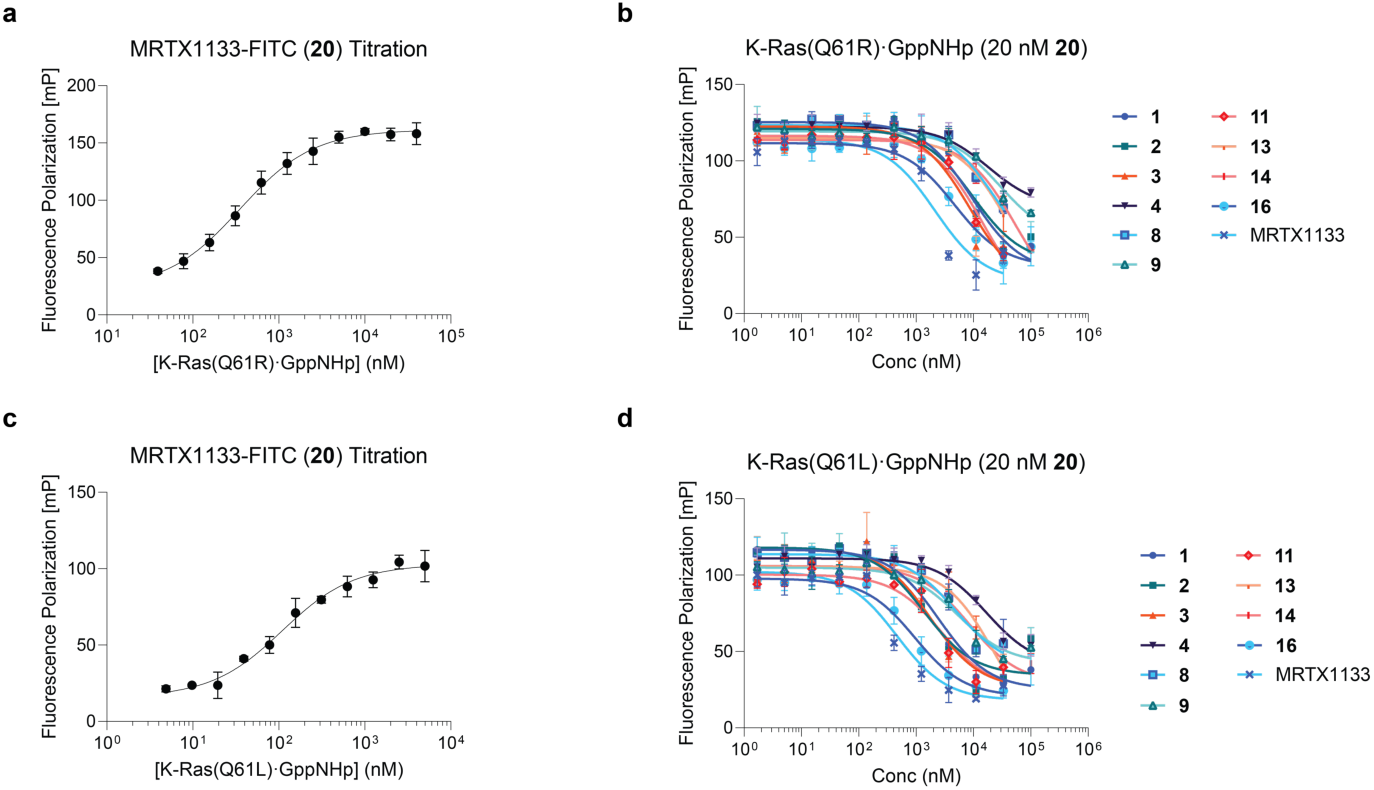
Representative fluorescence polarization (FP) binding and competition assays used for *K*_d_ determination between K-Ras(Q61R/L)•GppNHp and representative compounds. **a.** Titration curve of 20 nM MRTX1133-FITC (**20**) binding to K-Ras(Q61R)•GppNHp. **b.** FP competition assay showing displacement of 20 nM MRTX1133-FITC (**20**) by representative compounds in the presence of 1000 nM K-Ras(Q61R)•GppNHp. **c.** Titration curve of 20 nM MRTX1133-FITC (**20**) binding to K-Ras(Q61L)•GppNHp. **d.** FP competition assay showing displacement of 20 nM MRTX1133-FITC (**20**) by representative compounds in the presence of 200 nM K-Ras(Q61L)•GppNHp. Data is representative of three experiments and shown as mean ± SD of three technical replicates.

## References

(1) Prior, I. A.; Hood, F. E.; Hartley, J. L. The Frequency of Ras Mutations in Cancer. Cancer Res. 2020, 80 (14), 2969–2974. 10.1158/0008-5472.CAN-19-3682.

(2) Ostrem, J. M.; Peters, U.; Sos, M. L.; Wells, J. A.; Shokat, K. M. K-Ras(G12C) Inhibitors Allosterically Control GTP Affinity and Effector Interactions. Nature 2013, 503 (7477), 548–551. 10.1038/nature12796.

(3) Lanman, B. A.; Allen, J. R.; Allen, J. G.; Amegadzie, A. K.; Ashton, K. S.; Booker, S. K.; Chen, J. J.; Chen, N.; Frohn, M. J.; Goodman, G.; Kopecky, D. J.; Liu, L.; Lopez, P.; Low, J. D.; Ma, V.; Minatti, A. E.; Nguyen, T. T.; Nishimura, N.; Pickrell, A. J.; Reed, A. B.; Shin, Y.; Siegmund, A. C.; Tamayo, N. A.; Tegley, C. M.; Walton, M. C.; Wang, H. L.; Wurz, R. P.; Xue, M.; Yang, K. C.; Achanta, P.; Bartberger, M. D.; Canon, J.; Hollis, L. S.; McCarter, J. D.; Mohr, C.; Rex, K.; Saiki, A. Y.; San Miguel, T.; Volak, L. P.; Wang, K. H.; Whittington, D. A.; Zech, S. G.; Lipford, J. R.; Cee, V. J. Discovery of a Covalent Inhibitor of KRASG12C (AMG 510) for the Treatment of Solid Tumors. J. Med. Chem. 2020, 63 (1), 52–65. 10.1021/acs.jmedchem.9b01180.

(4) Hong, D. S.; Fakih, M. G.; Strickler, J. H.; Desai, J.; Durm, G. A.; Shapiro, G. I.; Falchook, G. S.; Price, T. J.; Sacher, A.; Denlinger, C. S.; Bang, Y.-J.; Dy, G. K.; Krauss, J. C.; Kuboki, Y.; Kuo, J. C.; Coveler, A. L.; Park, K.; Kim, T. W.; Barlesi, F.; Munster, P. N.; Ramalingam, S. S.; Burns, T. F.; Meric-Bernstam, F.; Henary, H.; Ngang, J.; Ngarmchamnanrith, G.; Kim, J.; Houk, B. E.; Canon, J.; Lipford, J. R.; Friberg, G.; Lito, P.; Govindan, R.; Li, B. T. KRAS G12C Inhibition with Sotorasib in Advanced Solid Tumors. N. Engl. J. Med. 2020, 383 (13), 1207–1217. 10.1056/nejmoa1917239.

(5) Fell, J. B.; Fischer, J. P.; Baer, B. R.; Blake, J. F.; Bouhana, K.; Briere, D. M.; Brown, K. D.; Burgess, L. E.; Burns, A. C.; Burkard, M. R.; Chiang, H.; Chicarelli, M. J.; Cook, A. W.; Gaudino, J. J.; Hallin, J.; Hanson, L.; Hartley, D. P.; Hicken, E. J.; Hingorani, G. P.; Hinklin, R. J.; Mejia, M. J.; Olson, P.; Otten, J. N.; Rhodes, S. P.; Rodriguez, M. E.; Savechenkov, P.; Smith, D. J.; Sudhakar, N.; Sullivan, F. X.; Tang, T. P.; Vigers, G. P.; Wollenberg, L.; Christensen, J. G.; Marx, M. A. Identification of the Clinical Development Candidate MRTX849, a Covalent KRASG12CInhibitor for the Treatment of Cancer. J. Med. Chem. 2020, 63 (13), 6679–6693. 10.1021/acs.jmedchem.9b02052.

(6) Schulze, C. J.; Seamon, K. J.; Zhao, Y.; Yang, Y. C.; Cregg, J.; Kim, D.; Tomlinson, A.; Choy, T. J.; Wang, Z.; Sang, B.; Pourfarjam, Y.; Lucas, J.; Cuevas-Navarro, A.; Ayala-Santos, C.; Vides, A.; Li, C.; Marquez, A.; Zhong, M.; Vemulapalli, V.; Weller, C.; Gould, A.; Whalen, D. M.; Salvador, A.; Milin, A.; Saldajeno-Concar, M.; Dinglasan, N.; Chen, A.; Evans, J.; Knox, J. E.; Koltun, E. S.; Singh, M.; Nichols, R.; Wildes, D.; Gill, A. L.; Smith, J. A. M.; Lito, P. Chemical Remodeling of a Cellular Chaperone to Target the Active State of Mutant KRAS. Science 2023, 381 (6659), 794–799. 10.1126/science.adg9652.

(7) Patricelli, M. P.; Janes, M. R.; Li, L.-S.; Hansen, R.; Peters, U.; Kessler, L. V.; Chen, Y.; Kucharski, J. M.; Feng, J.; Ely, T.; Chen, J. H.; Firdaus, S. J.; Babbar, A.; Ren, P.; Liu, Y. Selective Inhibition of Oncogenic KRAS Output with Small Molecules Targeting the Inactive State. Cancer Discov. 2016, 6 (3), 316–329. 10.1158/2159-8290.CD-15-1105.

(8) Lito, P.; Solomon, M.; Li, L.-S.; Hansen, R.; Rosen, N. Allele-Specific Inhibitors Inactivate Mutant KRAS G12C by a Trapping Mechanism. Science 2016, 351 (6273), 604–608. 10.1126/science.aad6204.

(9) Janes, M. R.; Zhang, J.; Li, L.-S.; Hansen, R.; Peters, U.; Guo, X.; Chen, Y.; Babbar, A.; Firdaus, S. J.; Darjania, L.; Feng, J.; Chen, J. H.; Li, S.; Li, S.; Long, Y. O.; Thach, C.; Liu, Y.; Zarieh, A.; Ely, T.; Kucharski, J. M.; Kessler, L. V.; Wu, T.; Yu, K.; Wang, Y.; Yao, Y.; Deng, X.; Zarrinkar, P. P.; Brehmer, D.; Dhanak, D.; Lorenzi, M. V.; Hu-Lowe, D.; Patricelli, M. P.; Ren, P.; Liu, Y. Targeting KRAS Mutant Cancers with a Covalent G12C-Specific Inhibitor. Cell 2018, 172 (3), 578–589.e17. 10.1016/j.cell.2018.01.006.

(10) Peng, S.-B.; Si, C.; Zhang, Y.; Van Horn, R. D.; Lin, X.; Gong, X.; Huber, L.; Donoho, G.; Curtis, C.; Strelow, J. M.; Bocchinfuso, W. P.; Guo, D.; Boulet, S. L.; Barda, D.; Manglicmot, D.; Saflor, M.-B. D.; Wang, J.; Xiao, J.; Chalmers, M. J.; Burns, L.; Linder, R. J.; Ackermann, B. L.; Cornwell, P. D.; Zhou, L.; McCann, D.; Henry, J. Abstract 1259: Preclinical Characterization of LY3537982, a Novel, Highly Selective and Potent KRAS-G12C Inhibitor. Cancer Res. 2021, 81 (13_Supplement), 1259–1259. 10.1158/1538-7445.AM2021-1259.

(11) Savarese, F.; Gollner, A.; Rudolph, D.; Lipp, J.; Popow, J.; Hofmann, M. H.; Arnhof, H.; Rinnenthal, J.; Trapani, F.; Gmachl, M.; Gerlach, D.; Broeker, J.; Ettmayer, P.; Mantoulidis, A.; Phan, J.; Smethurst, C. A.; Treu, M.; Waterson, A. G.; Lu, H.; Machado, A.; Daniele, J.; Fesik, S. W.; Vellano, C. P.; Heffernan, T. P.; Marszalek, J. R.; McConnell, D. B.; Petronczki, M.; Kraut, N.; Waizenegger, I. C. Abstract 1271: *In Vitro* and *in Vivo* Characterization of BI 1823911 -a Novel KRASG12C Selective Small Molecule Inhibitor. Cancer Res. 2021, 81 (13_Supplement), 1271–1271. 10.1158/1538-7445.AM2021-1271.

(12) Li, Z.; Song, Z.; Zhao, Y.; Wang, P.; Jiang, L.; Gong, Y.; Zhou, J.; Jian, H.; Dong, X.; Zhuang, W.; Cang, S.; Yang, N.; Fang, J.; Shi, J.; Lu, J.; Ma, R.; Wu, P.; Zhang, Y.; Song, M.; Xu, C.-W.; Shi, Z.; Zhang, L.; Wang, Y.; Wang, X.; Zhang, Y.; Lu, S. D-1553 (Garsorasib), a Potent and Selective Inhibitor of KRASG12C in Patients With NSCLC: Phase 1 Study Results. J. Thorac. Oncol. 2023, 18 (7), 940–951. 10.1016/j.jtho.2023.03.015.

(13) Weiss, A.; Lorthiois, E.; Barys, L.; Beyer, K. S.; Bomio-Confaglia, C.; Burks, H.; Chen, X.; Cui, X.; De Kanter, R.; Dharmarajan, L.; Fedele, C.; Gerspacher, M.; Guthy, D. A.; Head, V.; Jaeger, A.; Núñez, E. J.; Kearns, J. D.; Leblanc, C.; Maira, S.-M.; Murphy, J.; Oakman, H.; Ostermann, N.; Ottl, J.; Rigollier, P.; Roman, D.; Schnell, C.; Sedrani, R.; Shimizu, T.; Stringer, R.; Vaupel, A.; Voshol, H.; Wessels, P.; Widmer, T.; Wilcken, R.; Xu, K.; Zecri, F.; Farago, A. F.; Cotesta, S.; Brachmann, S. M. Discovery, Preclinical Characterization, and Early Clinical Activity of JDQ443, a Structurally Novel, Potent, and Selective Covalent Oral Inhibitor of KRASG12C. Cancer Discov. 2022, 12 (6), 1500–1517. 10.1158/2159-8290.CD-22-0158.

(14) Zhou, Q.; Yang, N.; Zhao, M.; Huang, D.; Zhao, J.; Yu, Y.; Yuan, Y.; Sun, L.; Dong, X.; Zhang, T.; Chu, Q.; Li, X.; Meng, X.; Wang, H.; Wang, X.; Wu, D.; Hu, S.; Shan, J.; Liu, L.; Sun, M.; Zhang, Z.; Zhu, H.; Huang, J.; Huang, M.; Cheng, L.; Zhang, S.; Zhou, H.; Wu, Y.-L. Potent Covalent Irreversible Inhibitor of KRAS G12C IBI351 in Patients with Advanced Solid Tumors: First-in-Human Phase I Study. Eur. J. Cancer 2024, 212, 114337. 10.1016/j.ejca.2024.114337.

(15) Sacher, A.; LoRusso, P.; Patel, M. R.; Miller, W. H.; Garralda, E.; Forster, M. D.; Santoro, A.; Falcon, A.; Kim, T. W.; Paz-Ares, L.; Bowyer, S.; De Miguel, M.; Han, S.-W.; Krebs, M. G.; Lee, J.-S.; Cheng, M. L.; Arbour, K.; Massarelli, E.; Choi, Y.; Shi, Z.; Mandlekar, S.; Lin, M. T.; Royer-Joo, S.; Chang, J.; Dharia, N. V.; Schutzman, J. L.; Desai, J. Single-Agent Divarasib (GDC-6036) in Solid Tumors with a *KRAS* G12C Mutation. N. Engl. J. Med. 2023, 389 (8), 710–721. 10.1056/NEJMoa2303810.

(16) Kim, D.; Herdeis, L.; Rudolph, D.; Zhao, Y.; Böttcher, J.; Vides, A.; Ayala-Santos, C. I.; Pourfarjam, Y.; Cuevas-Navarro, A.; Xue, J. Y.; Mantoulidis, A.; Bröker, J.; Wunberg, T.; Schaaf, O.; Popow, J.; Wolkerstorfer, B.; Kropatsch, K. G.; Qu, R.; De Stanchina, E.; Sang, B.; Li, C.; McConnell, D. B.; Kraut, N.; Lito, P. Pan-KRAS Inhibitor Disables Oncogenic Signalling and Tumour Growth. Nature 2023, 619 (7968), 160–166. 10.1038/s41586-023-06123-3.

(17) Zhang, Z.; Morstein, J.; Ecker, A. K.; Guiley, K. Z.; Shokat, K. M. Chemoselective Covalent Modification of K-Ras(G12R) with a Small Molecule Electrophile. J. Am. Chem. Soc. 2022, 144 (35), 15916–15921. 10.1021/jacs.2c05377.

(18) Zhang, Z.; Guiley, K. Z.; Shokat, K. M. Chemical Acylation of an Acquired Serine Suppresses Oncogenic Signaling of K-Ras(G12S). Nat. Chem. Biol. 2022, 18 (11), 1177–1183. 10.1038/s41589-022-01065-9.

(19) Lanman, B. A.; Wurz, R. P.; Verma, R.; Osgood, T.; Gaida, K.; Mohn, D.; Chen, Y.-C.; Diaz, G.; Saiki, A. Y.; Hughes, P. E.; Mohr, C.; Vaish, A.; Tudor, Y.; Dahal, U. P.; Agarwal, P.; Mollica, C. Abstract ND01: AMG 410: An H/NRAS-Sparing Pan-KRAS Inhibitor with Dual GTP(on)/GDP(off)-State Activity for the Treatment of Diverse KRAS-Mutant Tumors. Cancer Res. 2025, 85 (8_Supplement_2), ND01–ND01. 10.1158/1538-7445.AM2025-ND01.

(20) Kessler, D.; Gmachl, M.; Mantoulidis, A.; Martin, L. J.; Zoephel, A.; Mayer, M.; Gollner, A.; Covini, D.; Fischer, S.; Gerstberger, T.; Gmaschitz, T.; Goodwin, C.; Greb, P.; Häring, D.; Hela, W.; Hoffmann, J.; Karolyi-Oezguer, J.; Knesl, P.; Kornigg, S.; Koegl, M.; Kousek, R.; Lamarre, L.; Moser, F.; Munico-Martinez, S.; Peinsipp, C.; Phan, J.; Rinnenthal, J.; Sai, J.; Salamon, C.; Scherbantin, Y.; Schipany, K.; Schnitzer, R.; Schrenk, A.; Sharps, B.; Siszler, G.; Sun, Q.; Waterson, A.; Wolkerstorfer, B.; Zeeb, M.; Pearson, M.; Fesik, S. W.; McConnell, D. B. Drugging an Undruggable Pocket on KRAS. Proc. Natl. Acad. Sci. 2019, 116 (32), 15823–15829. 10.1073/pnas.1904529116.

(21) Zhou, C.; Li, C.; Luo, L.; Li, X.; Jia, K.; He, N.; Mao, S.; Wang, W.; Shao, C.; Liu, X.; Huang, K.; Yu, Y.; Cai, X.; Chen, Y.; Dai, Z.; Li, W.; Yu, J.; Li, J.; Shen, F.; Wang, Z.; He, F.; Sun, X.; Mao, R.; Shi, W.; Zhang, J.; Jiang, T.; Zhang, Z.; Li, F.; Ren, S. Anti-Tumor Efficacy of HRS-4642 and Its Potential Combination with Proteasome Inhibition in KRAS G12D-Mutant Cancer. Cancer Cell 2024, 42 (7), 1286–1300.e8. 10.1016/j.ccell.2024.06.001.

(22) Zheng, Q.; Zhang, Z.; Guiley, K. Z.; Shokat, K. M. Strain-Release Alkylation of Asp12 Enables Mutant Selective Targeting of K-Ras-G12D. Nat. Chem. Biol. 2024, 20 (9), 1114–1122. 10.1038/s41589-024-01565-w.

(23) Sirocchi, L. S.; Scharnweber, M.; Oberndorfer, S.; Siszler, G.; Zak, K. M.; Rumpel, K.; Neumüller, R. A.; Wilding, B. Discovery of Carbodiimide Warheads to Selectively and Covalently Target Aspartic Acid in KRAS^G12D^. J. Am. Chem. Soc. 2025, 147 (18), 15787–15795. 10.1021/jacs.5c03562.

(24) Chen, Z.; Eriksson, A.; Lee, B.; Dinglasan, J.; Montazer, N.; Cregg, J.; Edwards, A.; Sanders, K.; Smith, J. A.; Wildes, D.; Singh, M.; Wang, Z.; Jiang, J. Abstract 3340: RMC-5127, a First-in-Class, Orally Bioavailable Mutant-Selective RASG12V(ON) Inhibitor Is Central Nervous System (CNS)-Penetrant and Demonstrates Anti-Tumor Activity in a Preclinical Intracranial Xenograft Model. Cancer Res. 2024, 84 (6_Supplement), 3340–3340. 10.1158/1538-7445.AM2024-3340.

(25) Bröker, J.; Waterson, A. G.; Smethurst, C.; Kessler, D.; Böttcher, J.; Mayer, M.; Gmaschitz, G.; Phan, J.; Little, A.; Abbott, J. R.; Sun, Q.; Gmachl, M.; Rudolph, D.; Arnhof, H.; Rumpel, K.; Savarese, F.; Gerstberger, T.; Mischerikow, N.; Treu, M.; Herdeis, L.; Wunberg, T.; Gollner, A.; Weinstabl, H.; Mantoulidis, A.; Krämer, O.; McConnell, D. B.; W. Fesik, S. Fragment Optimization of Reversible Binding to the Switch II Pocket on KRAS Leads to a Potent, In Vivo Active KRAS^G12C^ Inhibitor. J. Med. Chem. 2022, 65 (21), 14614–14629. 10.1021/acs.jmedchem.2c01120.

(26) Maurer, T.; Garrenton, L. S.; Oh, A.; Pitts, K.; Anderson, D. J.; Skelton, N. J.; Fauber, B. P.; Pan, B.; Malek, S.; Stokoe, D.; Ludlam, M. J. C.; Bowman, K. K.; Wu, J.; Giannetti, A. M.; Starovasnik, M. A.; Mellman, I.; Jackson, P. K.; Rudolph, J.; Wang, W.; Fang, G. Small-Molecule Ligands Bind to a Distinct Pocket in Ras and Inhibit SOS-Mediated Nucleotide Exchange Activity. Proc. Natl. Acad. Sci. 2012, 109 (14), 5299–5304. 10.1073/pnas.1116510109.

(27) Quevedo, C. E.; Cruz-Migoni, A.; Bery, N.; Miller, A.; Tanaka, T.; Petch, D.; Bataille, C. J. R.; Lee, L. Y. W.; Fallon, P. S.; Tulmin, H.; Ehebauer, M. T.; Fernandez-Fuentes, N.; Russell, A.J.; Carr, S. B.; Phillips, S. E. V.; Rabbitts, T. H. Small Molecule Inhibitors of RAS-Effector Protein Interactions Derived Using an Intracellular Antibody Fragment. Nat. Commun. 2018, 9 (1), 3169. 10.1038/s41467-018-05707-2.

(28) Bery, N.; Legg, S.; Debreczeni, J.; Breed, J.; Embrey, K.; Stubbs, C.; Kolasinska-Zwierz, P.; Barrett, N.; Marwood, R.; Watson, J.; Tart, J.; Overman, R.; Miller, A.; Phillips, C.; Minter, R.; Rabbitts, T. H. KRAS-Specific Inhibition Using a DARPin Binding to a Site in the Allosteric Lobe. Nat. Commun. 2019, 10 (1), 2607. 10.1038/s41467-019-10419-2.

(29) Rojas, C. I.; Lugowska, I.; Juergens, R.; Sacher, A.; Weindler, S.; Sendur, M. A. N.; Dziadziuszko, R.; Pal, A.; Alvarez, E. C.; Ahern, E. S.; Lakhani, N.; Chen, L.; Jemielita, T.; Choi, S. Y.; Stathis, A. 44O Updated Results from a Phase I Study Evaluating the KRAS G12C Inhibitor MK-1084 in Solid Tumors and in Combination with Pembrolizumab in NSCLC. ESMO Open 2024, 9, 102273. 10.1016/j.esmoop.2024.102273.

(30) Li, J.; Zhao, J.; Cao, B.; Fang, J.; Li, X.; Wang, M.; Ba, Y.; Li, X.; Li, Z.; Liu, Z.; Wang, Y.; Cheng, Y.; Bai, C.; Shen, L. A Phase I/II Study of First-in-Human Trial of JAB-21822 (KRAS G12C Inhibitor) in Advanced Solid Tumors. J. Clin. Oncol. 2022, 40 (16_suppl), 3089–3089. 10.1200/JCO.2022.40.16_suppl.3089.

(31) Zhou, Q.; Meng, X.; Sun, L.; Huang, D.; Yang, N.; Yu, Y.; Zhao, M.; Zhuang, W.; Hu, Y.; Pan, Y.; Sun, M.; Shan, J.; Guo, R.-H.; Chu, Q.; Xu, C.; Lin, J.; Huang, J.; Huang, M.; Shen, Y.; Wu, Y.-L. LBA12 Efficacy and Safety of IBI351 (GFH925), a Selective KRASG12C Inhibitor, Monotherapy in Patients (Pts) with Advanced Non-Small Cell Lung Cancer (NSCLC): Initial Results from a Registrational Phase II Study. Ann. Oncol. 2023, 34, S1662. 10.1016/j.annonc.2023.10.584.

(32) Waizenegger, I. C.; Lu, H.; Thamer, C.; Savarese, F.; Gerlach, D.; Rudolph, D.; Vellano, C. P.; Marotti, M.; Heymach, J.; Kopetz, S.; Heffernan, T. P.; Marszalek, J. R.; Petronczki, M. P.; Hofmann, M. H.; Kraut, N. Abstract 2667: Trial in Progress: Phase 1 Study of BI 1823911, an Irreversible KRASG12C Inhibitor Targeting KRAS in Its GDP-Loaded State, as Monotherapy and in Combination with the Pan-KRAS SOS1 Inhibitor BI 1701963 in Solid Tumors Expressing KRASG12C Mutation. Cancer Res. 2022, 82 (12_Supplement), 2667–2667. 10.1158/1538-7445.AM2022-2667.

(33) Wang, J.; Martin-Romano, P.; Cassier, P.; Johnson, M.; Haura, E.; Lenox, L.; Guo, Y.; Bandyopadhyay, N.; Russell, M.; Shearin, E.; Lauring, J.; Dahan, L. Phase I Study of JNJ-74699157 in Patients with Advanced Solid Tumors Harboring the *KRAS G12C* Mutation. The Oncologist 2022, 27 (7), 536–e553. 10.1093/oncolo/oyab080.

(34) Liu, R.; Qu, X.; Yang, N.; Chai, X.; Xu, J. First-in-Human Study of ZG19018, Targeting KRAS G12C, as Monotherapy in Patients with Advanced Solid Tumors. J. Clin. Oncol. 2023, 41 (16_suppl), e15127–e15127. 10.1200/JCO.2023.41.16_suppl.e15127.

(35) Bröker, J.; Waterson, A. G.; Hodges, T. R.; Abbott, J. R.; Arnold, A.; Böttcher, J.; Braun, N.; Cui, J.; Fuchs, J. E.; Gerstberger, T.; Gogg, S.; Hanner, S.; Herdeis, L.; Howell, L. W.; Mantoulidis, A.; Mayer, M.; Phan, J.; Rocchetti, F.; Sankar, K.; Sarkar, D.; Schaaf, O.; Sensintaffar, J. L.; Sun, Q.; Wunberg, T.; Fesik, S. W. Discovery of BI-2493, a Pan-KRAS Inhibitor Showing *In Vivo* Efficacy. J. Med. Chem. 2025, 68 (15), 15649–15668. 10.1021/acs.jmedchem.5c00576.

(36) Aladinskiy, V.; Mantsyzov, A. B.; Kruse, C.; Noev, A.; Petrov, R.; Reshetnikov, V.; Shi, S.; Ding, X.; Cai, X.; Aliper, A.; Zhavoronkov, A.; Ren, F. Identification of Novel Pan-KRAS Inhibitors via Structure-Based Drug Design, Scaffold Hopping, and Biological Evaluation. ACS Med. Chem. Lett. 2025, 16 (7), 1282–1289. 10.1021/acsmedchemlett.5c00080.

(37) Yu, Z.; He, X.; Wang, R.; Xu, X.; Zhang, Z.; Ding, K.; Zhang, Z.-M.; Tan, Y.; Li, Z. Simultaneous Covalent Modification of K-Ras(G12D) and K-Ras(G12C) with Tunable Oxirane Electrophiles. J. Am. Chem. Soc. 2023, 145 (37), 20403–20411. 10.1021/jacs.3c05899.

(38) Tanada, M.; Tamiya, M.; Matsuo, A.; Chiyoda, A.; Takano, K.; Ito, T.; Irie, M.; Kotake, T.; Takeyama, R.; Kawada, H.; Hayashi, R.; Ishikawa, S.; Nomura, K.; Furuichi, N.; Morita, Y.; Kage, M.; Hashimoto, S.; Nii, K.; Sase, H.; Ohara, K.; Ohta, A.; Kuramoto, S.; Nishimura, Y.; Iikura, H.; Shiraishi, T. Development of Orally Bioavailable Peptides Targeting an Intracellular Protein: From a Hit to a Clinical KRAS Inhibitor. J. Am. Chem. Soc. 2023, 145 (30), 16610–16620. 10.1021/jacs.3c03886.

(39) Ye, Q.; Shvartsbart, A.; Li, Z.; Gan, P.; Policarpo, R. L.; Qi, C.; Roach, J. J.; Zhu, W.; McCammant, M. S.; Hu, B.; Li, G.; Yin, H.; Carlsen, P.; Hoang, G.; Zhao, L.; Susick, R.; Zhang, F.; Lai, C.-T.; Allali Hassani, A.; Epling, L. B.; Gallion, A.; Kurzeja-Lipinski, K.; Gallagher, K.; Roman, V.; Farren, M. R.; Kong, W.; Deller, M. C.; Zhang, G.; Covington, M.; Diamond, S.; Kim, S.; Yao, W.; Sokolsky, A.; Wang, X. Discovery of INCB159020, an Orally Bioavailable KRAS G12D Inhibitor. J. Med. Chem. 2025, 68 (2), 1924–1939. 10.1021/acs.jmedchem.4c02662.

(40) Zheng, Q.; Shokat, K. M. Denitrogenative Alkylation of K-Ras(G12D) Inhibits Oncogenic Signaling in Cancer Cells. J. Am. Chem. Soc. 2025, 147 (28), 24785–24792. 10.1021/jacs.5c06745.

(41) Bond, M. J.; Chu, L.; Nalawansha, D. A.; Li, K.; Crews, C. M. Targeted Degradation of Oncogenic KRAS^G12C^ by VHL-Recruiting PROTACs. ACS Cent. Sci. 2020, 6 (8), 1367–1375. 10.1021/acscentsci.0c00411.

(42) Yoshinari, T.; Nagashima, T.; Ishioka, H.; Inamura, K.; Nishizono, Y.; Tasaki, M.; Iguchi, K.; Suzuki, A.; Sato, C.; Nakayama, A.; Amano, Y.; Tateishi, Y.; Yamanaka, Y.; Osaki, F.; Yoshino, M.; Kuramoto, K.; Imaizumi, T.; Hayakawa, M. Discovery of KRAS(G12D) Selective Degrader ASP3082. Commun. Chem. 2025, 8 (1), 254. 10.1038/s42004-025-01662-4.

(43) Moore, A. R.; Rosenberg, S. C.; McCormick, F.; Malek, S. RAS-Targeted Therapies: Is the Undruggable Drugged? Nat. Rev. Drug Discov. 2020, 19 (8), 533–552. 10.1038/s41573-020-0068-6.

(44) Der, C. J.; Finkel, T.; Cooper, G. M. Biological and Biochemical Properties of Human *Ras*H Genes Mutated at Codon 61. Cell 1986, 44 (1), 167–176. 10.1016/0092-8674(86)90495-2.

(45) Trahey, M.; Mccormick, F. A Cytoplasmic Protein Stimulates Normal N-Ras P21 GTPase, but Does Not Affect Oncogenic Mutants. Science 1987, 238 (4826), 542–545. 10.1126/science.2821624.

(46) Pai, E. F.; Krengel, U.; Petsko, G. A.; Goody, R. S.; Kabsch, W.; Wittinghofer, A. Refined Crystal Structure of the Triphosphate Conformation of H-Ras P21 at 1.35 A Resolution: Implications for the Mechanism of GTP Hydrolysis. EMBO J. 1990, 9 (8), 2351–2359. 10.1002/j.1460-2075.1990.tb07409.x.

(47) Scheffzek, K.; Ahmadian, M. R.; Kabsch, W.; Wiesmüller, L.; Lautwein, A.; Schmitz, F.; Wittinghofer, A. The Ras-RasGAP Complex: Structural Basis for GTPase Activation and Its Loss in Oncogenic Ras Mutants. Science 1997, 277 (5324), 333–339. 10.1126/science.277.5324.333.

(48) Buhrman, G.; Holzapfel, G.; Fetics, S.; Mattos, C. Allosteric Modulation of Ras Positions Q61 for a Direct Role in Catalysis. Proc. Natl. Acad. Sci. 2010, 107 (11), 4931–4936. 10.1073/pnas.0912226107.

(49) Hunter, J. C.; Manandhar, A.; Carrasco, M. a; Gurbani, D.; Gondi, S.; Westover, K. D. Biochemical and Structural Analysis of Common Cancer-Associated KRAS Mutations. Mol. Cancer Res. MCR 2015, 13 (9), 1325–1336. 10.1158/1541-7786.MCR-15-0203.

(50) Alexander, P.; Chan, A. H.; Rabara, D.; Swain, M.; Larsen, E. K.; Dyba, M.; Chertov, O.; Ashraf, M.; Champagne, A.; Lin, K.; Maciag, A.; Gillette, W. K.; Nissley, D. V.; McCormick, F.; Simanshu, D. K.; Stephen, A. G. Biophysical and Structural Analysis of KRAS Switch-II Pocket Inhibitors Reveals Allele-Specific Binding Constraints. J. Biol. Chem. 2025, 301 (7), 110331. 10.1016/j.jbc.2025.110331.

(51) McCormick, F. K-Ras Protein as a Drug Target. J. Mol. Med. 2016, 94 (3), 253–258. 10.1007/s00109-016-1382-7.

(52) Schweins, T.; Geyer, M.; Scheffzek, K.; Warshel, A.; Kalbitzer, H. R.; Wittinghofer, A. Substrate-Assisted Catalysis as a Mechanism for GTP Hydrolysis of P21ras and Other GTP-Binding Proteins. Nat. Struct. Biol. 1995, 2 (1), 36–44. 10.1038/nsb0195-36.

(53) Ahmadian, M. R.; Zor, T.; Vogt, D.; Kabsch, W.; Selinger, Z.; Wittinghofer, a; Scheffzek, K. Guanosine Triphosphatase Stimulation of Oncogenic Ras Mutants. Proc. Natl. Acad. Sci. U. S. A. 1999, 96 (12), 7065–7070. 10.1073/pnas.96.12.7065.

(54) Toney, M. D.; Kirsch, J. F. Direct Brønsted Analysis of the Restoration of Activity to a Mutant Enzyme by Exogenous Amines. Science 1989, 243 (4897), 1485–1488. 10.1126/science.2538921.

(55) Craik, C. S.; Roczniak, S.; Largman, C.; Rutter, W. J. The Catalytic Role of the Active Site Aspartic Acid in Serine Proteases. Science 1987, 237 (4817), 909–913. 10.1126/science.3303334.

(56) Carter, P.; Wells, J. A. Engineering Enzyme Specificity by “Substrate-Assisted Catalysis.” Science 1987, 237 (4813), 394–399. 10.1126/science.3299704.

(57) Qiao, Y.; Molina, H.; Pandey, A.; Zhang, J.; Cole, P. A. Chemical Rescue of a Mutant Enzyme in Living Cells. Science 2006, 311 (5765), 1293–1297. 10.1126/science.1122224.

(58) O’Meara, T. R.; Palanski, B. A.; Chen, M.; Qiao, Y.; Cole, P. A. Mutant Protein Chemical Rescue: From Mechanisms to Therapeutics. J. Biol. Chem. 2025, 301 (4), 108417. 10.1016/j.jbc.2025.108417.

(59) Voorhees, R. M.; Schmeing, T. M.; Kelley, A. C.; Ramakrishnan, V. The Mechanism for Activation of GTP Hydrolysis on the Ribosome. Science 2010, 330 (6005), 835–838. 10.1126/science.1194460.

(60) Wang, X.; Allen, S.; Blake, J. F.; Bowcut, V.; Briere, D. M.; Calinisan, A.; Dahlke, J. R.; Fell, J. B.; Fischer, J. P.; Gunn, R. J.; Hallin, J.; Laguer, J.; Lawson, J. D.; Medwid, J.; Newhouse, B.; Nguyen, P.; O’Leary, J. M.; Olson, P.; Pajk, S.; Rahbaek, L.; Rodriguez, M.; Smith, C. R.; Tang, T. P.; Thomas, N. C.; Vanderpool, D.; Vigers, G. P.; Christensen, J. G.; Marx, M. A. Identification of MRTX1133, a Noncovalent, Potent, and Selective KRAS^G12D^ Inhibitor. J. Med. Chem. 2022, 65 (4), 3123–3133. 10.1021/acs.jmedchem.1c01688.

(61) Gebregiworgis, T.; Kano, Y.; St-Germain, J.; Radulovich, N.; Udaskin, M. L.; Mentes, A.; Huang, R.; Poon, B. P. K.; He, W.; Valencia-Sama, I.; Robinson, C. M.; Huestis, M.; Miao, J.; Yeh, J. J.; Zhang, Z.-Y.; Irwin, M. S.; Lee, J. E.; Tsao, M.-S.; Raught, B.; Marshall, C. B.; Ohh, M.; Ikura, M. The Q61H Mutation Decouples KRAS from Upstream Regulation and Renders Cancer Cells Resistant to SHP2 Inhibitors. Nat. Commun. 2021, 12 (1), 6274. 10.1038/s41467-021-26526-y.

(62) Rennella, E.; Henry, C.; Dickson, C. J.; Georgescauld, F.; Wales, T. E.; Erdmann, D.; Cotesta, S.; Maira, M.; Sedrani, R.; Brachmann, S. M.; Ostermann, N.; Engen, J. R.; Kay, L. E.; Beyer, K. S.; Wilcken, R.; Jahnke, W. Dynamic Conformational Equilibria in the Active States of KRAS and NRAS. RSC Chem. Biol. 2025, 6 (1), 106–118. 10.1039/D4CB00233D.

(63) Holderfield, M.; Lee, B. J.; Jiang, J.; Tomlinson, A.; Seamon, K. J.; Mira, A.; Patrucco, E.; Goodhart, G.; Dilly, J.; Gindin, Y.; Dinglasan, N.; Wang, Y.; Lai, L. P.; Cai, S.; Jiang, L.; Nasholm, N.; Shifrin, N.; Blaj, C.; Shah, H.; Evans, J. W.; Montazer, N.; Lai, O.; Shi, J.; Ahler, E.; Quintana, E.; Chang, S.; Salvador, A.; Marquez, A.; Cregg, J.; Liu, Y.; Milin, A.; Chen, A.; Ziv, T. B.; Parsons, D.; Knox, J. E.; Klomp, J. E.; Roth, J.; Rees, M.; Ronan, M.; Cuevas-Navarro, A.; Hu, F.; Lito, P.; Santamaria, D.; Aguirre, A. J.; Waters, A. M.; Der, C. J.; Ambrogio, C.; Wang, Z.; Gill, A. L.; Koltun, E. S.; Smith, J. A. M.; Wildes, D.; Singh, M. Concurrent Inhibition of Oncogenic and Wild-Type RAS-GTP for Cancer Therapy. Nature 2024, 629 (8013), 919–926. 10.1038/s41586-024-07205-6.

(64) Cuevas-Navarro, A.; Pourfarjam, Y.; Hu, F.; Rodriguez, D. J.; Vides, A.; Sang, B.; Fan, S.; Goldgur, Y.; De Stanchina, E.; Lito, P. Pharmacological Restoration of GTP Hydrolysis by Mutant RAS. Nature 2025, 637 (8044), 224–229. 10.1038/s41586-024-08283-2.

(65) Sharma, A. K.; Pei, J.; Yang, Y.; Dyba, M.; Smith, B.; Rabara, D.; Larsen, E. K.; Lightstone, F. C.; Esposito, D.; Stephen, A. G.; Wang, B.; Beltran, P. J.; Wallace, E.; Nissley, D. V.; McCormick, F.; Maciag, A. E. Revealing the Mechanism of Action of a First-in-Class Covalent Inhibitor of KRASG12C (ON) and Other Functional Properties of Oncogenic KRAS by 31P NMR. J. Biol. Chem. 2024, 300 (2), 105650. 10.1016/j.jbc.2024.105650.

(66) Pan, X.; Wang, H.; Li, C.; Zhang, J. Z. H.; Ji, C. MolGpka: A Web Server for Small Molecule p *K*_a_ Prediction Using a Graph-Convolutional Neural Network. J. Chem. Inf. Model. 2021, 61 (7), 3159–3165. 10.1021/acs.jcim.1c00075.

(67) Alexander, P.; Chan, A. H.; Rabara, D.; Swain, M.; Larsen, E. K.; Dyba, M.; Chertov, O.; Ashraf, M.; Champagne, A.; Lin, K.; Maciag, A.; Gillette, W. K.; Nissley, D. V.; McCormick, F.; Simanshu, D. K.; Stephen, A. G. Biophysical and Structural Analysis of KRAS Switch-II Pocket Inhibitors Reveals Allele-Specific Binding Constraints. J. Biol. Chem. 2025, 301 (7), 110331. 10.1016/j.jbc.2025.110331.

(68) Vasta, J. D.; Peacock, D. M.; Zheng, Q.; Walker, J. A.; Zhang, Z.; Zimprich, C. A.; Thomas, M. R.; Beck, M. T.; Binkowski, B. F.; Corona, C. R.; Robers, M. B.; Shokat, K. M. KRAS Is Vulnerable to Reversible Switch-II Pocket Engagement in Cells. Nat. Chem. Biol. 2022, 18 (6), 596–604. 10.1038/s41589-022-00985-w.

(69) Lechuga, C. G.; Salmón, M.; Paniagua, G.; Guerra, C.; Barbacid, M.; Drosten, M. RASless MEFs as a Tool to Study RAS-Dependent and RAS-Independent Functions. In Ras Activity and Signaling: Methods and Protocols; Rubio, I., Prior, I., Eds.; Springer US: New York, NY, 2021; pp 335–346. 10.1007/978-1-0716-1190-6_21.

(70) Jencks, W. P.; Carriuolo, J. Imidazole Catalysis: III. GENERAL BASE CATALYSIS AND THE REACTIONS OF ACETYL IMIDAZOLE WITH THIOLS AND AMINES. J. Biol. Chem. 1959, 234 (5), 1280–1285. 10.1016/S0021-9258(18)98173-1.

(71) Bruice, T. C.; Fife, T. H.; Bruno, J. J.; Benkovic, Patricia. Hydroxyl Group (V)1 and Imidazole (X)2 Catalysis. The General Base Catalysis of Ester Hydrolysis by Imidazole and the Influence of a Neighboring Hydroxyl Group. J. Am. Chem. Soc. 1962, 84 (15), 3012–3018. 10.1021/ja00874a035.

(72) Breslow, R. Bifunctional Acid—Base Catalysis by Imidazole Groups in Enzyme Mimics. J. Mol. Catal. 1994, 91 (2), 161–174. 10.1016/0304-5102(94)00046-8.

(73) Thompson, J. E.; Raines, R. T. Value of General Acid-Base Catalysis to Ribonuclease A. J. Am. Chem. Soc. 1994, 116 (12), 5467–5468. 10.1021/ja00091a060.

(74) Tripp, B. C.; Smith, K.; Ferry, J. G. Carbonic Anhydrase: New Insights for an Ancient Enzyme * 210. J. Biol. Chem. 2001, 276 (52), 48615–48618. 10.1074/jbc.R100045200.

(75) Mikutis, S.; Rebelo, M.; Yankova, E.; Gu, M.; Tang, C.; Coelho, A. R.; Yang, M.; Hazemi, M. E.; Pires De Miranda, M.; Eleftheriou, M.; Robertson, M.; Vassiliou, G. S.; Adams, D. J.; Simas, J. P.; Corzana, F.; Schneekloth, J. S.; Tzelepis, K.; Bernardes, G. J. L. Proximity-Induced Nucleic Acid Degrader (PINAD) Approach to Targeted RNA Degradation Using Small Molecules. ACS Cent. Sci. 2023, 9 (5), 892–904. 10.1021/acscentsci.3c00015.

(76) Cregg, J.; Edwards, A. V.; Chang, S.; Lee, B. J.; Knox, J. E.; Tomlinson, A. C. A.; Marquez, A, ; Liu, Y.; Freilich, R.; Aay, N.; Wang, Y.; Jiang, L.; Jiang, J.; Wang, Z.; Flagella, M.; Wildes, D.; Smith, J. A. M.; Singh, M.; Wang, Z.; Gill, A. L.; Koltun, E. S. Discovery of Daraxonrasib (RMC-6236), a Potent and Orally Bioavailable RAS(ON) Multi-Selective, Noncovalent Tri-Complex Inhibitor for the Treatment of Patients with Multiple RAS-Addicted Cancers. J. Med. Chem. 2025, 68 (6), 6064–6083. 10.1021/acs.jmedchem.4c02314.

(77) Parry, C. W.; Pellicano, F.; Schüttelkopf, A. W.; Beyer, K. S.; Bower, J.; Bryson, A.; Cameron, K.; Cerutti, N. M.; Clark, J. P.; Davidson, S. C.; Davies, K.; Drysdale, M. J.; Engelman, J.; Estevan-Barber, A.; Gohlke, A.; Gray, C. H.; Guthy, D. A.; Hong, M.; Hopkins, A.; Hutchinson, L. D.; Konczal, J.; Maira, M.; McArthur, D.; Mezna, M.; McKinnon, H.; Nepravishta, R.; Ostermann, N.; Pasquali, C. C.; Pollock, K.; Pugliese, A.; Rooney, N.; Schmiedeberg, N.; Shaw, P.; Velez-Vega, C.; West, C.; West, R.; Zecri, F.; Taylor, J. B. Reversible Small Molecule Multivariant Ras Inhibitors Display Tunable Affinity for the Active and Inactive Forms of Ras. J. Med. Chem. 2025, 68 (9), 9129–9161. 10.1021/acs.jmedchem.4c02929.

(78) Patel, S.; Bhhatarai, B.; Calses, P.; Erlanson, D.; Everley, R.; Fong, S.; Gerken, P.; Hermann, J. C.; Le, T.; Liu, L.; McMahon, E.; Neve, R. M.; Phan, T.; Roberts, A.; Shanafelt, M.; Siemsgluess, S.; Staunton, J.; Wang, Y.; Wang, W.; Williams, M.; Webster, K. R. Abstract 1142: Discovery of FMC-376 a Novel Orally Bioavailable Inhibitor of Activated KRASG12C. Cancer Res. 2023, 83 (7_Supplement), 1142–1142. 10.1158/1538-7445.AM2023-1142.

(79) Cregg, J.; Pota, K.; Tomlinson, A. C. A.; Yano, J.; Marquez, A.; Liu, Y.; Schulze, C. J.; Seamon, K. J.; Holderfield, M.; Wei, X.; Zhuang, Y.; Yang, Y. C.; Jiang, J.; Huang, Y.; Zhao, R.; Ling, Y.; Wang, Z.; Flagella, M.; Wang, Z.; Singh, M.; Knox, J. E.; Nichols, R.; Wildes, D.; Smith, J. A. M.; Koltun, E. S.; Gill, A. L. Discovery of Elironrasib (RMC-6291), a Potent and Orally Bioavailable, RAS(ON) G12C-Selective, Covalent Tricomplex Inhibitor for the Treatment of Patients with RAS G12C-Addicted Cancers. J. Med. Chem. 2025, 68 (6), 6041–6063. 10.1021/acs.jmedchem.4c02313.

(80) Bondeva, T.; Balla, A.; Várnai, P.; Balla, T. Structural Determinants of Ras-Raf Interaction Analyzed in Live Cells. Mol. Biol. Cell 2002, 13 (7), 2323–2333. 10.1091/mbc.e02-01-0019.

(81) Brtva, T. R.; Drugan, J. K.; Ghosh, S.; Terrell, R. S.; Campbell-Burk, S.; Bell, R. M.; Der, C. J. Two Distinct Raf Domains Mediate Interaction with Ras. J. Biol. Chem. 1995, 270 (17), 9809–9812. 10.1074/jbc.270.17.9809.

(82) KRAS: Methods and Protocols; Stephen, A. G., Esposito, D., Eds.; Methods in Molecular Biology; Springer US: New York, NY, 2024; Vol. 2797. 10.1007/978-1-0716-3822-4.

