## Supplementary Information for "Brønsted-basic small molecules activate GTP hydrolysis in Ras Q61 mutants"

#### Table of Contents

**Part 1. Supplemental Figure and tables**........S3

**Part 2. Materials and Reagents**........S8

Protein sequence list........S10

List of primers used in this study and their sequences........S12

List of primary antibodies used in this study........S12

**Part 3. Unprocessed Western Blots**........S13

**Part 4. Chemical Synthesis**........S18

**Part 5. $^{1}$H NMR, $^{19}$F NMR and $^{13}$C NMR Spectra**........S35

**Part 6. Reference**........S56

#### Part 1. Supplemental Figure and Tables.

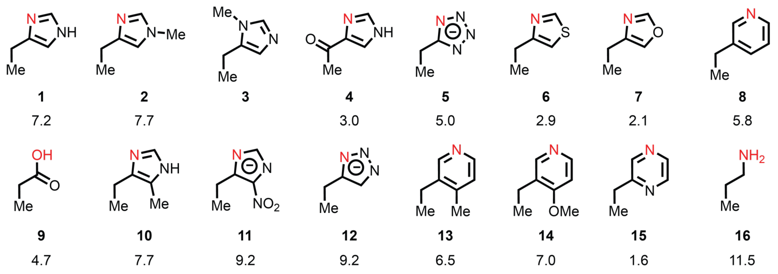


**Supplemental Figure 1.** Partial structures used for p$K_a$ prediction[1] and the calculated p$K_a$ values of the conjugate acids of the structures shown.

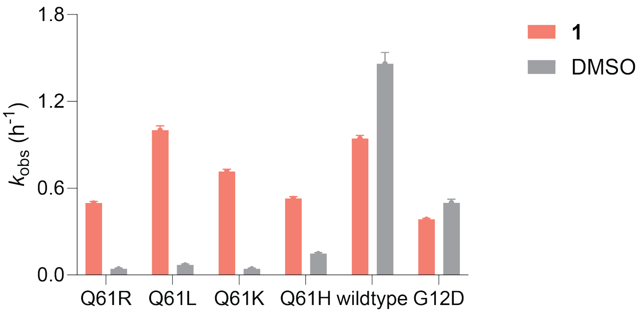


**Supplemental Figure 2.** Apparent first-order reaction rate constants for GTP hydrolysis of K-Ras wildtype or mutants in the presence of DMSO or compound **1** (50 μM).

**Table S1.** GTP hydrolysis rate $k$ ($h^{-1}$), SEM ($k$), SD ($k$), half-life $t_{1/2}$ (h), fold-change versus DMSO (x DMSO), SD (x DMSO) on K-RAS(Q61R) under compound treatment. (50 uM cpds, 1% DMSO, 1.5 uM Ras·GTP, 0-2h data, $N$=3).

| Cpd | $k$ ($h^{-1}$) | SEM ($k$) | SD ($k$) | $t_{1/2}$ (h) | x DMSO | SD (x DMSO) |
|---|---|---|---|---|---|---|
| **1** | 0.3241 | 0.003364 | 0.005827 | 2.1 | 21.1 | 5.88 |
| **2** | 0.1293 | 0.002498 | 0.004327 | 5.4 | 8.4 | 2.36 |
| **3** | 0.02352 | 0.002109 | 0.003653 | 29.5 | 1.5 | 0.49 |
| **4** | 0.0213 | 0.002694 | 0.004666 | 32.5 | 1.4 | 0.49 |
| **5** | 0.1225 | 0.003339 | 0.005783 | 5.7 | 8.0 | 2.25 |
| **6** | 0.08134 | 0.003612 | 0.006256 | 8.5 | 5.3 | 1.53 |
| **7** | 0.03214 | 0.002768 | 0.004794 | 21.6 | 2.1 | 0.66 |
| **8** | 0.2732 | 0.006001 | 0.010394 | 2.5 | 17.8 | 5.00 |
| **9** | 0.129 | 0.003493 | 0.00605 | 5.4 | 8.4 | 2.37 |
| **10** | 0.08061 | 0.003942 | 0.006828 | 8.6 | 5.3 | 1.53 |
| **11** | 0.3185 | 0.006388 | 0.011064 | 2.2 | 20.8 | 5.81 |
| **12** | 0.0913 | 0.002686 | 0.004652 | 7.6 | 6.0 | 1.68 |
| **13** | 0.3793 | 0.006250 | 0.010825 | 1.8 | 24.7 | 6.91 |
| **14** | 0.3445 | 0.004401 | 0.007623 | 2.0 | 22.5 | 6.26 |
| **15** | 0.02911 | 0.002564 | 0.004441 | 23.8 | 1.9 | 0.60 |
| **16** | 0.01155 | 0.002667 | 0.004619 | 60.0 | 0.8 | 0.37 |
| DMSO | 0.01533 | 0.002458 | 0.004257 | 45.2 | 1.0 | 0.39 |

**Table S2.** GTP hydrolysis rate ($h^{-1}$) by K-RAS(Q61X) after treatment with different concentrations of compound **1**. (1% DMSO, 1.5 uM Ras·GTP, 0-2 h data, *N*=3)

| c (μM) | K-RAS(Q61R) | | K-RAS(Q61L) | | K-RAS(Q61K) | | K-RAS(Q61H) | |
|---|---|---|---|---|---|---|---|---|
| | $k$($h^{-1}$) | SD | $k$($h^{-1}$) | SD | $k$($h^{-1}$) | SD | $k$($h^{-1}$) | SD |
| 100 | 0.597 | 0.014 | 0.682 | 0.023 | 0.546 | 0.017 | 0.392 | 0.014 |
| 50 | 0.497 | 0.011 | 0.665 | 0.011 | 0.582 | 0.010 | 0.453 | 0.007 |
| 25 | 0.375 | 0.006 | 0.601 | 0.009 | 0.630 | 0.010 | 0.475 | 0.008 |
| 12.5 | 0.237 | 0.005 | 0.402 | 0.007 | 0.494 | 0.009 | 0.382 | 0.012 |
| 6.25 | 0.111 | 0.004 | 0.203 | 0.008 | 0.263 | 0.007 | 0.362 | 0.006 |
| 3.13 | 0.049 | 0.004 | 0.070 | 0.004 | 0.095 | 0.005 | 0.198 | 0.005 |
| 1.56 | 0.026 | 0.004 | 0.056 | 0.006 | 0.039 | 0.004 | 0.153 | 0.005 |
| 0.78 | 0.019 | 0.004 | 0.045 | 0.005 | 0.032 | 0.004 | 0.136 | 0.005 |
| 0.39 | 0.018 | 0.004 | 0.053 | 0.004 | 0.035 | 0.004 | 0.132 | 0.004 |
| 0.20 | 0.018 | 0.004 | 0.052 | 0.004 | 0.030 | 0.005 | 0.144 | 0.005 |
| 0.10 | 0.018 | 0.005 | 0.056 | 0.005 | 0.038 | 0.004 | 0.144 | 0.004 |
| 0 | 0.017 | 0.003 | 0.042 | 0.004 | 0.037 | 0.004 | 0.120 | 0.004 |

**Table S3.** $EC_{50}$ (μM) and 95% CI (μM) on K-RAS(Q61X) under compound **1** treatment.

| RAS | $EC_{50}$ (μM) | 95% CI (μM) |
|---|---|---|
| K-RAS(Q61R)•GTP | 20.9 | 19.5 – 22.7 |
| K-RAS(Q61L)•GTP | 11.1 | 10.7 – 11.5 |
| K-RAS(Q61K)•GTP | 7.0 | 6.5 – 7.5 |
| K-RAS(Q61H)•GTP | 4.6 | 4.0 – 5.2 |

**Table S4.** $K_d$ (nM) and 95% CI (nM) of the tracer molecule **20** against K-RAS(Q61R)•GppNHp and K-RAS(Q61L)•GppNHp, determined by fluorescence polarization.

| RAS | $K_d$ (nM) | 95% CI (nM) |
|---|---|---|
| K-RAS(Q61R)•GppNHp | 341.7 | 280.3 – 417.4 |
| K-RAS(Q61L)•GppNHp | 99.4 | 74.0 – 133.4 |

**Table S5.** Fluorescence polarization (FP)-derived $IC_{50}$ values and apparent dissociation constants ($K_d$) for representative compounds binding to K-Ras(Q61R)•GppNHp.

| Compound | $IC_{50}$ (nM) | 95% CI (nM) | $K_d$ (μM) | 95% CI (μM) |
|---|---|---|---|---|
| **1** | 9304 | 6252 – 13895 | 2.2 | 1.4 – 3.3 |
| **2** | 9677 | 6196 – 15157 | 2.3 | 1.4 – 3.7 |
| **3** | 8893 | 5326 – 15515 | 2.1 | 1.1 – 3.8 |
| **4** | 18408 | 9718 – 36046 | 4.5 | 2.3 – 9.0 |
| **8** | 23475 | 12601 – 54793 | 5.8 | 3.0 – 13.8 |
| **9** | 28932 | 17749 – 49190 | 7.2 | 4.3 – 12.3 |
| **11** | 14617 | 9415 – 24128 | 3.5 | 2.2 – 6.0 |
| **13** | 43209 | 15397 – | 10.8 | 3.7 – |
| **14** | 54369 | 33694 – 95936 | 13.7 | 8.4 – 24.3 |
| **16** | 4676 | 3205 – 6812 | 1.0 | 0.6 – 1.5 |
| MRTX1133 | 2130 | 1344–3369 | 0.3 | 0.1 – 0.7 |

**Table S6.** Fluorescence polarization (FP)–derived $IC_{50}$ values and apparent dissociation constants ($K_d$) for representative compounds binding to K-RAS(Q61L)•GppNHp.

| Compound | $IC_{50}$ (nM) | 95% CI (nM) | $K_d$ (nM) | 95% CI (nM) |
|---|---|---|---|---|
| **1** | 2508 | 1788 – 3510 | 783.3 | 544.6 – 1115.5 |
| **2** | 1360 | 831.1 – 2198 | 402.7 | 227.4 – 680.6 |
| **3** | 1787 | 1125 – 2839 | 544.3 | 324.8 – 893.1 |
| **4** | 16777 | 10274 – 27932 | 5514 | 3358 – 9212 |
| **8** | 4957 | 2880 – 8637 | 1595 | 906.7 – 2815 |
| **9** | 5435 | 3450 – 8518 | 1754 | 1096 – 2776 |
| **11** | 2811 | 1659 – 4762 | 883.8 | 501.9 – 1530.6 |
| **13** | 23500 | 11749 – 63754 | 7743 | 3847 – 21088 |
| **14** | 8572 | 5605 – 13160 | 2794 | 1810 – 4315 |
| **16** | 946.3 | 649.1 – 1383 | 265.6 | 167.0 – 410.3 |
| MRTX1133 | 463.8 | 312.5 – 690.2 | 105.6 | 55.4 – 180.7 |

#### Part 2. Materials and Reagents

##### Protein sequence list.

Blue texts indicate affinity tag that were removed during the purification step.

K-RAS(wt):
MHHHHHHSSGRENLYFQGMTEYKLVVVGAGGVGKSALTIQLIQNHFVDEYDPTIEDSYRKQV
VIDGETCLLDILDTAGQEEYSAMRDQYMRTGEGFLCVFAINNTKSFEDIHHYREQIKRVKDS
EDVPMVLVGNKCDLPSRTVDTKQAQDLARSYGIPFIETSAKTRQGVDDAFYTLVREIRKHKE
K

K-RAS(G12D):
MHHHHHHSSGRENLYFQGMTEYKLVVVGADGVGKSALTIQLIQNHFVDEYDPTIEDSYRKQV
VIDGETCLLDILDTAGQEEYSAMRDQYMRTGEGFLCVFAINNTKSFEDIHHYREQIKRVKDS
EDVPMVLVGNKCDLPSRTVDTKQAQDLARSYGIPFIETSAKTRQGVDDAFYTLVREIRKHKE
K

K-RAS(Q61R):
MHHHHHHSSGRENLYFQGMTEYKLVVVGAGGVGKSALTIQLIQNHFVDEYDPTIEDSYRKQV
VIDGETCLLDILDTAGREEYSAMRDQYMRTGEGFLCVFAINNTKSFEDIHHYREQIKRVKDS
EDVPMVLVGNKCDLPSRTVDTKQAQDLARSYGIPFIETSAKTRQGVDDAFYTLVREIRKHKE
K

K-RAS(Q61L):
MHHHHHHSSGRENLYFQGMTEYKLVVVGAGGVGKSALTIQLIQNHFVDEYDPTIEDSYRKQV
VIDGETCLLDILDTAGLEEYSAMRDQYMRTGEGFLCVFAINNTKSFEDIHHYREQIKRVKDS
EDVPMVLVGNKCDLPSRTVDTKQAQDLARSYGIPFIETSAKTRQGVDDAFYTLVREIRKHKE
K

K-RAS(Q61K):
MHHHHHHSSGRENLYFQGMTEYKLVVVGAGGVGKSALTIQLIQNHFVDEYDPTIEDSYRKQV
VIDGETCLLDILDTAGKEEYSAMRDQYMRTGEGFLCVFAINNTKSFEDIHHYREQIKRVKDS
EDVPMVLVGNKCDLPSRTVDTKQAQDLARSYGIPFIETSAKTRQGVDDAFYTLVREIRKHKE
K

K-RAS(Q61H):
MHHHHHHSSGRENLYFQGMTEYKLVVVGAGGVGKSALTIQLIQNHFVDEYDPTIEDSYRKQV
VIDGETCLLDILDTAGHEEYSAMRDQYMRTGEGFLCVFAINNTKSFEDIHHYREQIKRVKDS
EDVPMVLVGNKCDLPSRTVDTKQAQDLARSYGIPFIETSAKTRQGVDDAFYTLVREIRKHKE
K

H-RAS(Q61R):
MHHHHHHSSGRENLYFQGMTEYKLVVVGAGGVGKSALTIQLIQNHFVDEYDPTIEDSYRKQV

VIDGETCLLDILDTAGREEYSAMRDQYMRTGEGFLCVFAINNTKSFEDIHQYREQIKRVKDS
DDVPMVLVGNKCDLAARTVESRQAQDLARSYGIPYIETSAKTRQGVEDAFYTLVREIRQH

H-RAS(Q61R/Q95H):
MHHHHHHSSGRENLYFQGMTEYKLVVVGAGGVGKSALTIQLIQNHFVDEYDPTIEDSYRKQV
VIDGETCLLDILDTAGREEYSAMRDQYMRTGEGFLCVFAINNTKSFEDIHHYREQIKRVKDS
DDVPMVLVGNKCDLAARTVESRQAQDLARSYGIPYIETSAKTRQGVEDAFYTLVREIRQH

N-RAS(Q61R):
MHHHHHHSSGRENLYFQGMTEYKLVVVGAGGVGKSALTIQLIQNHFVDEYDPTIEDSYRKQV
VIDGETCLLDILDTAGREEYSAMRDQYMRTGEGFLCVFAINNSKSFADINLYREQIKRVKDS
DDVPMVLVGNKCDLPTRTVDTKQAHELAKSYGIPFIETSAKTRQGVEDAFYTLVREIRQY

N-RAS(Q61R/L95H):
MHHHHHHSSGRENLYFQGMTEYKLVVVGAGGVGKSALTIQLIQNHFVDEYDPTIEDSYRKQV
VIDGETCLLDILDTAGREEYSAMRDQYMRTGEGFLCVFAINNSKSFADINHYREQIKRVKDS
DDVPMVLVGNKCDLPTRTVDTKQAHELAKSYGIPFIETSAKTRQGVEDAFYTLVREIRQY

NF1-GRD:
MHHHHHHSSGRENLYFQGDRFERLVELVTMMGDQGELPIAMALANVVPCSQWDELARVLVTL
FDSRHLLYQLLWNMFSKEVELADSMQTLFRGNSLASKIMTFCFKVYGATYLQKLLDPLLRIV
ITSSDWQHVSFEVDPTRLEPSESLEENQRNLLQMTEKFFHAIISSSSEFPPQLRSVCHCLYQ
VVSQRFPQNSIGAVGSAMFLRFINPAIVSPYEAGILDKKPPPRIERGLKLMSKILQSIANHV
LFTKEEHMRPFNDFVKSNFDAARRFFLDIASDCPTSDAVNHSLSFISDGNVLALHRLLWNNQ
EKIGQYLSSNRDHKAVGRRPFDKMATLLAYLGPPEH

CYPA:
MAHHHHHHHHSAALEVLFQGPDMVNPTVFFDIAVDGEPLGRVSFELFADKVPKTAENFRALS
TGEKGFGYKGSCFHRIIPGFMCQGGDFTRHNGTGGKSIYGEKFEDENFILKHTGPGILSMAN
AGPNTNGSQFFICTAKTEWLDGKHVVFGKVKEGMNIVEAMERFGSRNGKTSKKITIADCGQL
E

GST-Raf1-RBD:
MSPILGYWKIKGLVQPTRLLLEYLEEKYEEHLYERDEGDKWRNKKFELGLEFPNLPYYIDGD
VKLTQSMAIIRYIADKHNMLGGCPKERAEISMLEGAVLDIRYGVSRIAYSKDFETLKVDFLS
KLPEMLKMFEDRLCHKTYLNGDHVTHPDFMLYDALDVVLYMDPMCLDAFPKLVCFKKRIEAI
PQIDKYLKSSKYIAWPLQGWQATFGGGDHPPKSDLVPRGSPIHIMEHIQGAWKTISNGFGFK
DAVFDGSSCISPTIVQQFGYQRRASDDGKLTDPSKTSNTIRVFLPNKQRTVVNVRNGMSLHD
CLMKALKVRGLQPECCAVFRLLHEHKGKKARLDWNTDAASLIGEELQVDFLDHVPLTTHNFA
RKTFLKLGIHRD

**List of primers used in this study and their sequences.**

```
K-Ras(Q61H)F: 5′-CACTGCCGGCCATGAGGAATACTCGGCCATG-3′
K-Ras(Q61H)R: 5′-GTATTCCTCATGGCCGGCAGTGTCTAAAATATCG-3′
H-Ras(Q95H)F: 5′-GATATTCATCAtTATCGTGAACAAATTAAGCGCG-3′
H-Ras(Q95H)R: 5′-GTTCACGATAaTGATGAATATCTTCGAAACTTTTCG-3′
L-Ras(L95H)F: 5′-CATCAATCatTATCGCGAACAAATCAAACG-3′
L-Ras(L95H)R: 5′-CCGCGATAatGATTGATGTCTGCGAAAC-3′
```

**List of primary antibodies used in this study.**

| Antibody | Host species | Supplier | Catalog No. | Dilution |
|---|---|---|---|---|
| Pan-Ras [EPR3255] | Rabbit mAb | abcam | ab108602 | 1:10,000 |
| Total ERK | Mouse mAb | Cell Signaling Technology | 4696 | 1:5,000 |
| P-ERK [T202/Y204] | Rabbit mAb | Cell Signaling Technology | 9101 | 1:5,000 |
| GAPDH | Mouse mAb | Proteintech | 60004 | 1:50,000 |

### Part 3. Unprocessed Western Blots.

Uncropped gel images for Figure 5A
Rasless MEF (KRas$^{Q61R}$), 4h treatment by **2**

*Membrane 1*

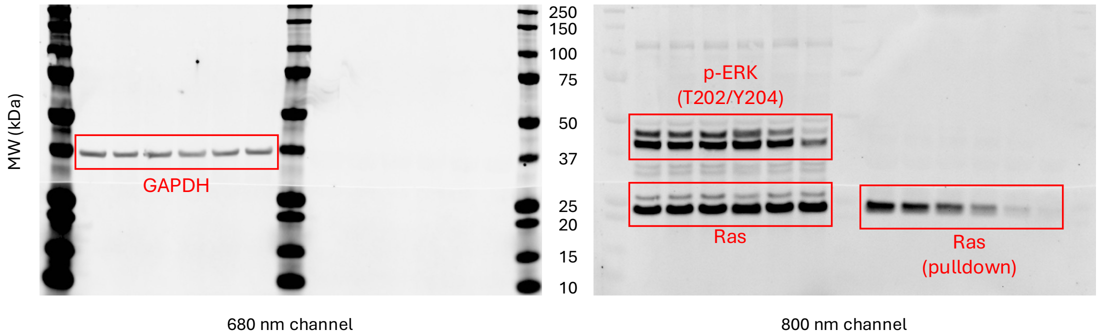


Uncropped gel images for Figure 5A
Rasless MEF (KRas$^{Q61R}$), 4h treatment by **2**

*Membrane 2*

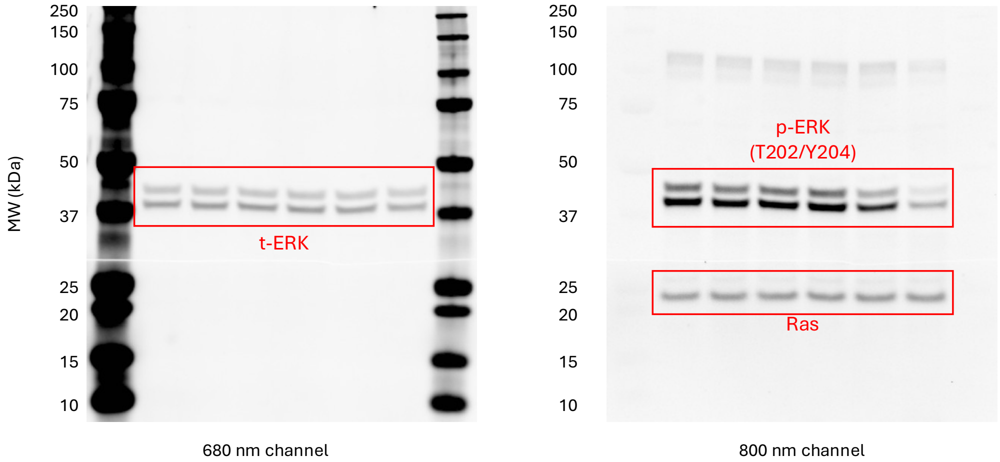

Uncropped gel images for Figure 5B
Rasless MEF (KRas$^{Q61R}$), treatment by 30 μM **2**

*Membrane 1*

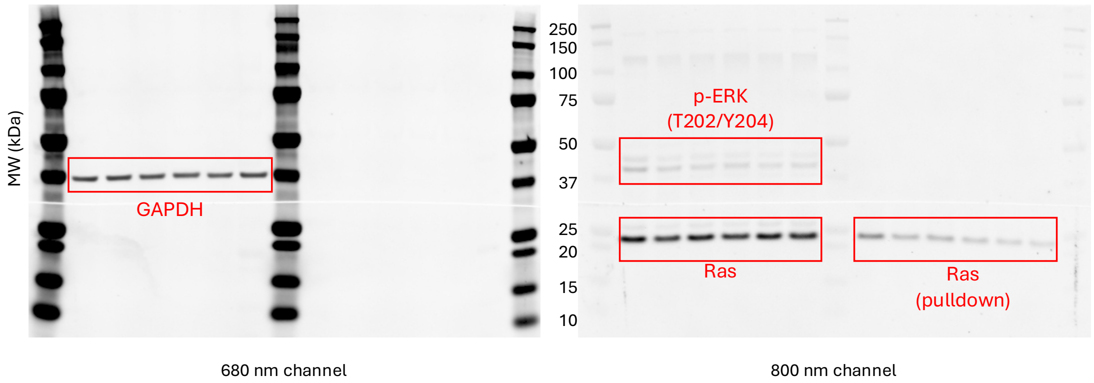


Uncropped gel images for Figure 5B
Rasless MEF (KRas$^{Q61R}$), treatment by 30 μM **2**

*Membrane 2*

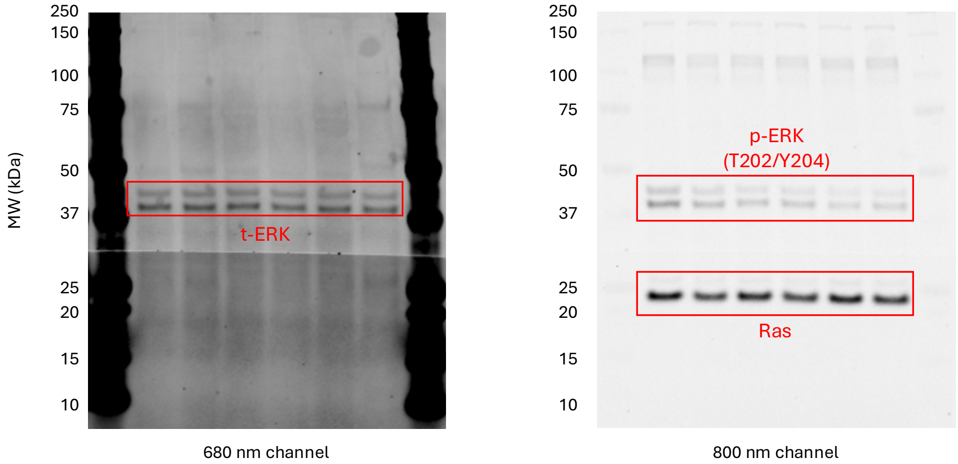

Uncropped gel images for Figure 5C

Rasless MEF (KRas$^{Q61L}$)

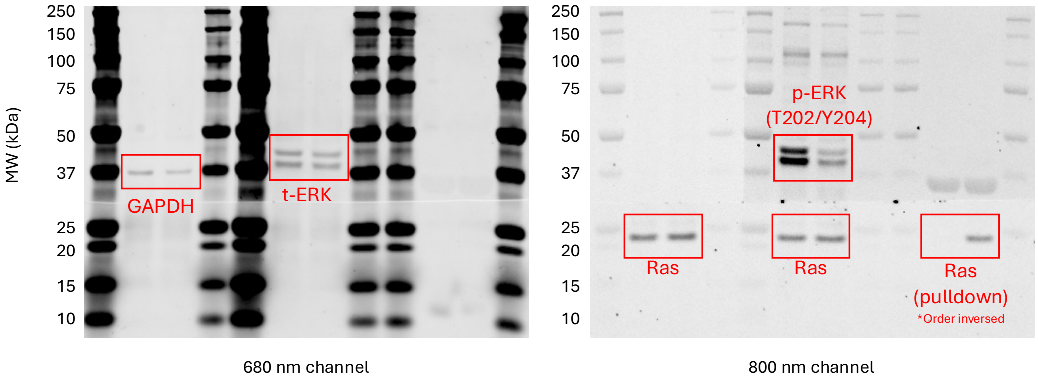


Uncropped gel images for Figure 5D

SW948 (KRas$^{Q61L/WT}$)

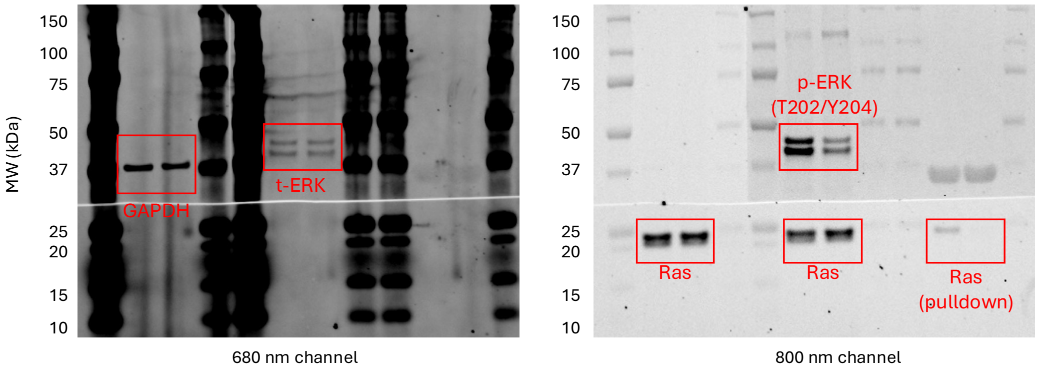

Uncropped gel images for Figure 5D
Calu-6 (KRas$^{Q61K/WT}$)

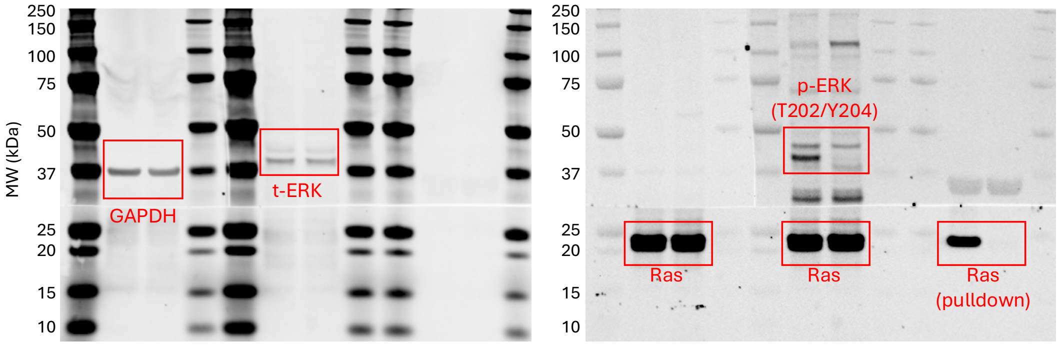


680 nm channel

800 nm channel

Uncropped gel images for Figure 5D
NCI-H460 (KRas$^{Q61H/Q61H}$)

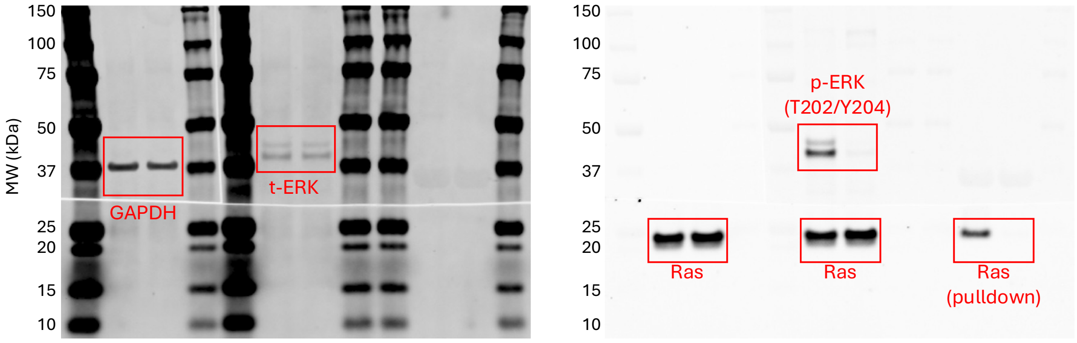


680 nm channel

800 nm channel

Uncropped gel images for Figure 5E
Rasless MEF ($BRaf^{V600E}$)

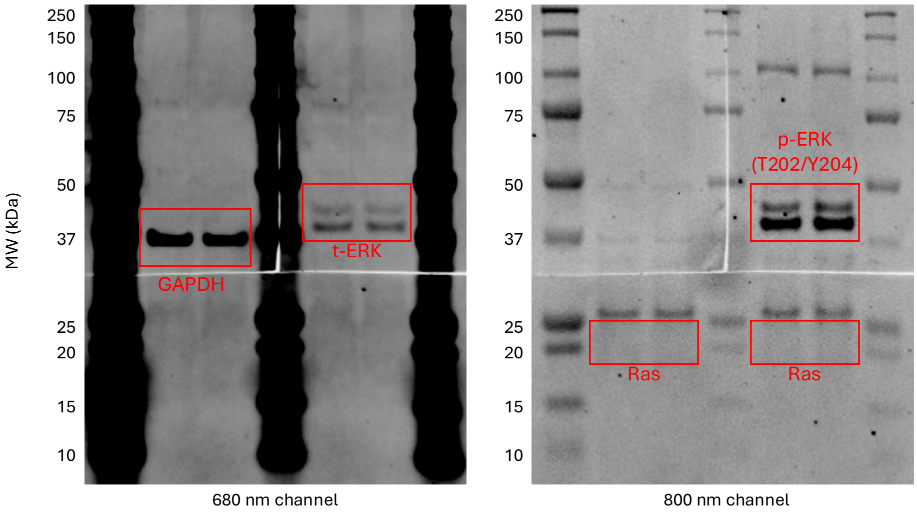


Uncropped gel images for Figure 5E
A375 ($BRaf^{V600E/V600E}$)

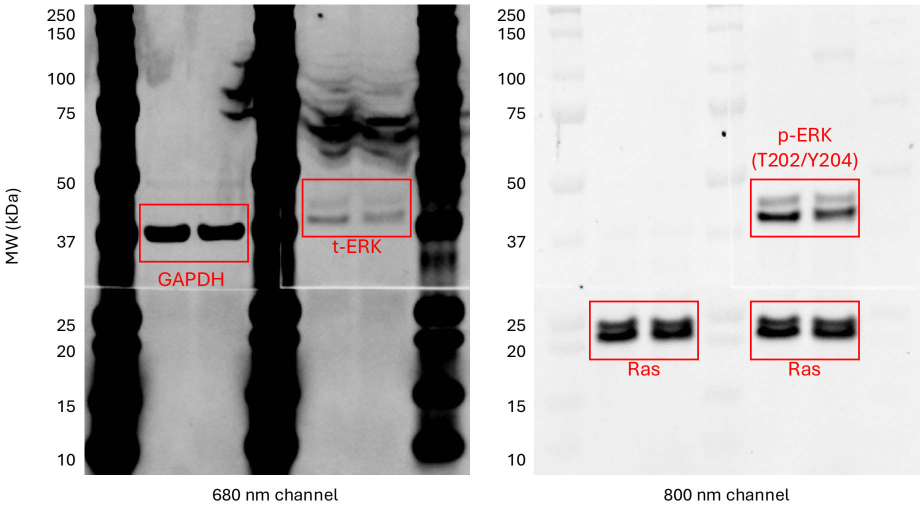

**Part 4. Chemical Synthesis.**

**General methods for chemical synthesis.** NMR spectra were recorded on two 500 MHz Bruker Avance NEO spectrometers ($^{1}H$ at 499.99 and 500.17 MHz, $^{13}C$ at 125.72 and 125.77 MHz, $^{19}F$ at 470.47 and 470.63 MHz, $^{31}P$ at 242.40 and 202.47 MHz) equipped with iProbe and Prodigy Cryoprobes, respectively. Chemical shifts (δ) were given in ppm with reference to d-solvent signals [$^{1}H$ NMR: $CHCl_3$ (7.26), $CD_3OD$ (3.31), $^{13}C$ NMR: $CD_3OD$ (49.0)]. Column chromatography was performed on silica gel (RediSep® Silver Silica Gel Disposable Flash Columns 4 grams from Teledyne ISCO, 692203304) or preparative high-performance liquid chromatography (prepHPLC, CombiFlash EZprep, C18 20x150mm column). All reactions sensitive to air or moisture were conducted under nitrogen atmosphere in dry solvents under anhydrous conditions, unless otherwise noted. MRTX1133 is purchased from Ambeed, A1501270, 98%. All other solvents and reagents were used as obtained from commercial sources: Fisher Scientific, Ambeed, Sigma Aldrich, AK Scientific and Chemscene without further purification.

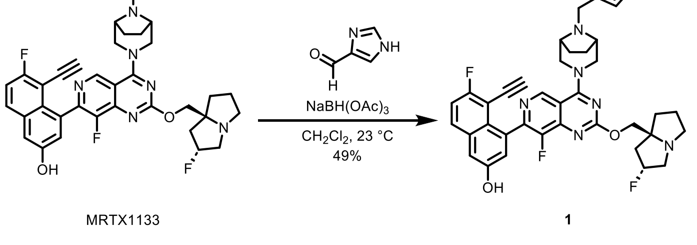


MRTX1133 1

**Compound 1.** A 3-mL vial was charged with MRTX1133 (11.4 mg, 1 equiv., 19.0 μmol), 4-Imidazolecarboxaldehyde (17.0 mg, 9.30 equiv., 177 μmol), dichloromethane (1.27 mL) and a magnetic stir bar. Sodium triacetoxyborohydride (16.1 mg, 4.00 equiv., 75.9 μmol) was added to the stirred solution at 23 ºC. The reaction solution was stirred at 23 ºC for 16 h. The reaction mixture was concentrated under reduced pressure to afford the crude. The crude product was purified by prepHPLC (CombiFlash EZprep, C18 20x150mm column, 5–100% acetonitrile in water, 0.1% formic acid, 30 min) to afford imidazole **1** as pale orange solid 6.3 mg, 49% yield. **$^{1}$H NMR (500 MHz, MeOD)** δ 9.06 (s, 1H), 7.87 (dd, *J* = 9.2, 5.7 Hz, 1H), 7.84 (d, *J* = 1.2 Hz, 1H), 7.36 (d, *J* = 2.6 Hz, 1H), 7.33 (t, *J* = 8.9 Hz, 1H), 7.21 (d, *J* = 2.6 Hz, 1H), 7.17 (s, 1H), 5.50 (d, *J* = 52.3 Hz, 1H), 4.69 – 4.47 (m, 4H), 3.89 – 3.63 (m, 6H), 3.56 (s, 2H), 3.35 (dd, *J* = 12.5, 5.3 Hz, 2H), 2.67 – 2.44 (m, 2H), 2.39 – 2.32 (m, 1H), 2.26 (t, *J* = 5.9 Hz, 2H), 2.17 – 2.02 (m, 2H), 1.76 (d, *J* = 8.7 Hz, 2H). **$^{13}$C NMR (126 MHz, MeOD)** δ 170.00, 166.25 (d, *J* = 2.0 Hz), 165.29 (d, *J* = 1.4 Hz), 165.21, 163.24, 155.59 (d, *J* = 2.3 Hz), 152.43 (d, *J* = 257.7 Hz), 150.14 (d, *J* = 12.0 Hz), 146.86 (d, *J* = 15.6 Hz), 145.25 (d, *J* = 6.6 Hz), 136.49, 135.50, 134.52 (d, *J* = 5.2 Hz), 134.22, 131.55 (d, *J* = 9.3 Hz), 127.09, 124.14, 119.81, 117.04 (d, *J* = 26.2 Hz), 113.09, 112.43, 105.41 (d, *J* = 16.8 Hz), 98.52, 97.12, 89.93, 77.14, 76.22, 73.03, 61.13 (d, *J* = 19.9 Hz), 60.52, 58.60, 56.03, 42.82 (d, *J* = 20.5 Hz), 36.85 (d, *J* = 1.9 Hz), 26.19, 25.66, 25.56. **$^{19}$F NMR (470 MHz, MeOD)** δ -111.62, -111.64, -139.89, -173.97, -174.02. **HRMS (ESI):** *m/z* Calc. for $C_{37}H_{35}F_3N_8O_2$ $[M+H]^+$: 681.2908, found: 681.2929.

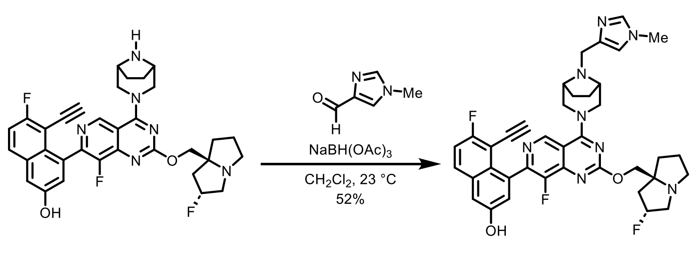


MRTX1133 **2**

**Compound 2.** A 3-mL vial was charged with MRTX1133 (3.0 mg, 1 equiv., 5.0 μmol), 1-methyl-5-imidazolecarboxaldehyde (1.1 mg, 2.0 equiv., 10 μmol), dichloromethane (0.50 mL) and a magnetic stir bar. Sodium triacetoxyborohydride (2.1 mg, 2.0 equiv., 10 μmol) was added to the stirred solution at 23 ºC. The reaction solution was stirred at 23 ºC for 16 h. The reaction mixture was concentrated under reduced pressure to afford the crude. The crude product was purified by prepHPLC (CombiFlash EZprep, C18 20x150mm column, 5–100% acetonitrile in water, 0.1% formic acid, 30 min) to afford imidazole **2** as pale orange solid 1.8 mg, 52% yield. **$^1$H NMR (500 MHz, MeOD)** δ 9.08 (s, 1H), 7.87 (dd, *J* = 9.1, 5.7 Hz, 1H), 7.84 (s, 1H), 7.36 (d, *J* = 2.6 Hz, 1H), 7.33 (t, *J* = 9.0 Hz, 1H), 7.25 (d, *J* = 1.4 Hz, 1H), 7.21 (d, *J* = 2.6 Hz, 1H), 5.70 – 5.43 (m, 1H), 4.77 – 4.55 (m, 4H), 4.03 – 3.74 (m, 10H), 3.73 – 3.62 (m, 2H), 3.46 – 3.33 (m, 2H), 2.72 – 2.49 (m, 2H), 2.44 – 2.37 (m, 1H), 2.37 – 2.26 (m, 2H), 2.25 – 2.05 (m, 3H), 1.86 – 1.73 (m, 2H); **$^{19}$F NMR (470 MHz, MeOD)** δ -111.61, -111.63, -139.83, -174.03, -174.08. **HRMS (ESI):** *m/z* Calc. for $C_{38}H_{37}F_3N_8O_2$ $[M+H]^+$: 695.3064, found: 695.3089.

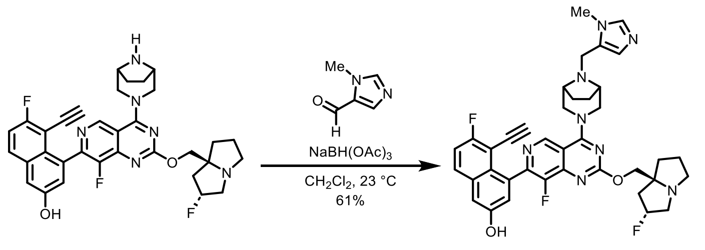


MRTX1133 3

**Compound 3.** A 3-mL vial was charged with MRTX1133 (3.0 mg, 1 equiv., 5.0 μmol), 1-methyl-1H-imidazole-4-carbaldehyde (1.1 mg, 2.0 equiv., 10 μmol), dichloromethane (0.50 mL) and a magnetic stir bar. Sodium triacetoxyborohydride (2.1 mg, 2.0 equiv, 10 μmol) was added to the stirred solution at 23 ºC. The reaction solution was stirred at 23 ºC for 16 h. The reaction mixture was concentrated under reduced pressure to afford the crude. The crude product was purified by prepHPLC (CombiFlash EZprep, C18 20x150mm column, 5–100% acetonitrile in water, 0.1% formic acid, 30 min) to afford imidazole **2** as pale orange solid 2.1 mg, 61% yield. **$^1$H NMR (500 MHz, MeOD)** δ 9.06 (s, 1H), 8.02 (s, 1H), 7.87 (dd, *J* = 9.2, 5.7 Hz, 1H), 7.41

– 7.28 (m, 2H), 7.21 (d, $J$ = 2.6 Hz, 1H), 7.10 (s, 1H), 5.65 – 5.41 (m, 1H), 4.75 – 4.51 (m, 4H), 4.09 – 3.73 (m, 8H), 3.68 (s, 2H), 3.52 – 3.37 (m, 4H), 2.75 – 2.51 (m, 2H), 2.44 – 2.26 (m, 3H), 2.15 – 2.07 (m, 3H), 1.78 – 1.66 (m, 2H). **$^{19}$F NMR (470 MHz, MeOD)** δ -111.60, -111.62, -140.01, -174.03, -174.08. **HRMS (ESI):** *m/z* Calc. for $C_{38}H_{37}F_3N_8O_2$ $[M+H]^+$: 695.3064, found: 695.3095.

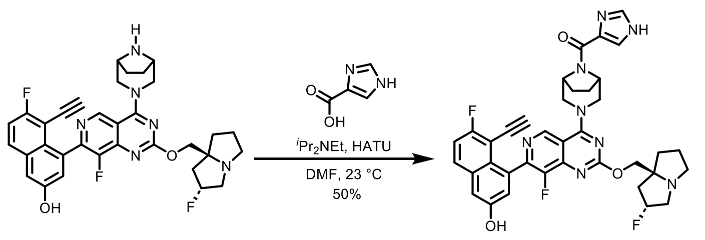


MRTX1133 **4**

**Compound 4.** A 3-mL vial was charged with 4-Imidazolecarboxylic acid (1.0 mg, 1.1 equiv., 9,2 μmol), dimethylformamide (0.55 mL) and a magnetic stir bar. N,N-Diisopropylethylamine (1.2 mg, 1.6 μL, 1.1 equiv., 9.2 μmol) and O-(7-Azabenzotriazol-1-yl)-N,N,N',N'-tetramethyluronium hexafluorophosphate (3.5 mg, 1.1 eq, 9.2 μmol) was added to the stirred solution at 23 ºC. The reaction solution was stirred at 23 ºC for 10 min. Then MRTX1133 (5.0 mg, 1.0 equiv., 8.3 μmol) was added to the stirred solution at 23 ºC. The reaction solution was stirred at 23 ºC for 16 h. The reaction mixture was concentrated under air blow to afford the crude. The crude product was purified by prepHPLC (CombiFlash EZprep, C18 20x150mm column, 5–100% acetonitrile in water, 0.1% formic acid, 30 min) and then purified by silica gel (RediSep® Silver Silica Gel Disposable Flash Columns 4 grams from Teledyne ISCO, 0–30% methanol in dichloromethane, 20 min) to afford imidazole **4** as white solid 2.9 mg, 50% yield. **$^1$H NMR** (500 MHz, MeOD) δ 9.11 (s, 1H), 7.88 (dd, $J$ = 9.2, 5.7 Hz, 1H), 7.77 (dd, $J$ = 13.5, 1.2 Hz, 2H), 7.39 – 7.30 (m, 2H), 7.22 (d, $J$ = 2.6 Hz, 1H), 5.49 (s, 3H), 4.72 – 4.56 (m, 2H),

4.11 – 3.68 (m, 5H), 3.35 (s, 4H), 2.75 – 1.96 (m, 10H). **$^{19}$F NMR (470 MHz, MeOD)** δ -111.44, -111.50, -139.71, -173.72, -173.77. **HRMS (ESI):** *m/z* Calc. for $C_{37}H_{33}F_3N_8O_3$ [M+H]$^+$: 695.2700, found: 695.2733.

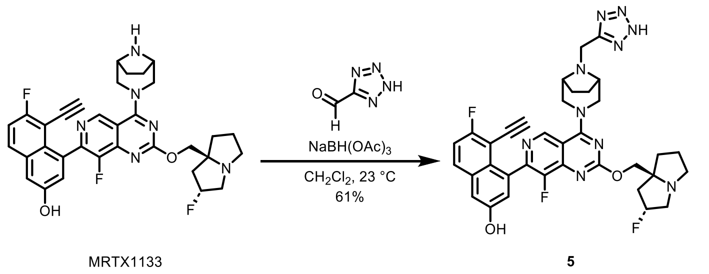


**Compound 5.** A 3-mL vial was charged with MRTX1133 (3.0 mg, 1 equiv., 5.0 μmol), 2H-tetrazole-5-carbaldehyde (2.4 mg, 5.0 equiv., 25 μmol), dichloromethane (0.5 mL) and a magnetic stir bar. Sodium triacetoxyborohydride (5.3 mg, 5.0 eq, 25 μmol) was added to the stirred solution at 23 ºC. The reaction solution was stirred at 23 ºC for 16 h. The reaction mixture was concentrated under reduced pressure to afford the crude. The crude product was purified by prepHPLC (CombiFlash EZprep, C18 20x150mm column, 5–100% acetonitrile in water, 0.1% formic acid, 30 min) to afford tetrazole **5** as pale orange solid 2.2 mg, 61% yield. **$^1$H NMR (500 MHz, $CDCl_3$)** δ 9.08 (s, 1H), 7.87 (dd, *J* = 9.2, 5.6 Hz, 1H), 7.41 – 7.29 (m, 2H), 7.21 (d, *J* = 2.6 Hz, 1H), 4.72 – 4.55 (m, 4H), 4.05 – 3.79 (m, 7H), 3.67 – 3.56 (m, 2H), 3.50 – 3.40 (m, 1H), 3.36 – 3.32 (m, 1H), 2.74 – 2.49 (m, 2H), 2.43 – 2.28 (m, 3H), 2.21 – 2.11 (m, 4H), 1.87 – 1.70 (m, 2H). **$^{19}$F NMR (470 MHz, MeOD)** δ -111.69, -111.70, -139.98, -173.88, -173.93. **HRMS (ESI):** *m/z* Calc. for $C_{35}H_{33}F_3N_{10}O_2$ [M+H]$^+$: 683.2813, found: 683.2846.

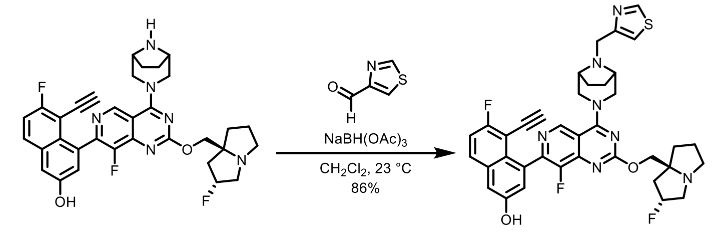


MRTX1133 6

**Compound 6.** A 3-mL vial was charged with MRTX1133 (3.0 mg, 1 equiv., 5.0 µmol), thiazole-4-carboxyaldehyde (2.8 mg, 5.0 equiv., 25 µmol), dichloromethane (0.50 mL) and a magnetic stir bar. Sodium triacetoxyborohydride (5.3 mg, 5.0 equiv., 25 µmol) was added to the stirred solution at 23 ºC. The reaction solution was stirred at 23 ºC for 16 h. The reaction mixture was concentrated under reduced pressure to afford the crude. The crude product was purified by prepHPLC (CombiFlash EZprep, C18 20x150mm column, 5–100% acetonitrile in water, 0.1% formic acid, 30 min) to afford thiazole **6** as pale orange solid 3.0 mg, 86% yield. **$^{1}$H NMR (500 MHz, MeOD)** δ 9.08 (s, 1H), 9.03 (d, *J* = 2.1 Hz, 1H), 7.87 (dd, *J* = 9.2, 5.7 Hz, 1H), 7.62 (d, *J* = 2.1 Hz, 1H), 7.36 (d, *J* = 2.6 Hz, 1H), 7.33 (t, *J* = 8.9 Hz, 1H), 7.21 (d, *J* = 2.5 Hz, 1H), 5.53 (d, *J* = 52.0 Hz, 1H), 4.61 (dtd, *J* = 19.8, 11.9, 6.3 Hz, 4H), 3.96 – 3.70 (m, 7H), 3.54 (d, *J* = 7.6 Hz, 2H), 3.44 – 3.33 (m, 2H), 2.70 – 2.49 (m, 2H), 2.39 (t, *J* = 9.0 Hz, 1H), 2.28 (dq, *J* = 12.2, 6.8 Hz, 3H), 2.21 – 2.06 (m, 3H), 1.75 (d, *J* = 8.6 Hz, 2H). **$^{19}$F NMR (470 MHz, MeOD)** δ -111.61, -111.63, -139.98, -174.00, -174.05. **HRMS (ESI):** *m/z* Calc. for $C_{14}H_{13}BrClNO_2S$ $[M+H]^+$: 698.2520, found: 698.2529.

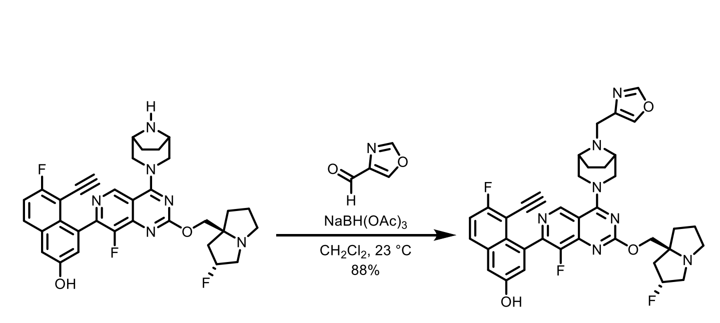


MRTX1133 7

**Compound 7.** A 3-mL vial was charged with MRTX1133 (3.0 mg, 1 equiv., 5.0 μmol), 4-oxazolecarboxaldehyde (2.4 mg, 5.0 equiv., 25 μmol), dichloromethane (0.50 mL) and a magnetic stir bar. Sodium triacetoxyborohydride (5.3 mg, 5.0 equiv., 25 μmol) was added to the stirred solution at 23 ºC. The reaction solution was stirred at 23 ºC for 16 h. The reaction mixture was concentrated under reduced pressure to afford the crude. The crude product was purified by silica gel (RediSep® Silver Silica Gel Disposable Flash Columns 4 grams from Teledyne ISCO, 0–30% methanol in dichloromethane, 20 min) to afford oxazole **7** as pale orange solid 3.0 mg, 88% yield. **$^1$H NMR (500 MHz, MeOD)** δ 9.06 (s, 1H), 8.22 (s, 1H), 7.95 (s, 1H), 7.87 (dd, *J* = 9.1, 5.7 Hz, 1H), 7.36 (d, *J* = 2.7 Hz, 1H), 7.33 (t, *J* = 9.0 Hz, 1H), 7.21 (d, *J* = 2.5 Hz, 1H), 5.46 (d, *J* = 52.8 Hz, 1H), 4.61 (t, *J* = 13.3 Hz, 2H), 4.56 – 4.40 (m, 2H), 3.86 – 3.48 (m, 9H), 3.33 (d, *J* = 8.8 Hz, 1H), 3.27 (dd, *J* = 10.2, 5.7 Hz, 1H), 2.59 – 2.38 (m, 2H), 2.34 – 2.28 (m, 1H), 2.23 – 2.15 (m, 2H), 2.15 – 2.00 (m, 3H), 1.73 (d, *J* = 9.0 Hz, 2H). **$^{19}$F NMR (470 MHz, MeOD)** δ -111.65, -111.66, -139.91, -139.92, -173.93, -173.98. **HRMS (ESI):** *m/z* Calc. for $C_{14}H_{13}BrClNO_3$ $[M+H]^+$: 682.2748, found: 682.2758.

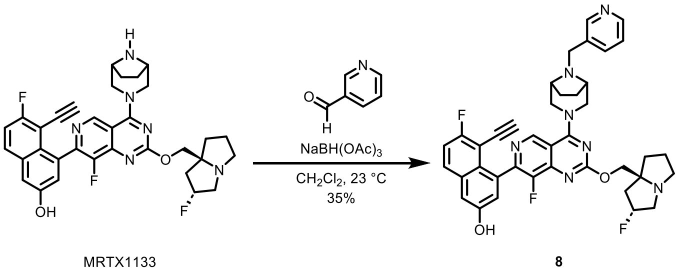


**Compound 8.** A 3-mL vial was charged with MRTX1133 (3.0 mg, 1 equiv., 5.0 μmol), nicotinaldehyde (1.6 mg, 3.0 equiv., 15 μmol), dichloromethane (0.50 mL) and a magnetic stir bar. Sodium triacetoxyborohydride (3.2 mg, 3.0 equiv, 15 μmol) was added to the stirred solution at 23 ºC. The reaction solution was stirred at 23 ºC for 16 h. The reaction mixture was

concentrated under reduced pressure to afford the crude. The crude product was purified by prepHPLC (CombiFlash EZprep, C18 20x150mm column, 5–100% acetonitrile in water, 0.1% formic acid, 30 min) to afford pyridine **8** as pale yellow solid 1.2 mg, 35% yield. **$^1$H NMR (500 MHz, MeOD)** δ 9.07 (s, 1H), 8.71 – 8.46 (m, 2H), 8.00 (d, *J* = 7.8 Hz, 1H), 7.87 (dd, *J* = 9.1, 5.7 Hz, 1H), 7.55 – 7.42 (m, 1H), 7.39 – 7.28 (m, 2H), 7.20 (d, *J* = 2.6 Hz, 1H), 5.58 – 5.40 (m, 1H), 4.69 – 4.40 (m, 4H), 3.90 – 3.57 (m, 7H), 3.47 – 3.31 (m, 4H), 2.69 – 2.42 (m, 2H), 2.38 – 2.00 (m, 6H), 1.75 (d, *J* = 8.6 Hz, 2H). **$^{19}$F NMR (470 MHz, MeOD)** δ -111.64, -111.66, -139.96, -173.96, -174.02. **HRMS (ESI):** *m/z* Calc. for $C_{39}H_{36}F_3N_7O_2$ $[M+H]^+$: 692.2955, found: 692.2981.

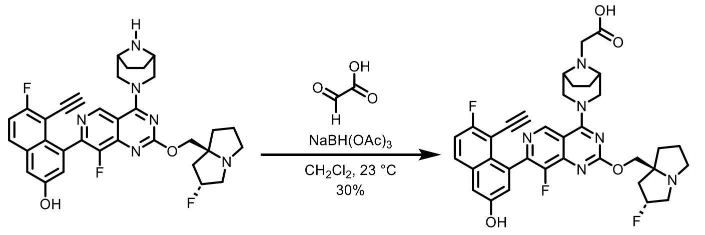


MRTX1133 **9**

**Compound 9.** A 3-mL vial was charged with MRTX1133 (3.0 mg, 1 equiv., 5.0 µmol), glyoxylic acid (50% in water, 2.8 mg, 3.0 equiv., 15 µmol), dichloromethane (0.50 mL) and a magnetic stir bar. Sodium triacetoxyborohydride (3.2 mg, 3.0 equiv., 15 µmol) was added to the stirred solution at 23 ºC. The reaction solution was stirred at 23 ºC for 16 h. The reaction mixture was concentrated under reduced pressure to afford the crude. The crude product was purified by prepHPLC (CombiFlash EZprep, C18 20x150mm column, 5–100% acetonitrile in water, 0.1% formic acid, 30 min) to afford carboxylic acid **9** as pale orange solid 0.9 mg, 30% yield. **$^1$H NMR (500 MHz, MeOD)** δ 9.11 (s, 1H), 7.88 (dd, J = 9.2, 5.7 Hz, 1H), 7.42 – 7.28 (m, 2H), 7.21 (d, J = 2.6 Hz, 1H), 5.54 (d, J = 51.4 Hz, 1H), 4.81 (d, J = 14.1 Hz, 2H), 4.73 –

4.56 (m, 2H), 4.09 (t, J = 14.8 Hz, 4H), 3.91 – 3.71 (m, 3H), 3.60 (s, 2H), 3.49 – 3.34 (m, 2H), 2.78 – 2.47 (m, 2H), 2.38 – 2.06 (m, 6H), 1.99 (d, J = 9.1 Hz, 2H). **$^{19}$F NMR (470 MHz, MeOD)** δ -111.61, -111.63, -139.44, -174.05, -174.09. **HRMS (ESI):** *m/z* Calc. for $C_{35}H_{33}F_3N_6O_4$ $[M+H]^+$: 659.2588, found: 659.2612.

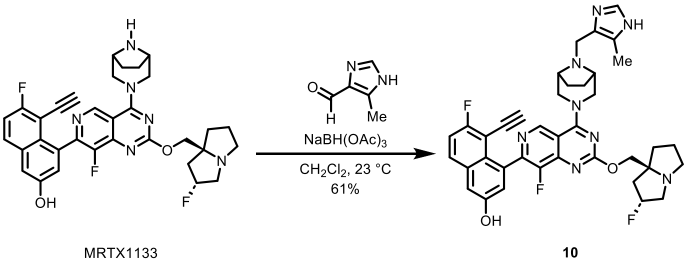

**Compound 10.** A 3-mL vial was charged with MRTX1133 (3.0 mg, 1 equiv., 5.0 μmol), 5-methyl-1H-imidazole-4-carbaldehyde (1.1 mg, 2.0 equiv., 10 μmol), dichloromethane (0.5 mL) and a magnetic stir bar. Sodium triacetoxyborohydride (2.1 mg, 2.0 equiv., 10 μmol) was added to the stirred solution at 23 ºC. The reaction solution was stirred at 23 ºC for 16 h. The reaction mixture was concentrated under reduced pressure to afford the crude. The crude product was purified by prepHPLC (CombiFlash EZprep, C18 20x150mm column, 5–100% acetonitrile in water, 0.1% formic acid, 30 min) to afford imidazole **10** as pale yellow solid 2.1 mg, 61% yield. **$^{1}$H NMR (500 MHz, MeOD)** δ 9.07 (s, 1H), 8.10 (s, 1H), 7.89 (dd, *J* = 9.2, 5.7 Hz, 1H), 7.47 – 7.30 (m, 2H), 7.23 (d, *J* = 2.6 Hz, 1H), 5.73 – 5.33 (m, 1H), 4.74 – 4.46 (m, 4H), 3.97 – 3.70 (m, 5H), 3.69 (s, 2H), 3.61 – 3.45 (m, 2H), 3.45 – 3.35 (m, 2H), 2.75 – 2.46 (m, 3H), 2.35 (s, 3H), 2.32 – 2.11 (m, 5H), 1.87 – 1.74 (m, 2H). **$^{19}$F NMR (470 MHz, MeOD)** δ -111.62, -111.64, -139.86, -173.96, -174.01. **HRMS (ESI):** *m/z* Calc. for $C_{38}H_{37}F_3N_8O_2$ $[M+H]^+$: 695.3064, found: 695.3097.

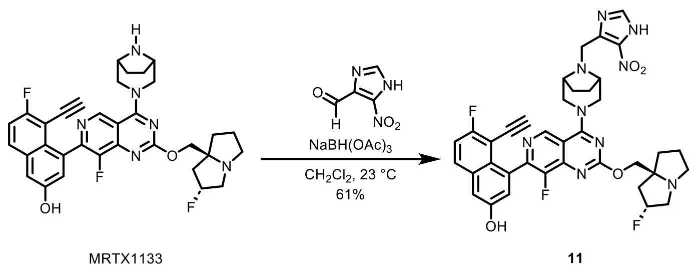


**Compound 11.** A 3-mL vial was charged with MRTX1133 (3.0 mg, 1 equiv., 5.0 μmol), 5-nitro-1H-imidazole-4-carbaldehyde (1.4 mg, 2.0 equiv., 10 μmol), dichloromethane (0.5 mL) and a magnetic stir bar. Sodium triacetoxyborohydride (2.1 mg, 2.0 equiv., 10 μmol) was added to the stirred solution at 23 ºC. The reaction solution was stirred at 23 ºC for 16 h. The reaction mixture was concentrated under reduced pressure to afford the crude. The crude product was purified by prepHPLC (CombiFlash EZprep, C18 20x150mm column, 5–100% acetonitrile in water, 0.1% formic acid, 30 min) to afford imidazole **11** as pale yellow solid 2.2 mg, 61% yield. **$^1$H NMR (500 MHz, MeOD)** δ 9.07 (s, 1H), 7.87 (dd, *J* = 9.2, 5.7 Hz, 1H), 7.75 (s, 1H), 7.44 – 7.30 (m, 2H), 7.21 (d, *J* = 2.6 Hz, 1H), 5.73 – 5.41 (m, 1H), 4.71 – 4.57 (m, 4H), 4.09 (s, 2H), 4.01 – 3.75 (m, 5H), 3.56 – 3.39 (m, 3H), 3.33 (dd, *J* = 4.5, 0.9 Hz, 1H), 2.78 – 2.49 (m, 2H), 2.43 – 2.25 (m, 3H), 2.15 (t, *J* = 7.1 Hz, 3H), 1.79 – 1.73 (m, 2H). **$^{19}$F NMR (470 MHz, MeOD)** δ -111.58, -111.60, -139.95, -174.04, -174.09. **HRMS (ESI):** *m/z* Calc. for $C_{37}H_{34}F_3N_9O_4$ $[M+H]^+$: 726.2759, found: 726.2790.

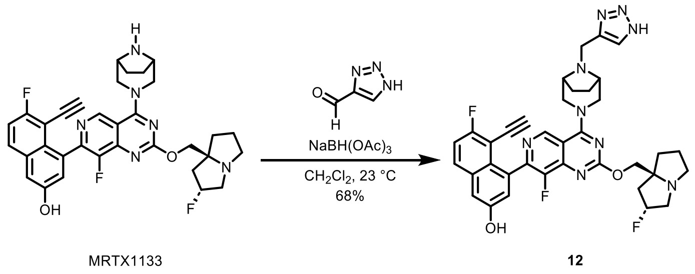


**Compound 12.** A 3-mL vial was charged with MRTX1133 (3.0 mg, 1 equiv., 5.0 μmol), 1H-

1,2,3-triazole-4-carbaldehyde (1.5 mg, 3.0 equiv., 15 μmol), dichloromethane (0.50 mL) and a magnetic stir bar. Sodium triacetoxyborohydride (3.2 mg, 3.0 equiv., 15 μmol) was added to the stirred solution at 23 ºC. The reaction solution was stirred at 23 ºC for 16 h. The reaction mixture was concentrated under reduced pressure to afford the crude. The crude product was purified by prepHPLC (CombiFlash EZprep, C18 20x150mm column, 5–100% acetonitrile in water, 0.1% formic acid, 30 min) to afford imidazole **12** as pale yellow solid 2.3 mg, 68% yield. **$^1$H NMR (500 MHz, MeOD)** δ 9.08 (s, 1H), 8.01 – 7.73 (m, 2H), 7.44 – 7.29 (m, 2H), 7.21 (d, *J* = 2.6 Hz, 1H), 5.72 – 5.31 (m, 1H), 4.75 – 4.47 (m, 4H), 4.07 – 3.75 (m, 7H), 3.61 – 3.51 (m, 2H), 3.47 – 3.33 (m, 2H), 2.77 – 2.49 (m, 2H), 2.47 – 2.24 (m, 3H), 2.22 – 2.07 (m, 3H), 1.81 – 1.73 (m, 2H). **$^{19}$F NMR (470 MHz, MeOD)** δ -111.60, -111.62, -139.98, -174.04, -174.09. **HRMS (ESI):** *m/z* Calc. for $C_{36}H_{34}F_3N_9O_2$ $[M+H]^+$: 682.2860, found: 682.2891.

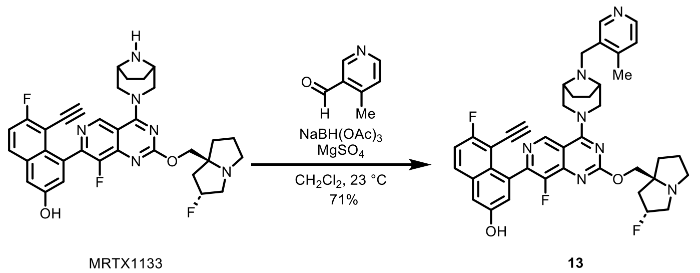


**Compound 13.** A 3-mL vial was charged with MRTX1133 (5.0 mg, 1 equiv., 8.3 μmol), 4-methylnicotinaldehyde (9.4 mg, 9.3 equiv., 77 μmol), dichloromethane (0.55 mL) and a magnetic stir bar. Anhydrous magnesium sulfate (20 mg, 20 equiv, 170 μmol) was added to the stirred solution at 23 ºC. Then sodium triacetoxyborohydride (7.1 mg, 4.0 equiv., 33 μmol) was added to the stirred solution at 23 ºC. The reaction solution was stirred at 23 ºC for 16 h. The reaction mixture was concentrated under reduced pressure to afford the crude. The crude product was purified by prepHPLC (CombiFlash EZprep, C18 20x150mm column, 5–100%

acetonitrile in water, 0.1% formic acid, 30 min) to afford pyridine **13** as pale orange solid 4.2 mg, 71% yield. $^{1}$**H NMR (500 MHz, MeOD)** δ 9.01 (s, 1H), 8.45 (s, 1H), 8.33 (d, *J* = 5.0 Hz, 1H), 7.86 (dd, *J* = 9.1, 5.7 Hz, 1H), 7.37 – 7.33 (m, 1H), 7.33 – 7.27 (m, 2H), 7.20 (d, *J* = 2.6 Hz, 1H), 5.37 (d, *J* = 53.4 Hz, 1H), 4.59 (q, *J* = 12.0 Hz, 2H), 4.36 (ddd, *J* = 36.7, 11.0, 7.1 Hz, 2H), 3.88 – 3.61 (m, 4H), 3.52 – 3.33 (m, 6H), 3.20 – 3.04 (m, 1H), 2.56 (s, 3H), 2.45 – 2.25 (m, 2H), 2.22 – 2.13 (m, 3H), 2.11 – 2.05 (m, 2H), 1.95 (s, 1H), 1.75 (d, *J* = 8.9 Hz, 2H). $^{19}$**F NMR (470 MHz, MeOD)** δ -111.68, -111.69, -139.75, -139.78, -173.74, -173.78. **HRMS (ESI):** *m/z* Calc. for $C_{40}H_{38}F_3N_7O_2$ $[M+H]^+$: 706.3112, found: 706.3144.

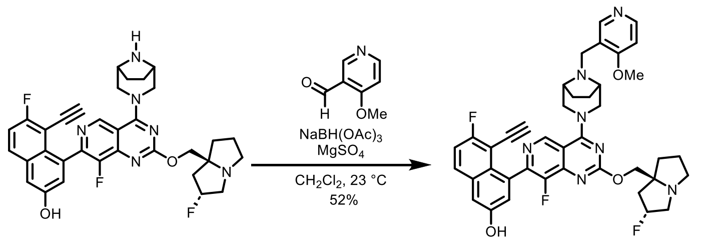


MRTX1133 **14**

**Compound 14.** A 3-mL vial was charged with MRTX1133 (5.0 mg, 1 equiv., 8.3 μmol), 4-methoxynicotinaldehyde (11 mg, 9.3 equiv., 77 μmol), dichloromethane (0.55 mL) and a magnetic stir bar. Anhydrous magnesium sulfate (20 mg, 20 eq, 170 μmol) was added to the stirred solution at 23 ºC. Then sodium triacetoxyborohydride (7.1 mg, 4.0 equiv., 33 μmol) was added to the stirred solution at 23 ºC. The reaction solution was stirred at 23 ºC for 16 h. The reaction mixture was concentrated under reduced pressure to afford the crude. The crude product was purified by prepHPLC (CombiFlash EZprep, C18 20x150mm column, 5–100% acetonitrile in water, 0.1% formic acid, 30 min) to afford pyridine **14** as pale orange solid 3.1 mg, 52% yield. $^{1}$**H NMR (500 MHz, MeOD)** δ 9.05 (s, 1H), 8.59 (s, 1H), 8.40 (d, *J* = 5.9 Hz, 1H), 7.85 (dd, *J* = 9.2, 5.7 Hz, 1H), 7.35 (d, *J* = 2.5 Hz, 1H), 7.32 (t, *J* = 8.9 Hz, 1H), 7.20 (d,

$J$ = 2.5 Hz, 1H), 7.11 (d, $J$ = 5.9 Hz, 1H), 5.63 – 5.33 (m, 1H), 4.69 – 4.42 (m, 4H), 3.97 (s, 3H), 3.89 – 3.67 (m, 6H), 3.60 (t, $J$ = 16.1 Hz, 1H), 3.48 (s, 2H), 3.35 (d, $J$ = 2.2 Hz, 1H), 3.27 (dd, $J$ = 10.1, 5.7 Hz, 1H), 2.61 – 2.39 (m, 2H), 2.32 (t, $J$ = 8.7 Hz, 1H), 2.23 – 2.02 (m, 5H), 1.75 (d, $J$ = 9.6 Hz, 2H). **$^{19}$F NMR (470 MHz, MeOD)** δ -111.63, -111.64, -139.81, -139.82, -173.79, -173.82. **HRMS (ESI):** *m/z* Calc. for $C_{40}H_{38}F_3N_7O_3$ $[M+H]^+$: 722.3061, found: 722.3088.

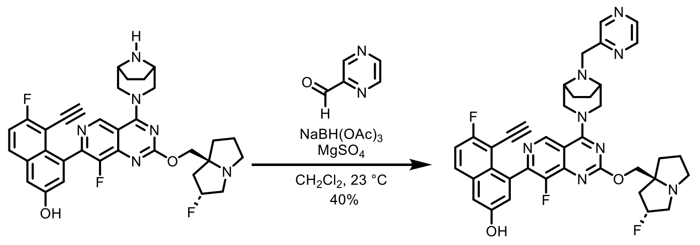


MRTX1133 **15**

**Compound 15.** A 3-mL vial was charged with MRTX1133 (8.0 mg, 1 equiv., 13.3 μmol), pyrazine-2-carbaldehyde (14 mg, 10 equiv., 133 μmol), dichloromethane (0.50 mL) and a magnetic stir bar. Anhydrous magnesium sulfate (31 mg, 20 equiv., 266 μmol) was added to the stirred solution at 23 ºC. Then sodium triacetoxyborohydride (28 mg, 10 equiv., 133 μmol) was added to the stirred solution at 23 ºC. The reaction solution was stirred at 23 ºC for 16 h. The reaction mixture was concentrated under reduced pressure to afford the crude. The crude product was purified by prepHPLC (CombiFlash EZprep, C18 20x150mm column, 5–100% acetonitrile in water, 0.1% formic acid, 30 min) to afford pyrazine **15** as pale orange solid 4.0 mg, 40% yield. **$^1$H NMR (500 MHz, MeOD)** δ 9.09 (s, 1H), 8.91 (d, $J$ = 1.6 Hz, 1H), 8.60 (dd, $J$ = 2.7, 1.5 Hz, 1H), 8.55 (d, $J$ = 2.7 Hz, 1H), 7.87 (dd, $J$ = 9.2, 5.7 Hz, 1H), 7.41 – 7.29 (m, 2H), 7.21 (d, $J$ = 2.5 Hz, 1H), 5.54 (d, $J$ = 51.9 Hz, 1H), 4.75 – 4.53 (m, 5H), 4.13 – 3.72 (m, 8H), 3.52 (s, 2H), 3.42 (ddd, $J$ = 16.0, 8.5, 3.8 Hz, 1H), 3.36 – 3.31 (m, 6H), 2.76 – 2.01 (m,

10H). **$^{19}$F NMR (470 MHz, MeOD)** δ -111.72, -111.73, -139.61, -139.65, -173.65, -173.70.

**HRMS (ESI):** *m/z* Calc. for $C_{38}H_{35}F_3N_8O_2$ $[M+H]^+$: 693.2908, found: 693.2937.

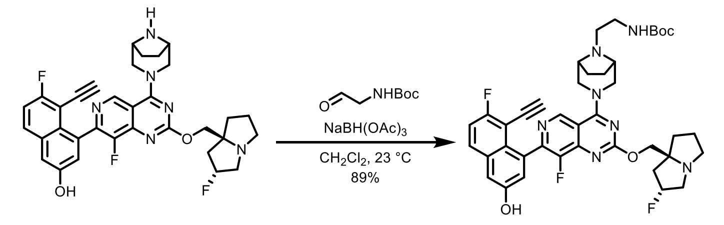


MRTX1133 **19**

**Compound 19.** A 3-mL vial was charged with MRTX1133 (30.0 mg, 1 equiv., 50.0 μmol), N-Boc-2-aminoacetaldehyde (15.9 mg, 2.00 equiv., 100 μmol), dichloromethane (1.00 mL) and a magnetic stir bar. Sodium triacetoxyborohydride (21.2 mg, 2.00 equiv., 100 μmol) was added to the stirred solution at 23 ºC. The reaction solution was stirred at 23 ºC for 16 h. The reaction mixture was concentrated under reduced pressure to afford the crude. The crude product was purified by prepHPLC (CombiFlash EZprep, C18 20x150mm column, 5–100% acetonitrile in water, 0.1% formic acid, 30 min) to afford **19** as pale yellow solid 33.0 mg, 89% yield. **$^1$H NMR (500 MHz, MeOD)** δ 9.07 (s, 1H), 7.85 (dd, *J* = 9.2, 5.6 Hz, 1H), 7.37 (d, *J* = 2.6 Hz, 1H), 7.31 (t, *J* = 8.7 Hz, 1H), 7.18 (s, 1H), 5.65 – 5.43 (m, 1H), 4.80 – 4.61 (m, 4H), 4.12 – 3.75 (m, 5H), 3.56 – 3.36 (m, 4H), 3.13 – 2.97 (m, 2H), 2.79 – 2.37 (m, 4H), 2.35 – 2.11 (m, 6H), 2.09 – 1.88 (m, 2H), 1.45 (s, 9H). **$^{19}$F NMR (470 MHz, MeOD)** δ -111.54, -111.56, -139.64, -173.93, -173.96. **HRMS (ESI):** *m/z* Calc. for $C_{40}H_{44}F_3N_7O_4$ $[M+H]^+$: 744.3480, found: 744.3509.

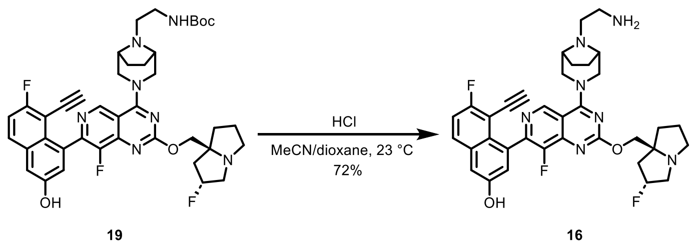


**Compound 16.** A 3-mL vial was charged with **16** (3.0 mg, 1 equiv., 5.0 µmol), acetonitrile (1.0 mL) and a magnetic stir bar. Hydrogen chloride (4 M in 1,4-dioxane, 0.20 mL) was added to the stirred solution at 23 ºC. The reaction solution was stirred at 23 ºC for 1 h. The reaction mixture was concentrated under reduced pressure to afford the crude. The crude product was purified by prepHPLC (CombiFlash EZprep, C18 20x150mm column, 5–100% acetonitrile in water, 0.1% formic acid, 30 min) to afford primary amine **16** as pale yellow solid 20.6 mg, 72% yield. **$^1$H NMR (500 MHz, MeOD)** δ 9.05 (s, 1H), 7.87 (dd, *J* = 9.2, 5.7 Hz, 1H), 7.37 (d, *J* = 2.6 Hz, 1H), 7.33 (t, *J* = 8.9 Hz, 1H), 7.22 (s, 1H), 5.68 – 5.43 (m, 1H), 4.70 – 4.54 (m, 4H), 4.08 – 3.71 (m, 5H), 3.53 – 3.33 (m, 4H), 3.09 (dd, *J* = 6.8, 4.7 Hz, 2H), 2.77 – 2.49 (m, 4H), 2.46 – 2.25 (m, 3H), 2.18 – 1.92 (m, 3H), 1.71 (d, *J* = 8.7 Hz, 2H). **$^{19}$F NMR (470 MHz, MeOD)** δ -111.57, -111.58, -139.82, -139.83, -173.89, -173.91. **HRMS (ESI):** *m/z* Calc. for $C_{35}H_{36}F_3N_7O_2$ $[M+H]^+$: 644.2955, found: 644.2986.

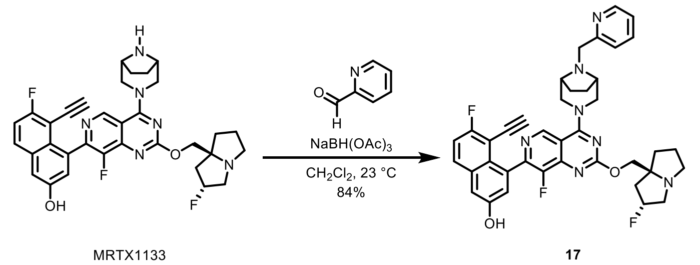


**Compound 17.** A 3-mL vial was charged with MRTX1133 (3.0 mg, 1 equiv., 5.0 µmol), 2-pyridinecarboxaldehyde (1.6 mg, 3.0 equiv., 15 µmol), dichloromethane (0.50 mL) and a magnetic stir bar. Sodium triacetoxyborohydride (3.2 mg, 3.0 equiv., 15 µmol) was added to

the stirred solution at 23 ºC. The reaction solution was stirred at 23 ºC for 16 h. The reaction mixture was concentrated under reduced pressure to afford the crude. The crude product was purified by silica gel (RediSep® Silver Silica Gel Disposable Flash Columns 4 grams from Teledyne ISCO, 0–30% methanol in dichloromethane, 20 min) to afford pyridine **17** as pale yellow solid 2.9 mg, 84% yield. **$^1$H NMR (500 MHz, MeOD)** δ 9.05 (s, 1H), 8.51 (dt, *J* = 4.9, 1.5 Hz, 1H), 7.87 (ddd, *J* = 10.6, 9.2, 6.7 Hz, 2H), 7.76 (d, *J* = 7.8 Hz, 1H), 7.39 – 7.29 (m, 3H), 7.21 (d, *J* = 2.5 Hz, 1H), 5.41 (d, *J* = 53.8 Hz, 1H), 4.66 – 4.56 (m, 2H), 4.54 – 4.39 (m, 2H), 3.84 (t, *J* = 10.9 Hz, 2H), 3.79 (s, 2H), 3.71 – 3.49 (m, 3H), 3.45 (d, *J* = 7.8 Hz, 2H), 3.34 (dd, *J* = 7.5, 1.0 Hz, 1H), 3.24 (td, *J* = 10.3, 5.7 Hz, 1H), 2.58 – 2.35 (m, 2H), 2.33 – 2.25 (m, 1H), 2.22 – 2.11 (m, 4H), 2.08 – 1.99 (m, 1H), 1.75 (d, *J* = 8.6 Hz, 2H). **$^{19}$F NMR (470 MHz, MeOD)** δ -111.64, -111.65, -139.85, -139.86, -173.86, -173.91. **HRMS (ESI):** *m/z* Calc. for $C_{39}H_{36}F_3N_7O_2$ $[M+H]^+$: 692.2955, found: 692.2955.

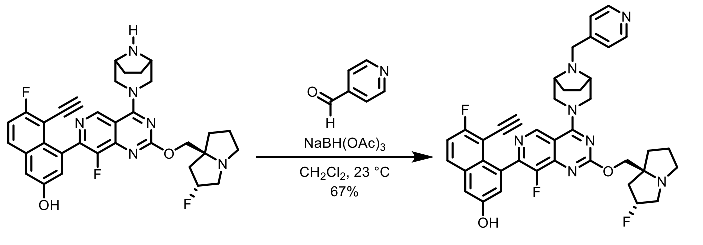


MRTX1133 18

**Compound 18.** A 3-mL vial was charged with MRTX1133 (3.0 mg, 1 equiv., 5.0 μmol), 4-pyridinecarboxaldehyde (1.6 mg, 3.0 equiv., 15 μmol), dichloromethane (0.50 mL) and a magnetic stir bar. Sodium triacetoxyborohydride (3.2 mg, 3.0 equiv., 15 μmol) was added to the stirred solution at 23 ºC. The reaction solution was stirred at 23 ºC for 16 h. The reaction mixture was concentrated under reduced pressure to afford the crude. The crude product was purified by prepHPLC (CombiFlash EZprep, C18 20x150mm column, 5–100% acetonitrile in water, 0.1% formic acid, 30 min) to afford pyridine **18** as pale yellow solid 2.3 mg, 67% yield. **$^1$H NMR (500 MHz, MeOD)** δ 9.09 (s, 1H), 8.51 (d, *J* = 5.1 Hz, 2H), 7.87 (dd, *J* = 9.2, 5.7 Hz, 1H), 7.62 – 7.57 (m, 2H), 7.36 (d, *J* = 2.6 Hz, 1H), 7.33 (t, *J* = 8.9 Hz, 1H), 7.21 (d, *J* = 2.6 Hz, 1H), 5.51 (d, *J* = 52.3 Hz, 1H), 4.70 – 4.44 (m, 4H), 4.08 – 3.61 (m, 7H), 3.58 – 3.35 (m, 3H), 2.84 – 2.47 (m, 2H), 2.39 – 2.07 (m, 6H), 1.74 (d, *J* = 8.7 Hz, 2H). **$^{19}$F NMR (470 MHz, MeOD)** δ -111.64, -111.65, -139.85, -139.86, -173.86, -173.91. **HRMS (ESI):** *m/z* Calc. for $C_{39}H_{36}F_3N_7O_2$ $[M+H]^+$: 692.2955, found: 692.2963.

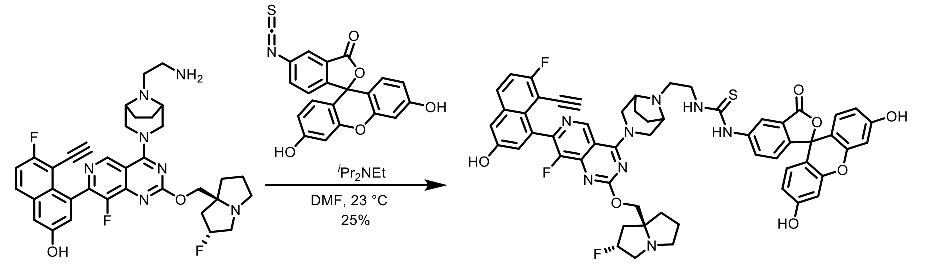


16 20

**Compound 20.** A 3-mL vial was charged with amine **16** (6.4 mg, 1 equiv., 10 μmol), FITC (7.8 mg, 2.0 equiv., 20 μmol), DMF (0.50 mL) and a magnetic stir bar. N,N-Diisopropylethylamine (3.9 mg, 5.2 μL, 3.0 equiv., 30 μmol) was added to the stirred solution at 23 ºC. The reaction solution was stirred at 23 ºC for 16 h. The reaction mixture was concentrated under reduced pressure to afford the crude. The crude product was purified by prepHPLC (CombiFlash EZprep, C18 20x150mm column, 5–100% acetonitrile in water, 0.1% formic acid, 30 min) to afford MRTX1133-FITC **20** as orange solid 2.6 mg, 25% yield. **$^1$H NMR (500 MHz, MeOD)** δ 9.05 (s, 1H), 8.18 (s, 1H), 7.87 (dd, *J* = 9.2, 5.7 Hz, 1H), 7.69 (s, 1H), 7.36 (d, *J* = 2.6 Hz, 1H), 7.32 (t, *J* = 8.9 Hz, 1H), 7.27 – 7.22 (m, 2H), 6.76 – 6.60 (m, 4H), 6.60 – 6.41 (m, 2H), 5.69 – 5.34 (m, 1H), 4.69 – 4.46 (m, 4H), 3.96 – 3.51 (m, 9H), 2.79 (d, *J* = 6.4 Hz, 2H), 2.65 –

2.19 (m, 5H), 2.06 (d, $J$ = 20.6 Hz, 3H), 1.74 (d, $J$ = 11.6 Hz, 2H). **$^{19}$F NMR (470 MHz, MeOD)** δ -111.64, -139.78, -139.86, -173.96. **HRMS (ESI):** *m/z* Calc. for $C_{56}H_{47}F_3N_8O_7S$ $[M+H]^+$: 1033.3313, found: 1033.3315.

#### Part 5. $^{1}H$ NMR, $^{19}F$ NMR and $^{13}C$ NMR Spectra.

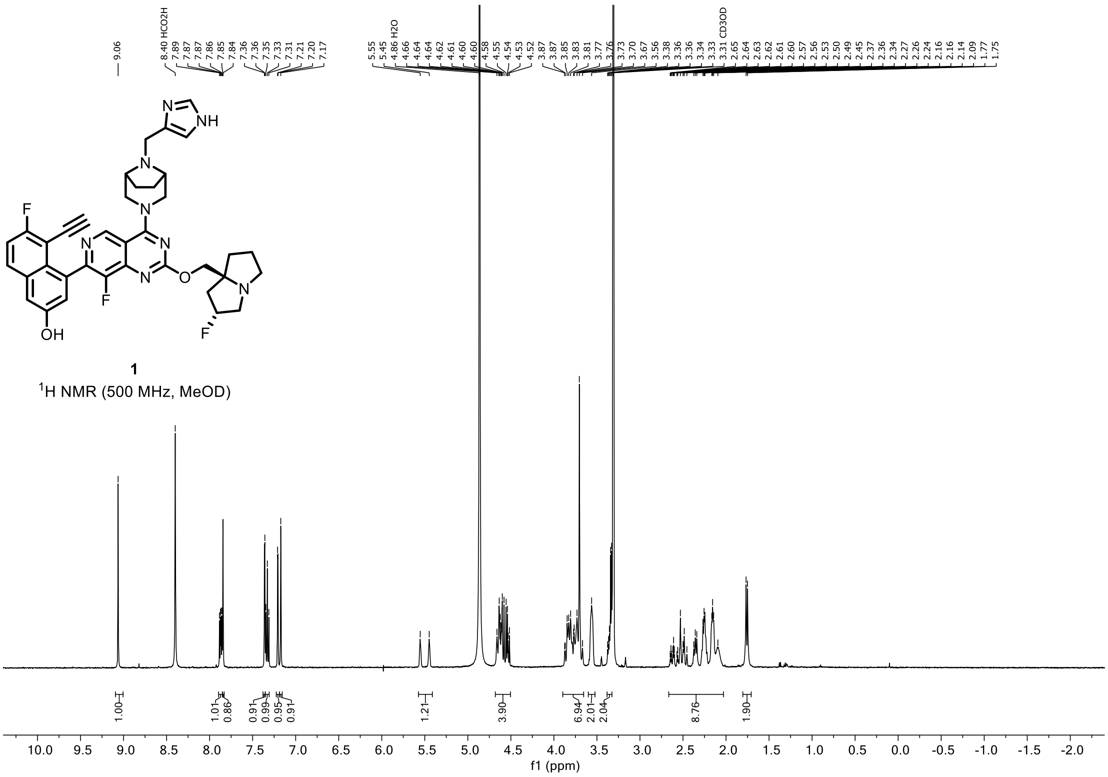


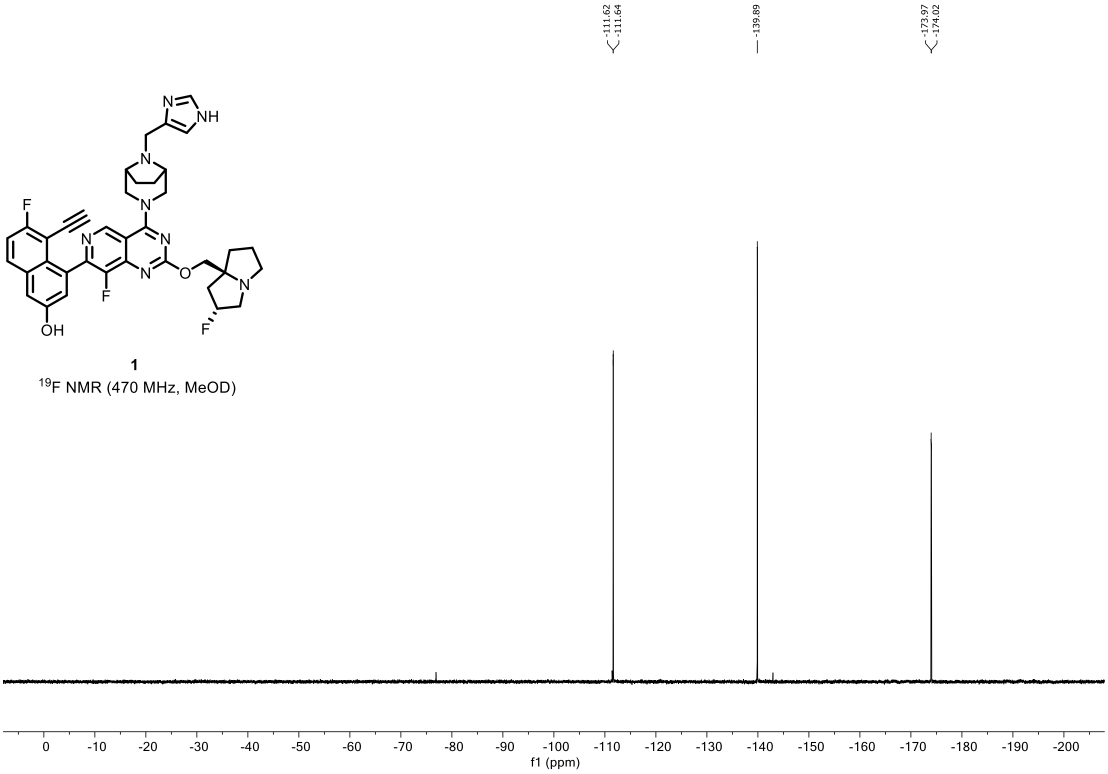

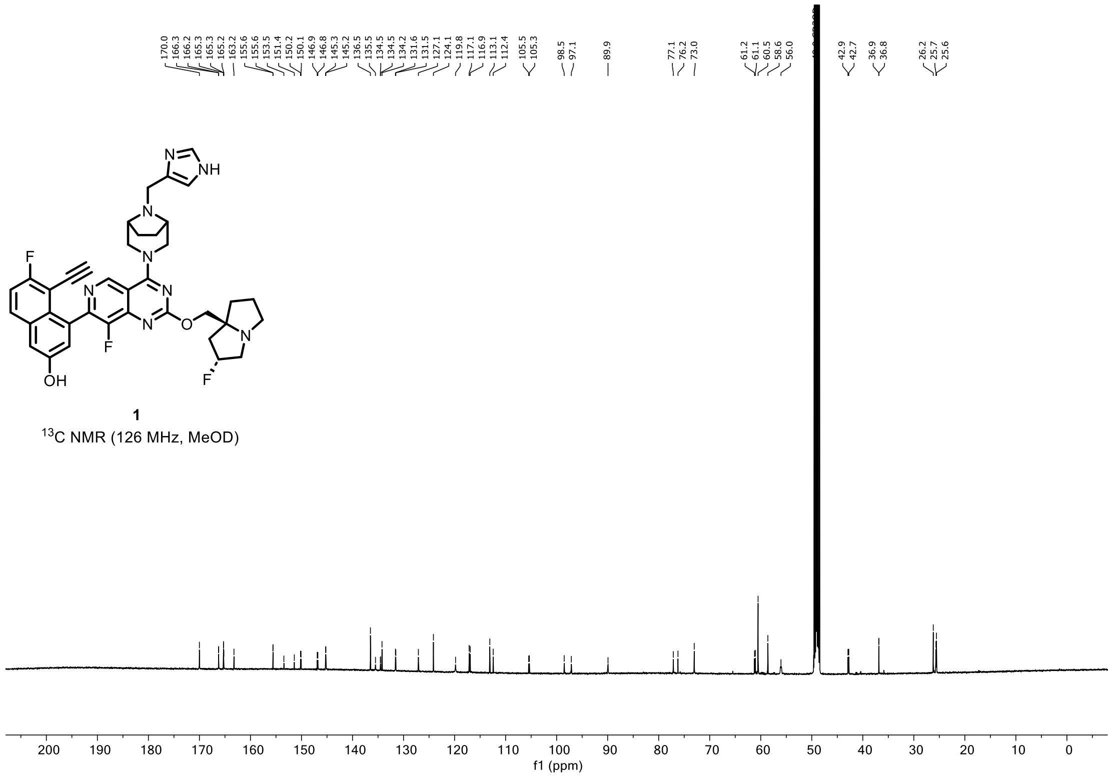


**1**
$^{13}$C NMR (126 MHz, MeOD)

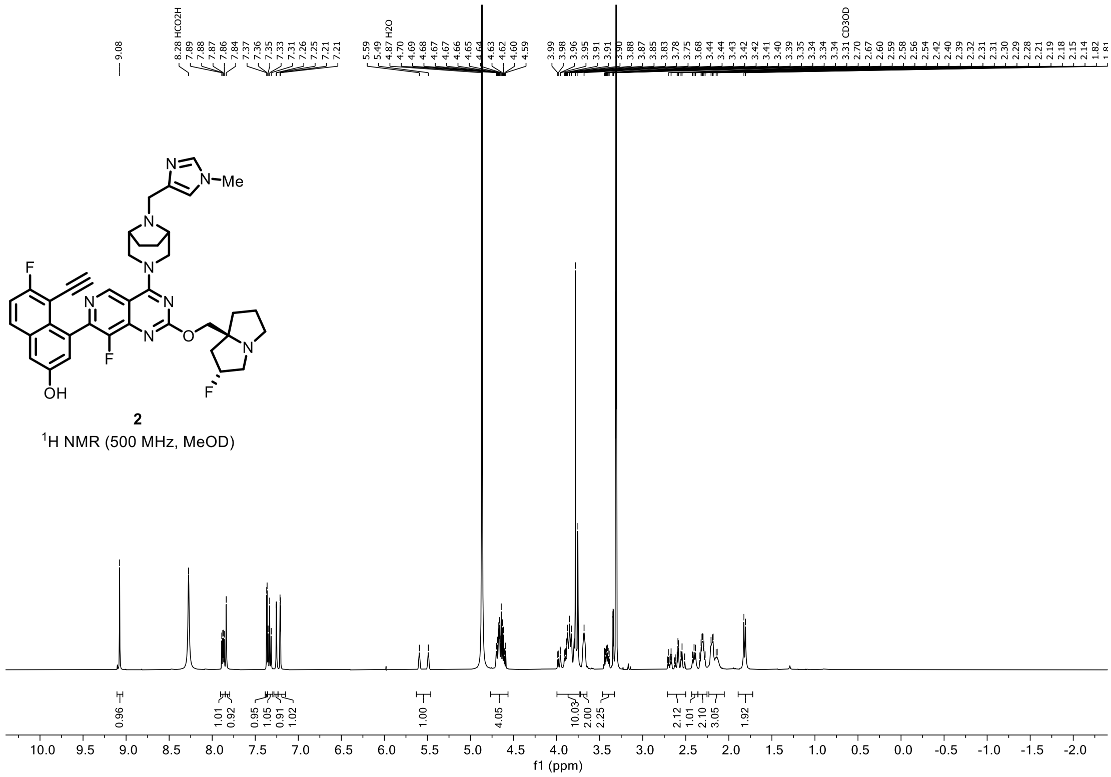
2
$^1$H NMR (500 MHz, MeOD)
f1 (ppm)

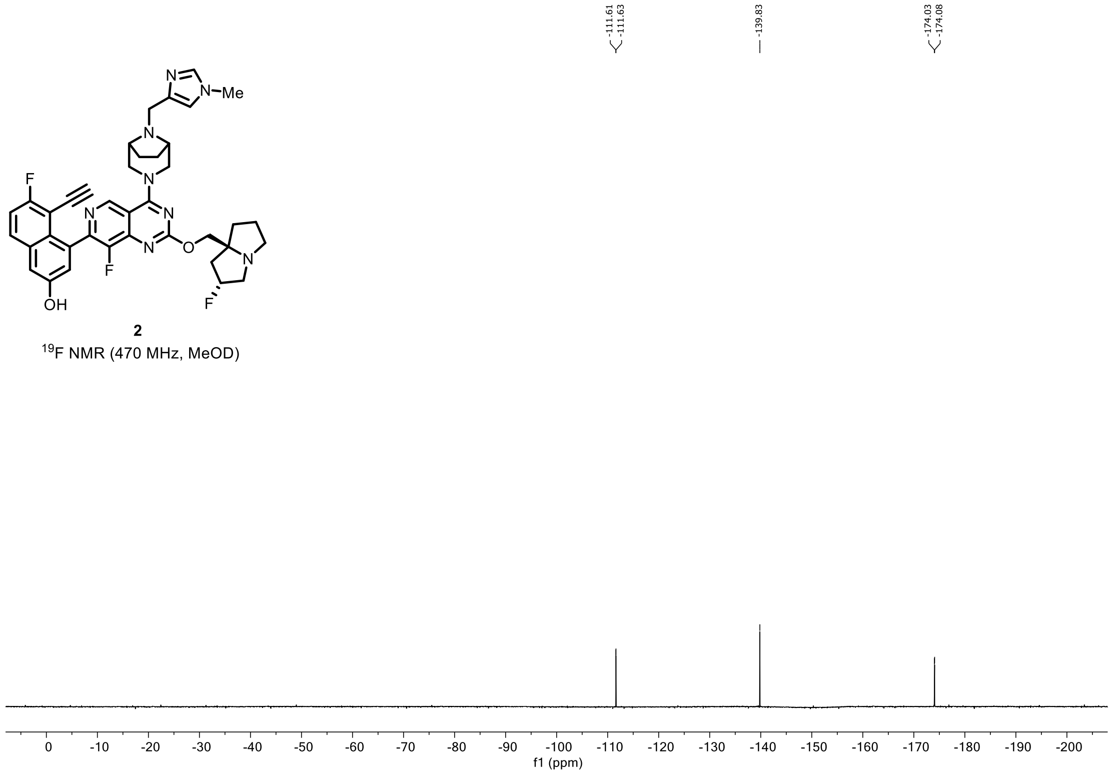
2
$^{19}$F NMR (470 MHz, MeOD)
f1 (ppm)

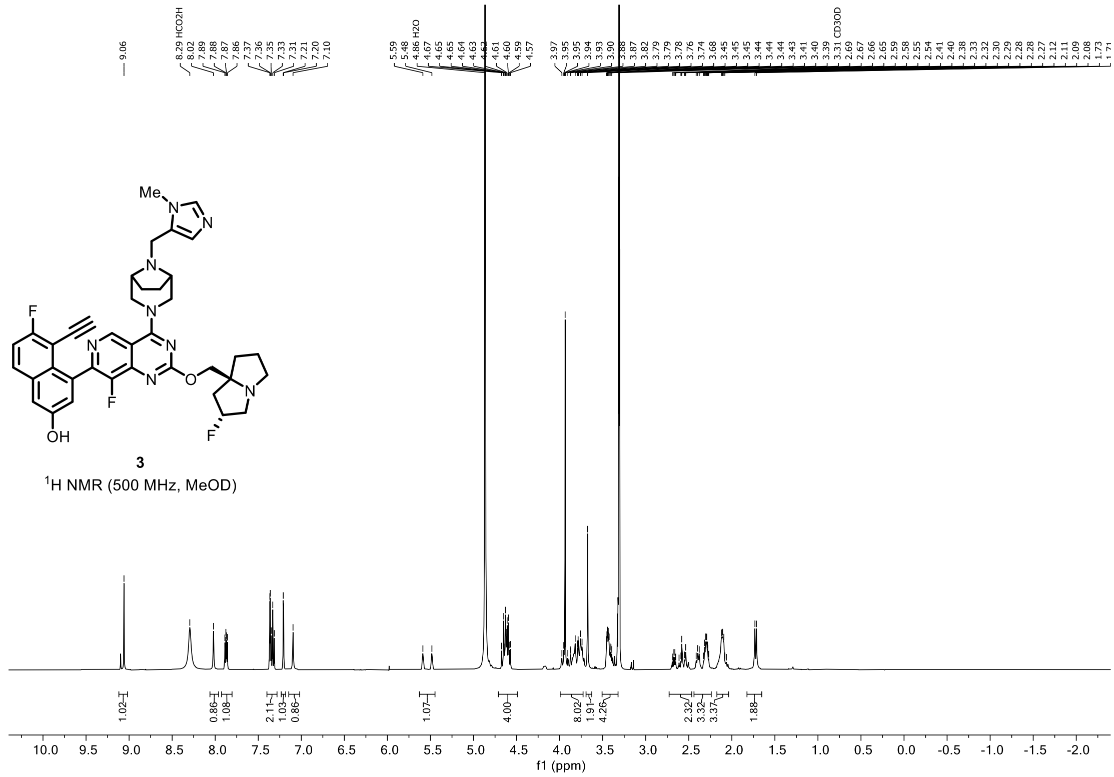


**3**

$^1$H NMR (500 MHz, MeOD)

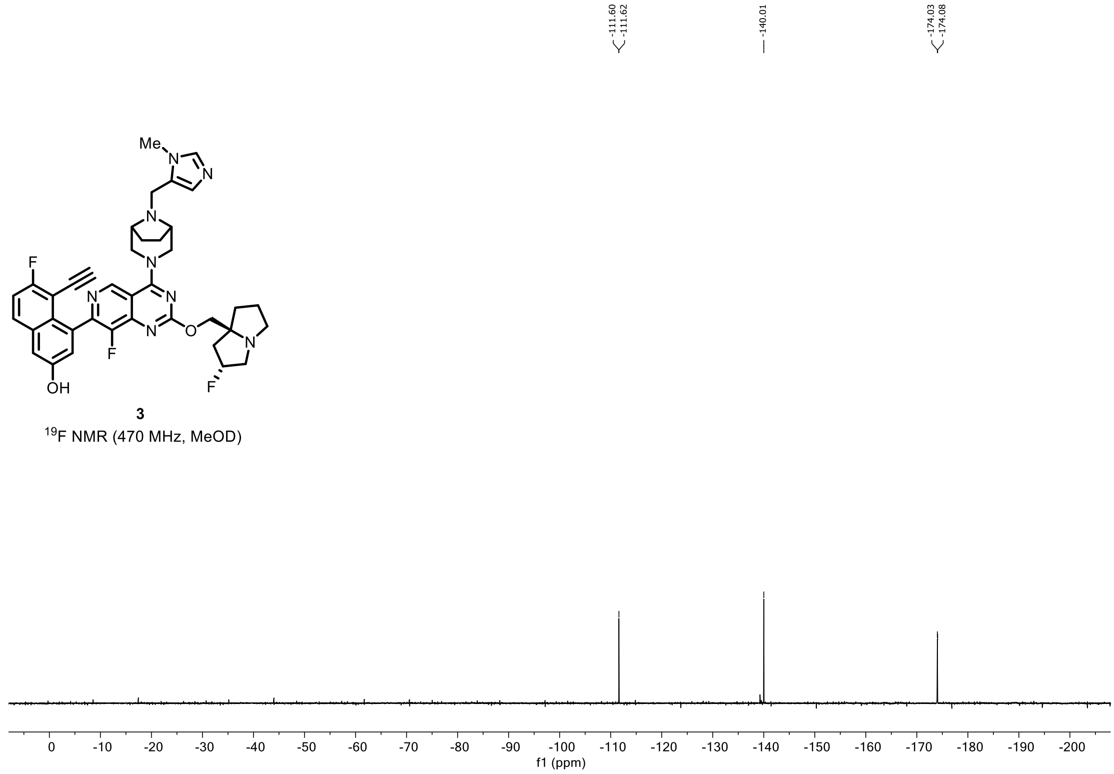


**3**

$^{19}$F NMR (470 MHz, MeOD)

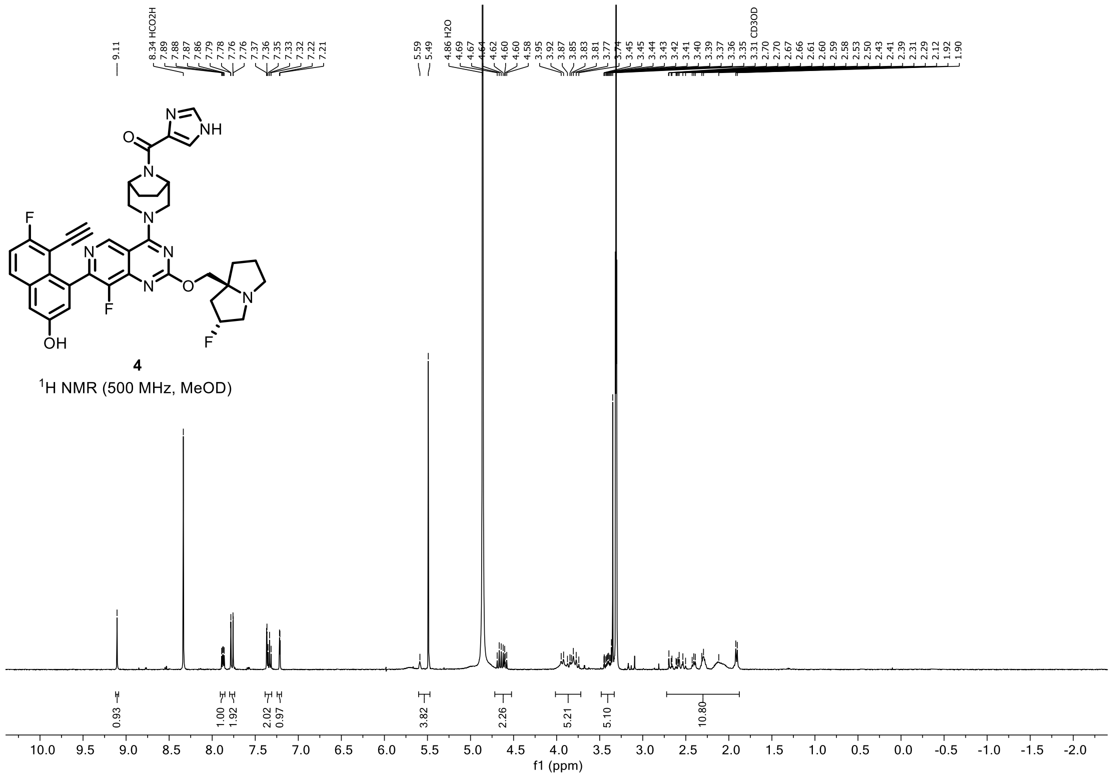


**4**
$^1$H NMR (500 MHz, MeOD)

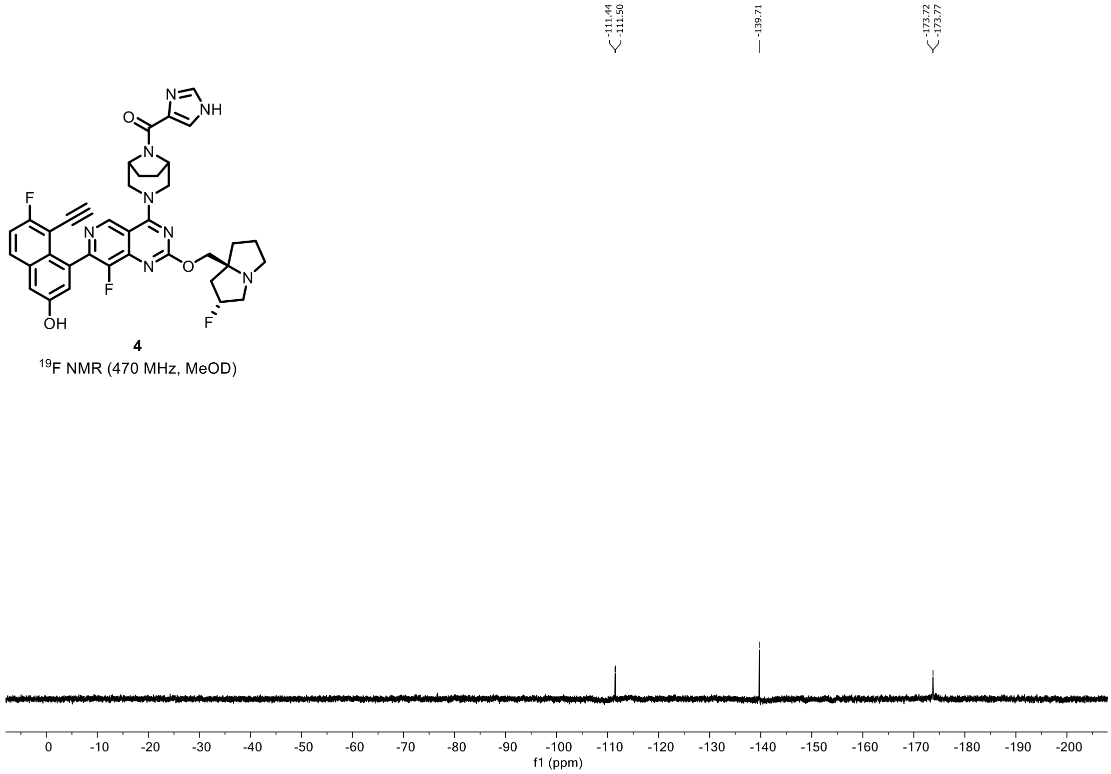


**4**
$^{19}$F NMR (470 MHz, MeOD)

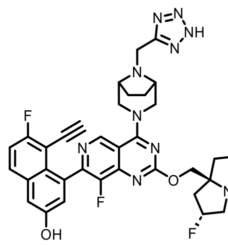


**5**

$^{1}$H NMR (500 MHz, MeOD)

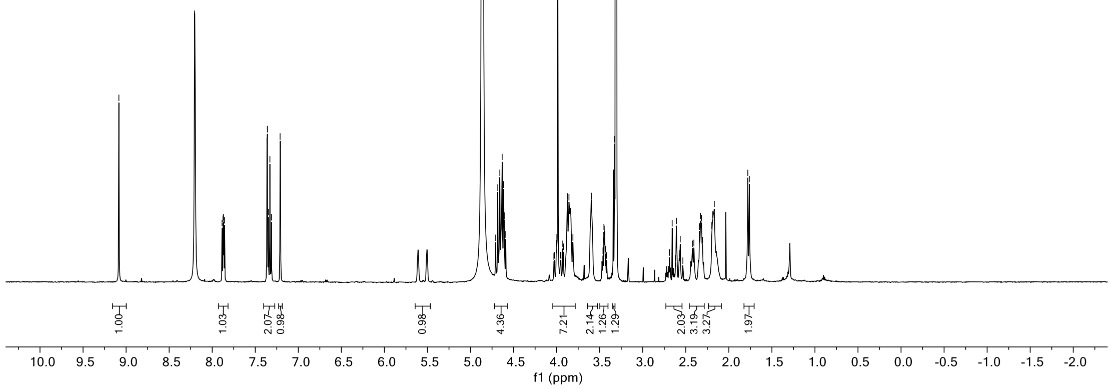


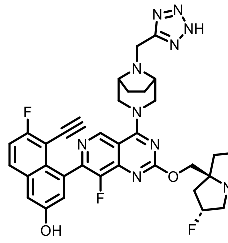


**5**

$^{19}$F NMR (470 MHz, MeOD)

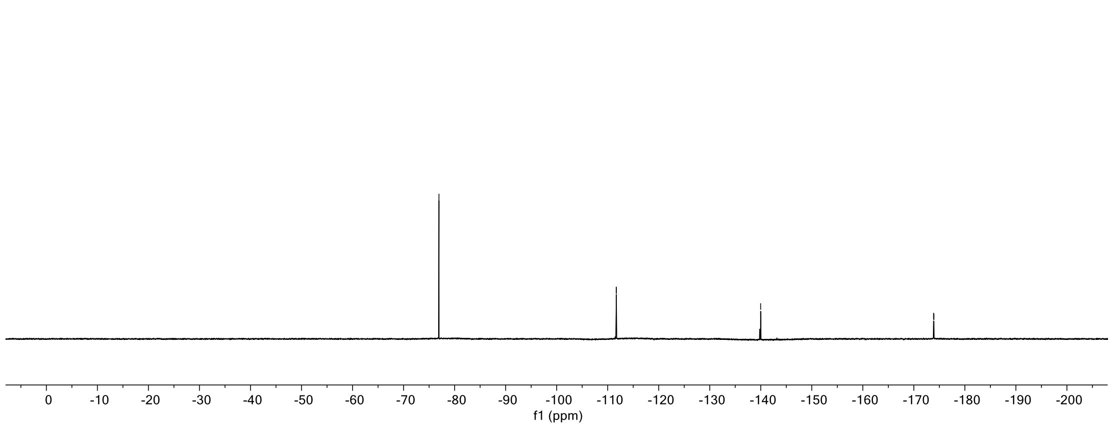

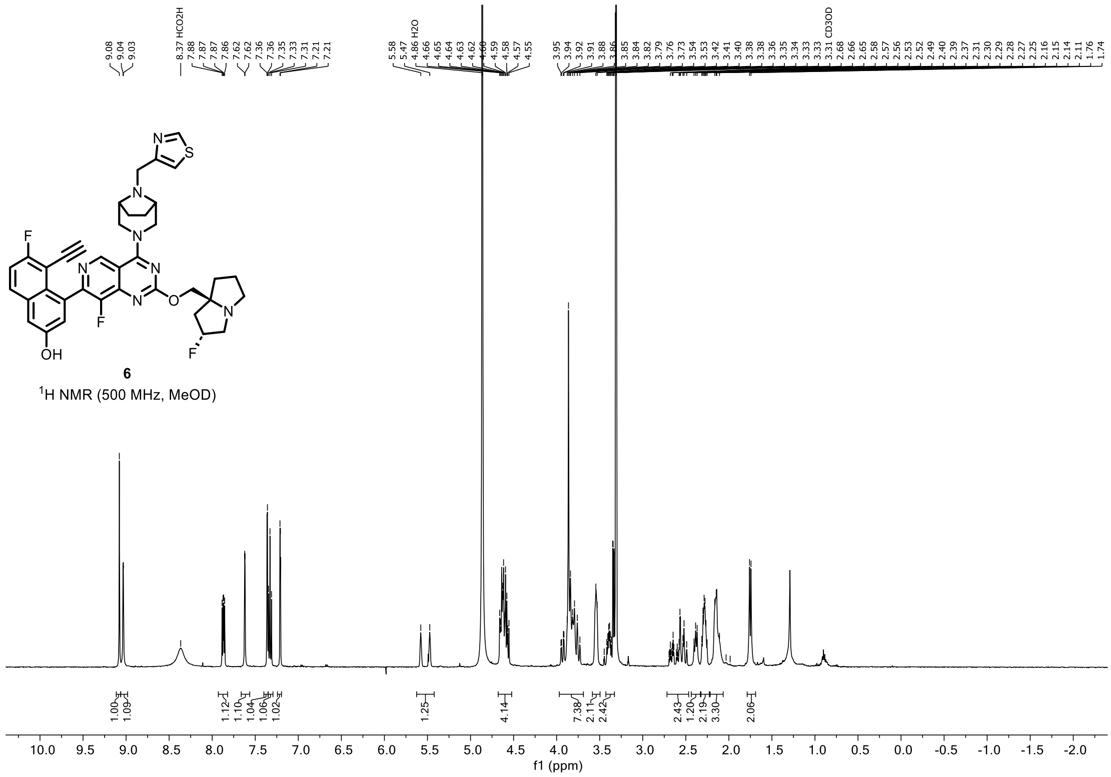

6

[1]H NMR (500 MHz, MeOD)

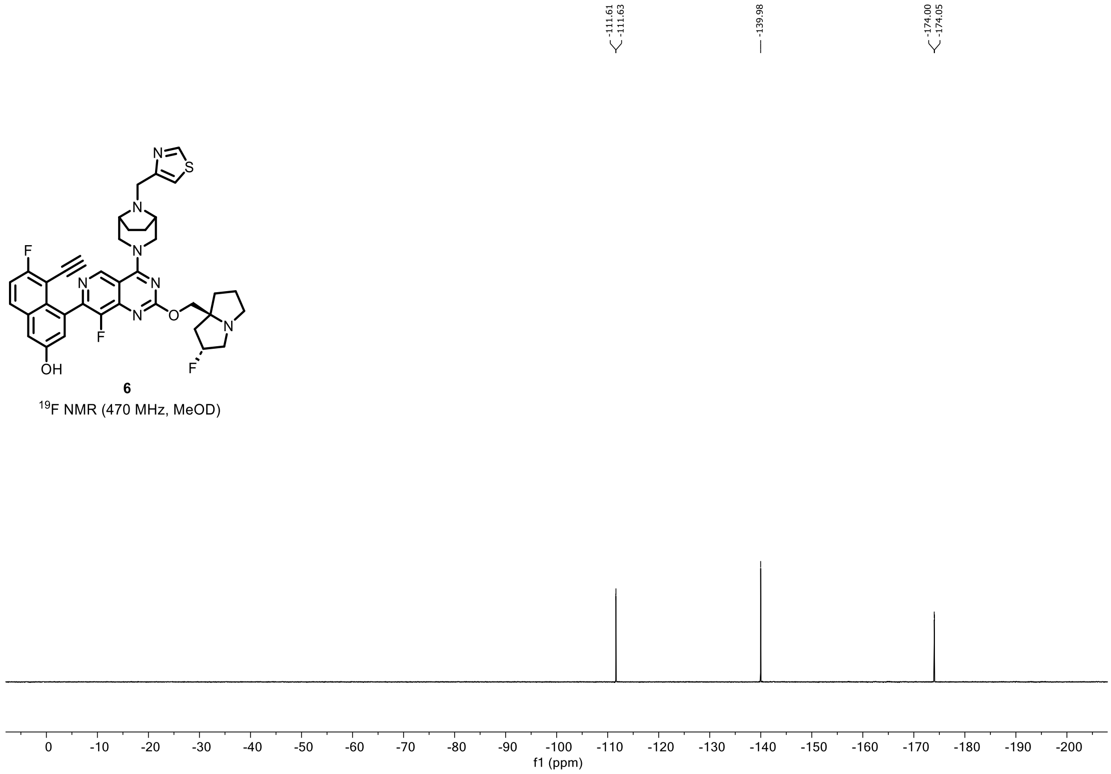

6

[19]F NMR (470 MHz, MeOD)

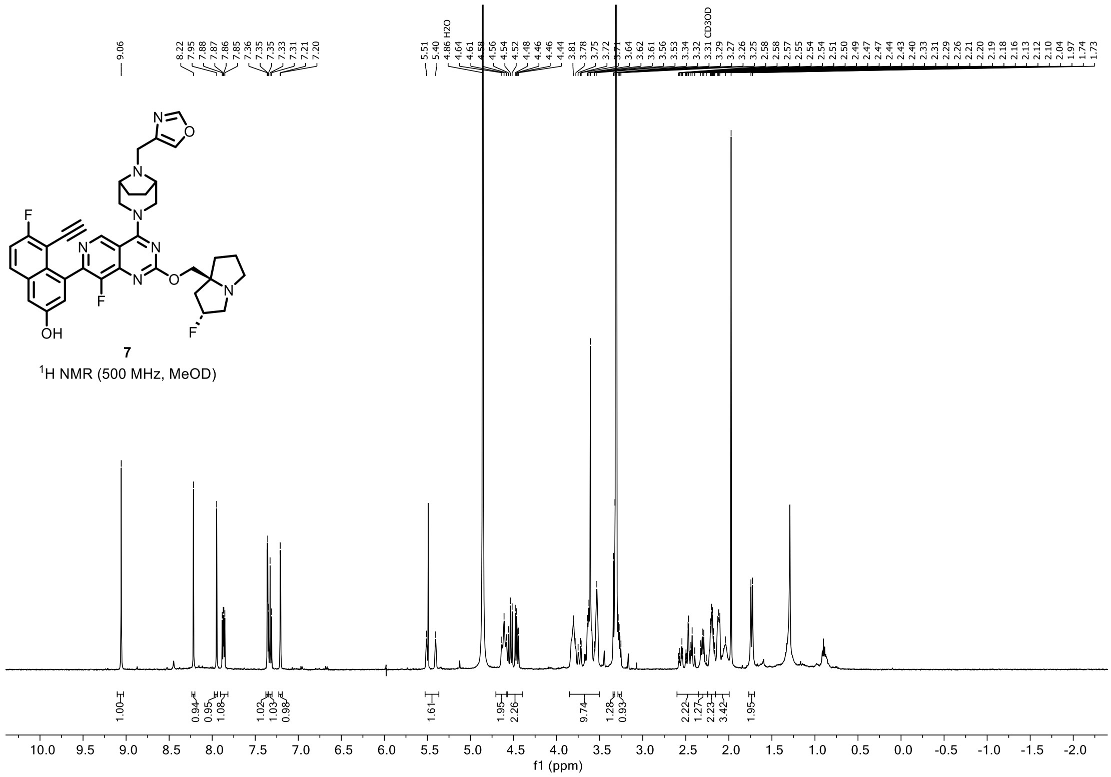


**7**

$^{1}$H NMR (500 MHz, MeOD)

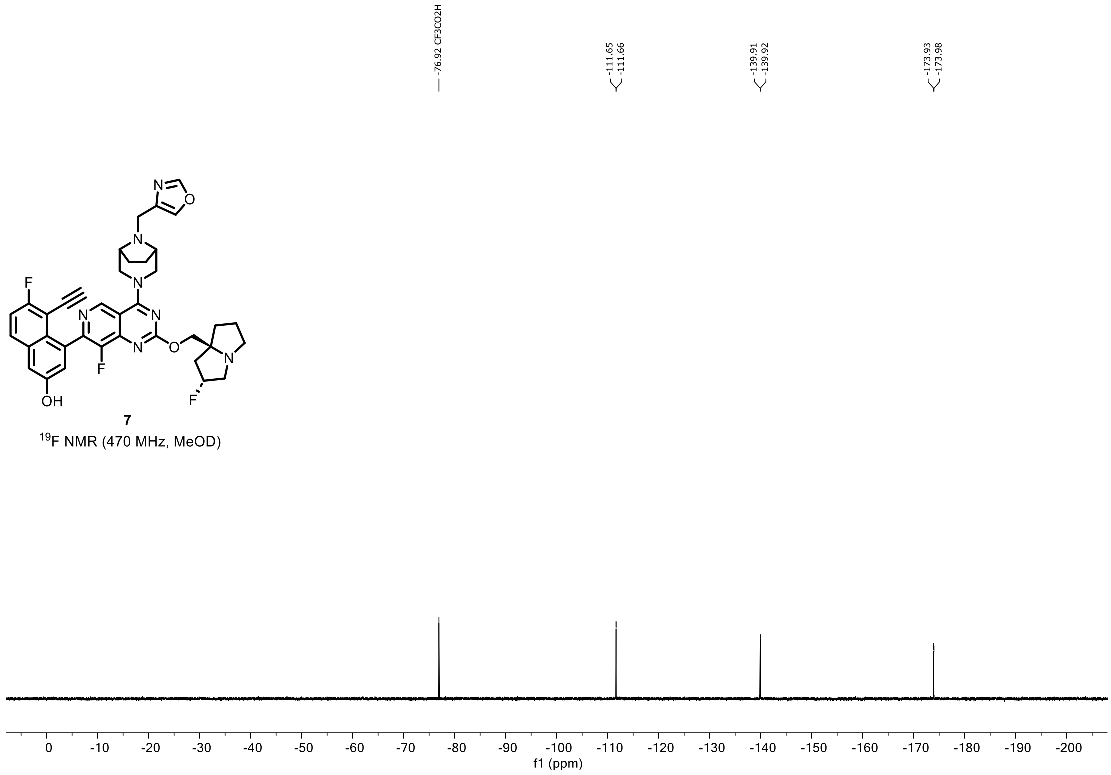


**7**

$^{19}$F NMR (470 MHz, MeOD)

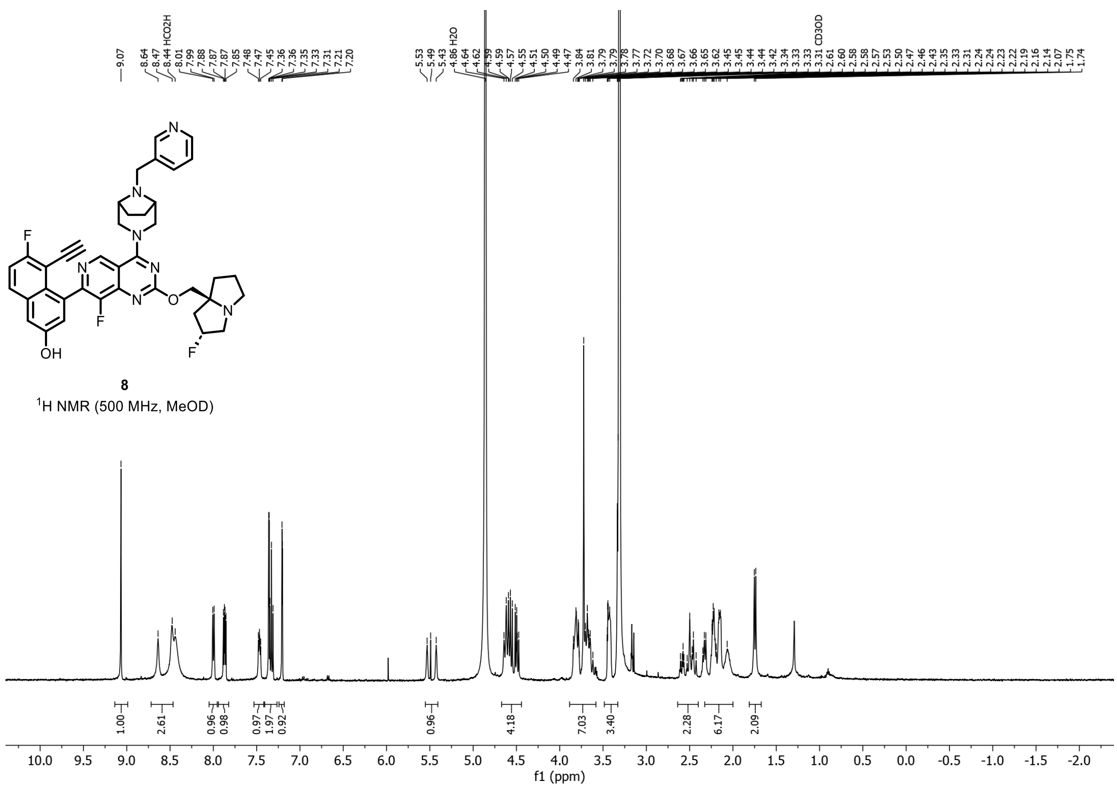

**8**

$^{1}H$ NMR (500 MHz, MeOD)

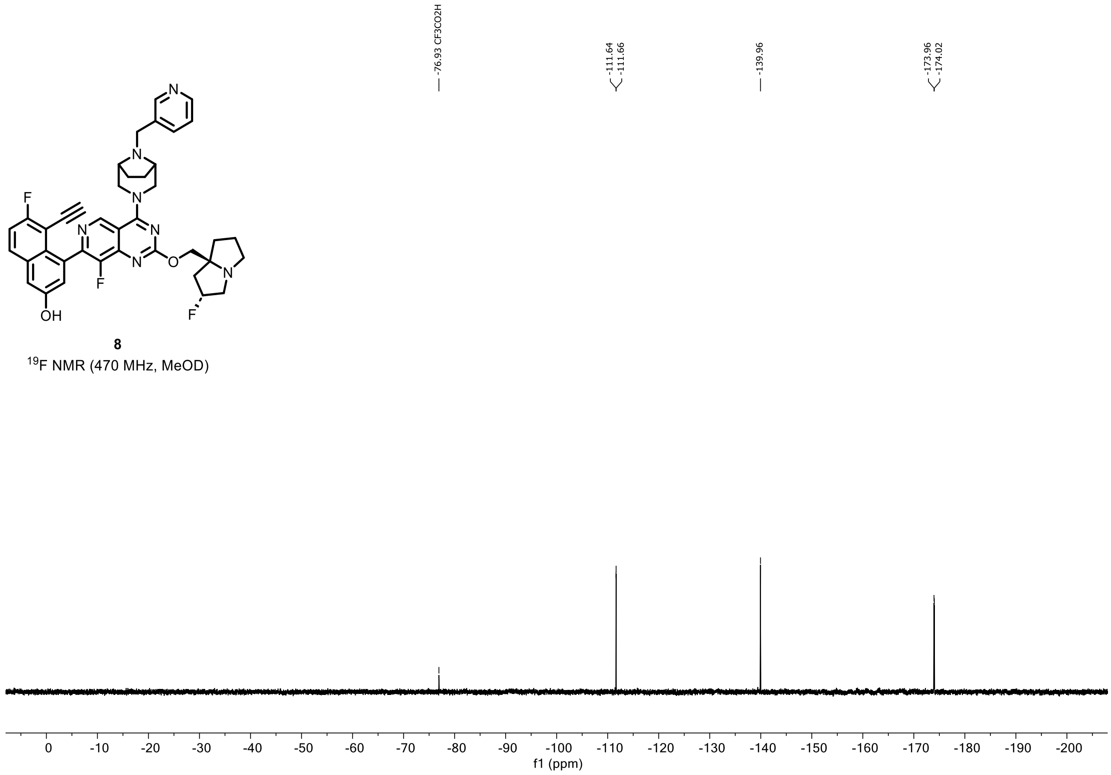

**8**

$^{19}F$ NMR (470 MHz, MeOD)

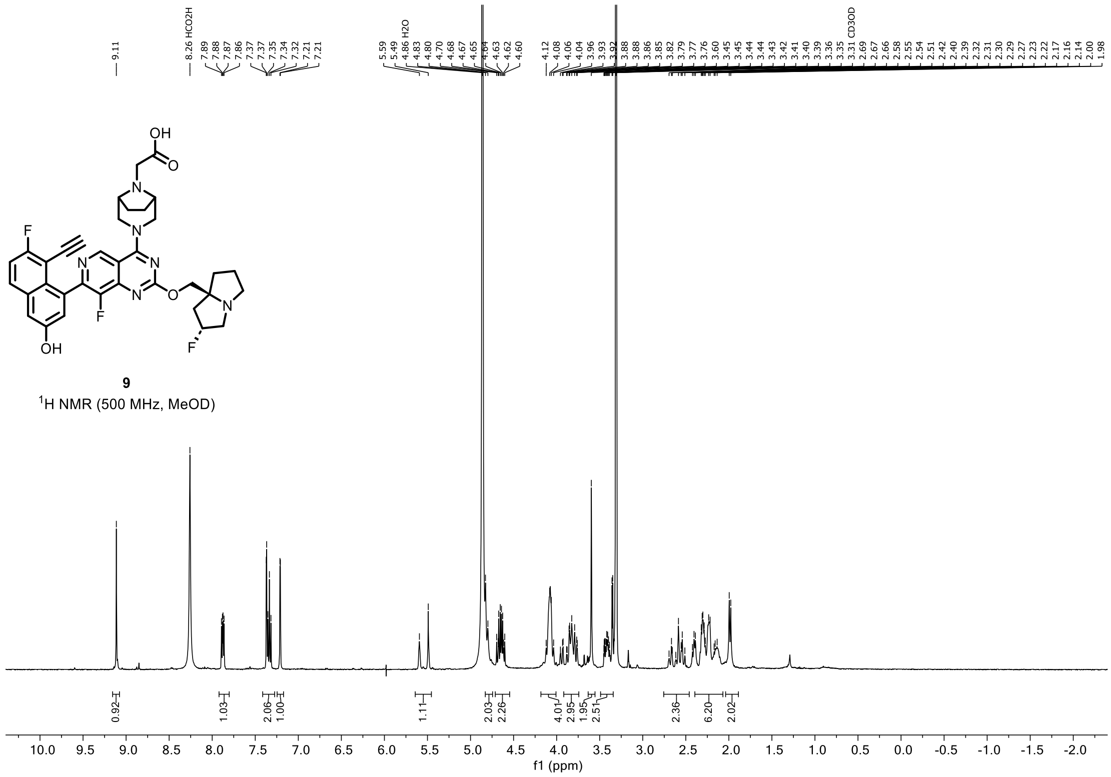
9.11
8.26 HCO2H
7.89 7.88 7.87 7.86 7.37 7.37 7.35 7.34 7.32 7.21 7.21
5.59 5.49 4.86 H2O 4.83 4.80 4.70 4.68 4.67 4.65 4.64 4.63 4.62 4.60
4.12 4.08 4.06 4.04 3.96 3.93 3.92 3.88 3.88 3.86 3.85 3.82 3.79 3.77 3.76 3.60 3.45 3.45 3.44 3.44 3.43 3.42 3.41 3.40 3.39 3.36 3.35 3.31 CD3OD 2.69 2.67 2.66 2.58 2.55 2.54 2.51 2.42 2.40 2.39 2.32 2.31 2.30 2.29 2.27 2.23 2.22 2.17 2.16 2.14 2.00 1.98
0.92 1.03 2.06 1.00 1.11 2.03 2.26 4.01 2.95 1.95 2.51 2.36 6.20 2.02
f1 (ppm)
9
1H NMR (500 MHz, MeOD)

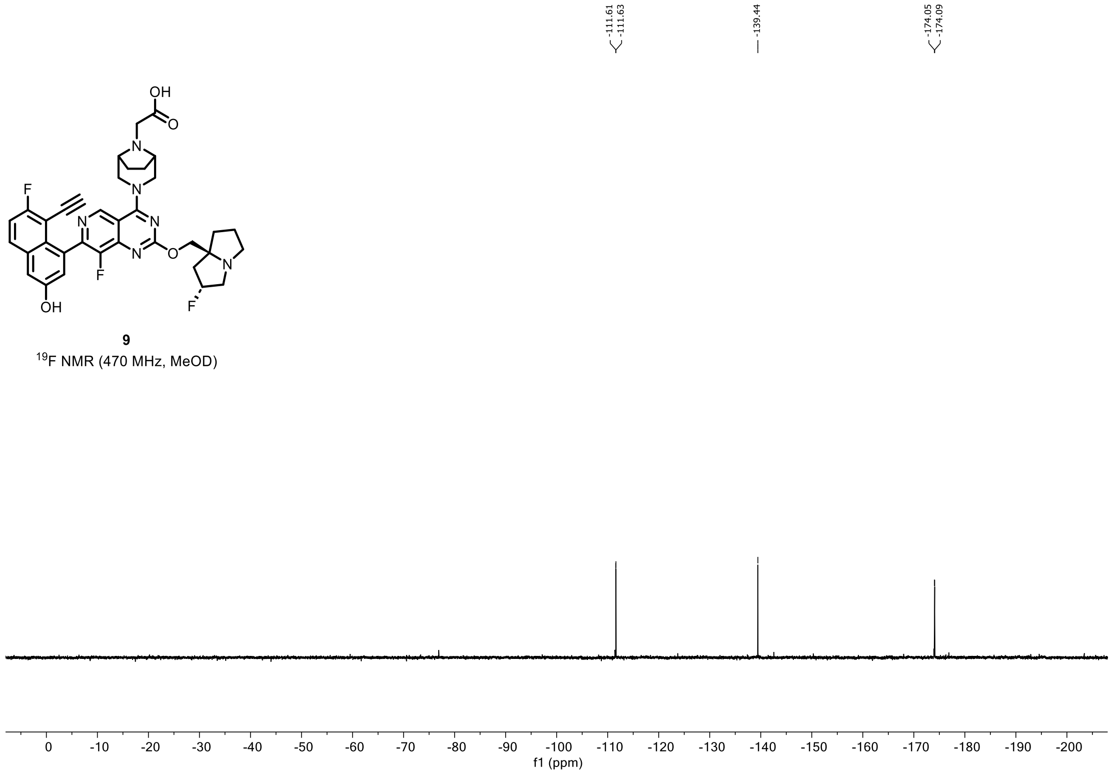
-111.61 -111.63
-139.44
-174.05 -174.09
f1 (ppm)
9
19F NMR (470 MHz, MeOD)

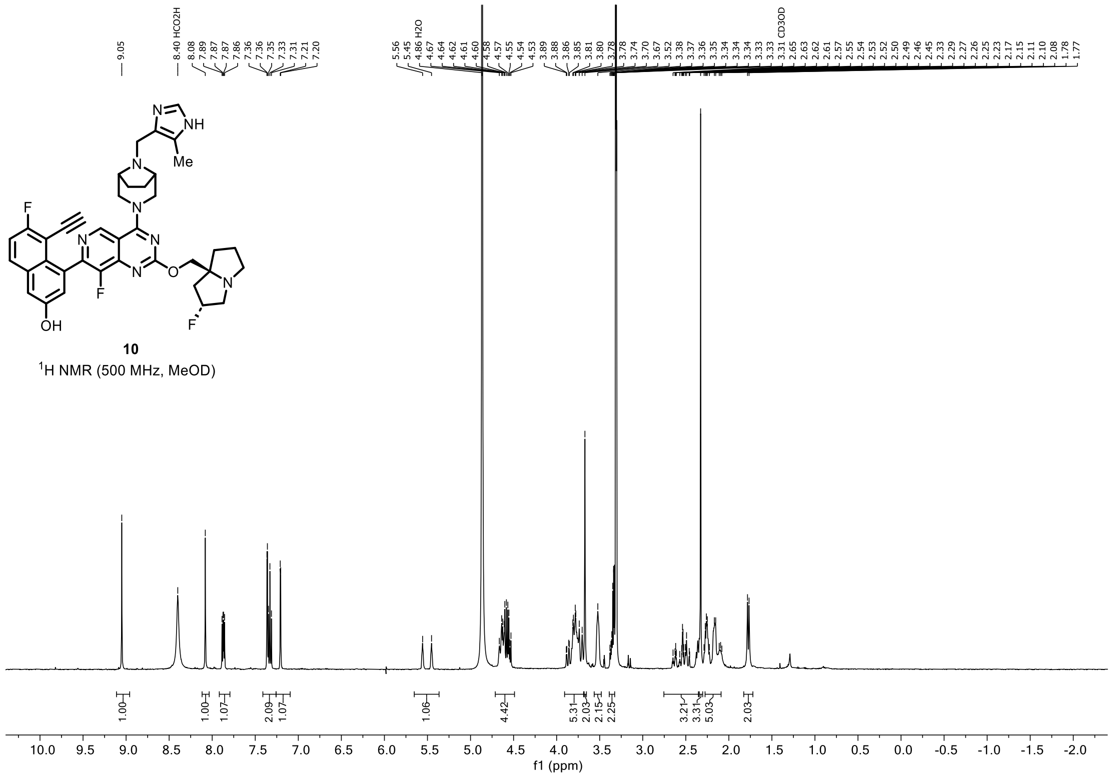


**10**
$^{1}H$ NMR (500 MHz, MeOD)

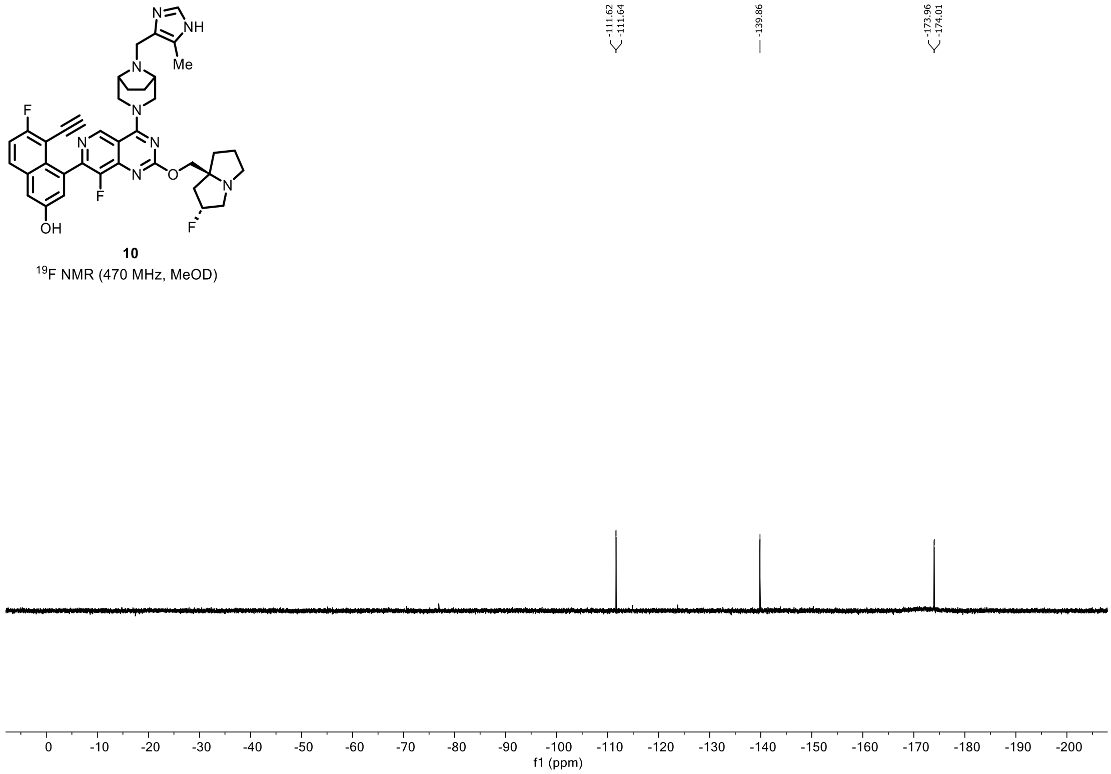


**10**
$^{19}F$ NMR (470 MHz, MeOD)

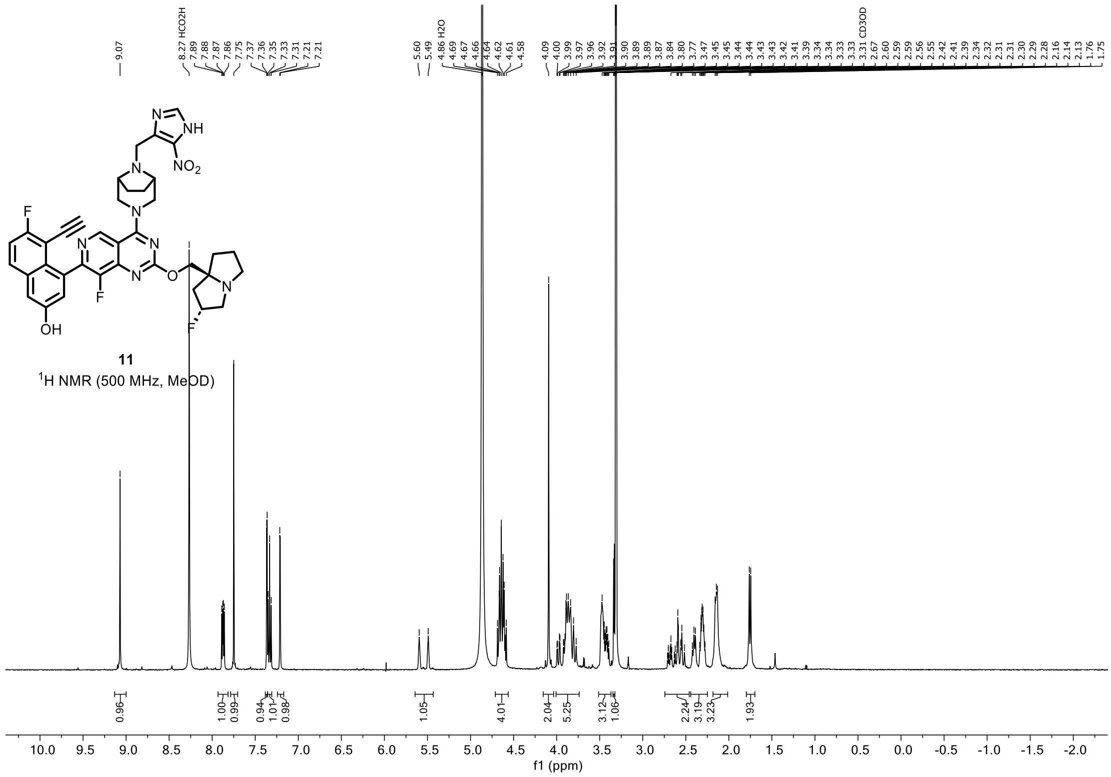
11
$^{1}$H NMR (500 MHz, MeOD)
f1 (ppm)

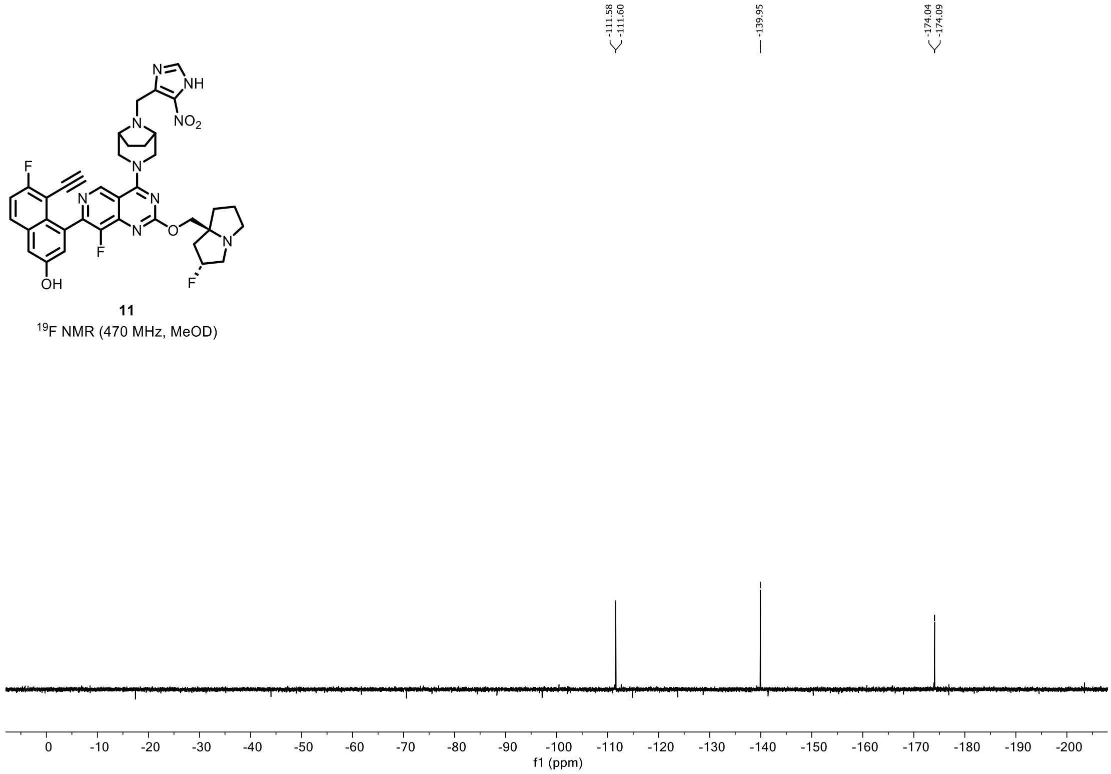
11
$^{19}$F NMR (470 MHz, MeOD)
f1 (ppm)

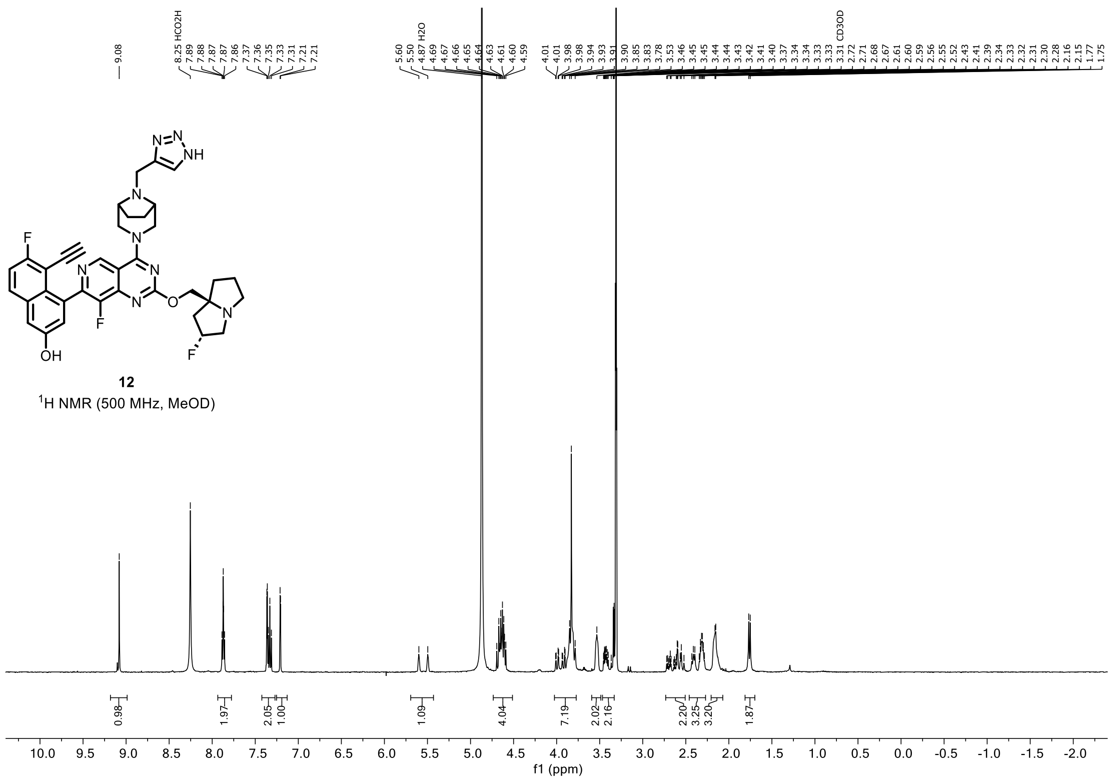


**12**

$^1$H NMR (500 MHz, MeOD)

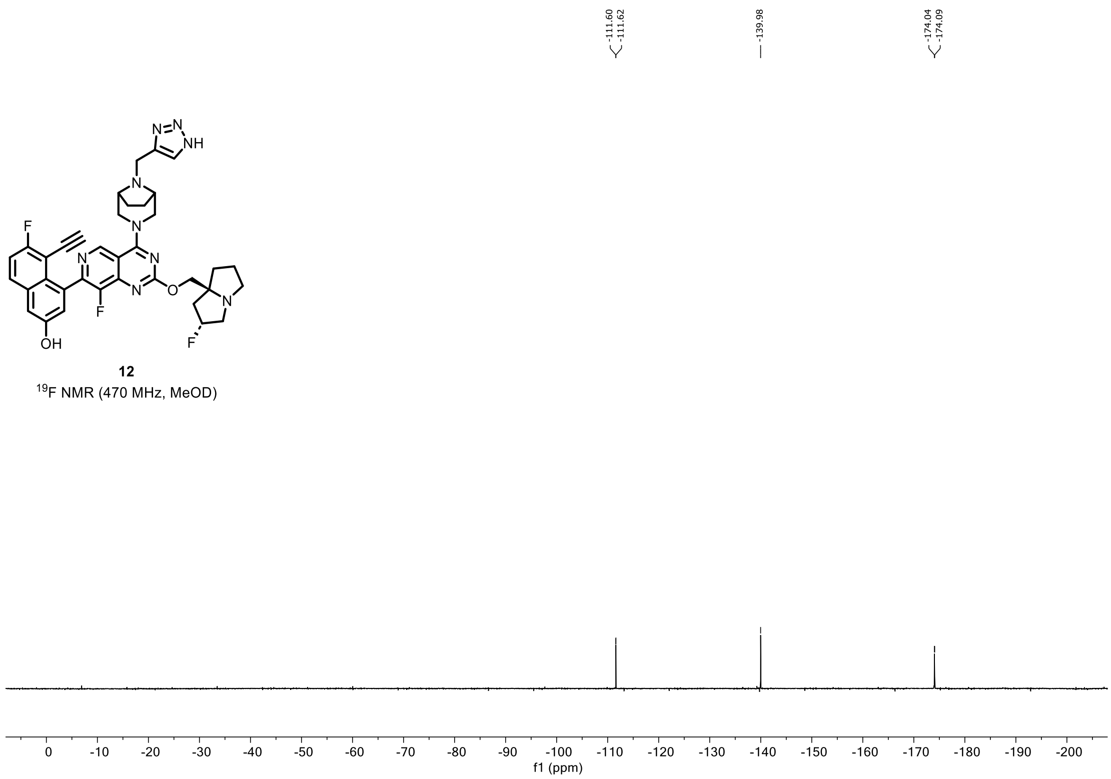


**12**

$^{19}$F NMR (470 MHz, MeOD)

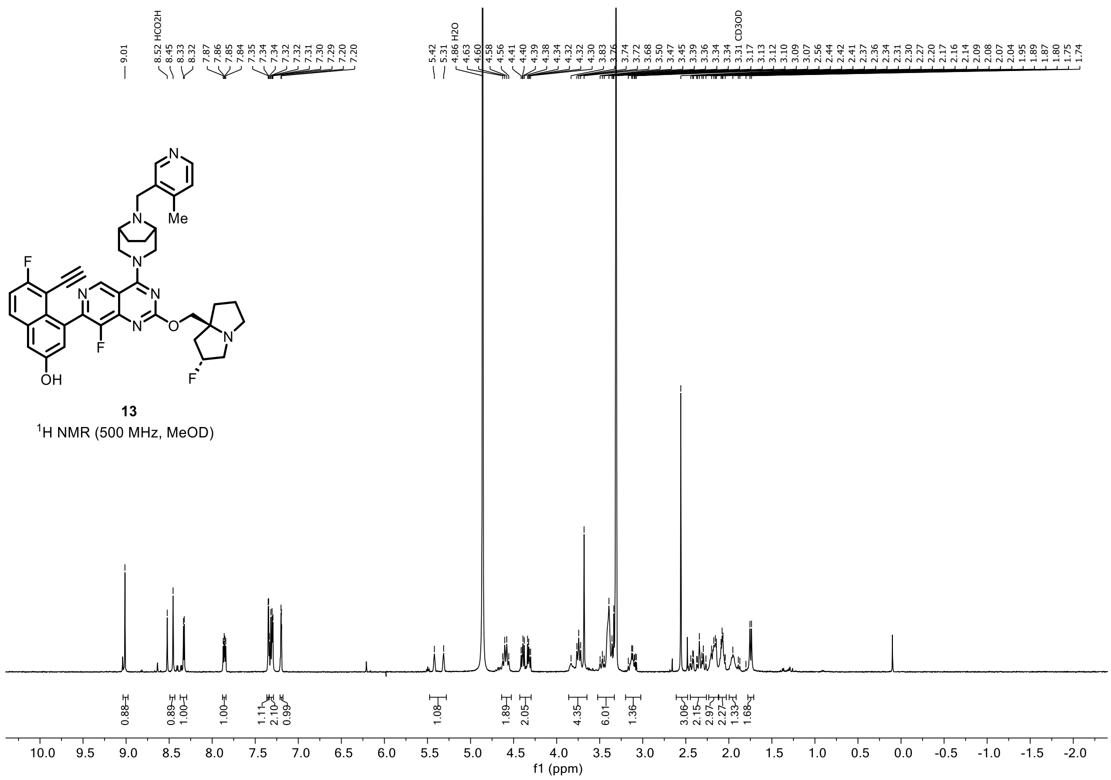
F
N
N
Me
N
N
O
N
F
OH
F
13
$^1$H NMR (500 MHz, MeOD)
10.0 9.5 9.0 8.5 8.0 7.5 7.0 6.5 6.0 5.5 5.0 4.5 4.0 3.5 3.0 2.5 2.0 1.5 1.0 0.5 0.0 -0.5 -1.0 -1.5 -2.0
f1 (ppm)

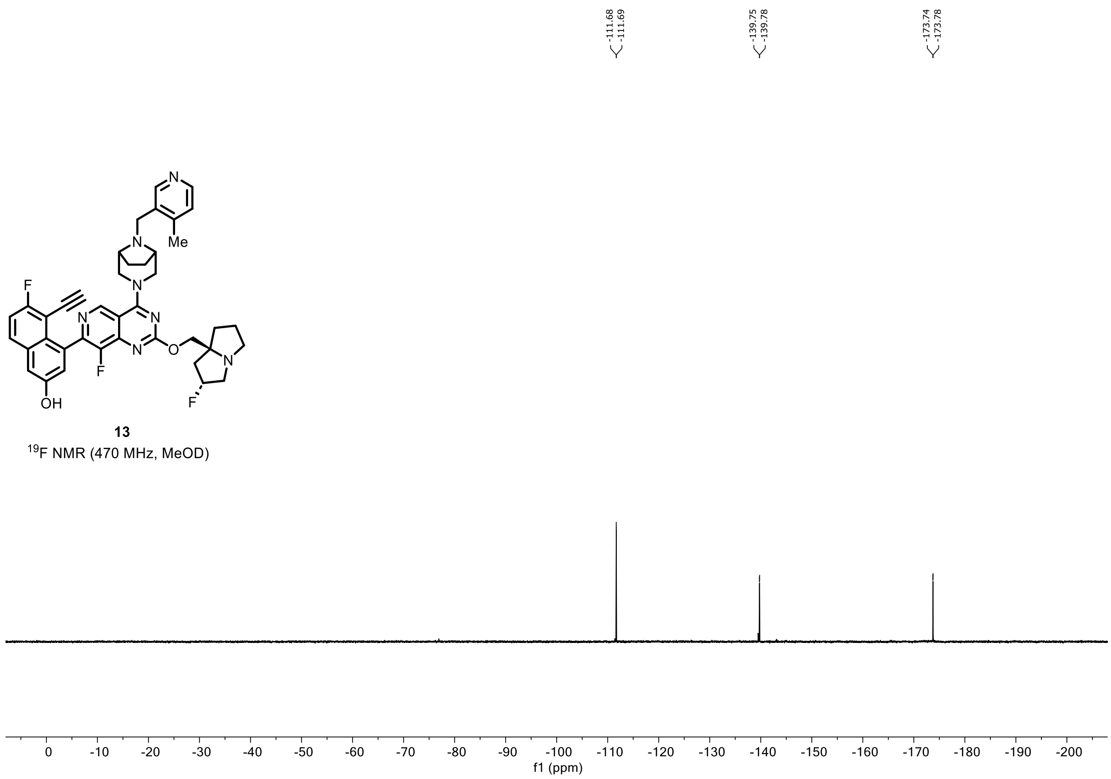
F
N
N
Me
N
N
O
N
F
OH
F
13
$^{19}$F NMR (470 MHz, MeOD)
0 -10 -20 -30 -40 -50 -60 -70 -80 -90 -100 -110 -120 -130 -140 -150 -160 -170 -180 -190 -200
f1 (ppm)

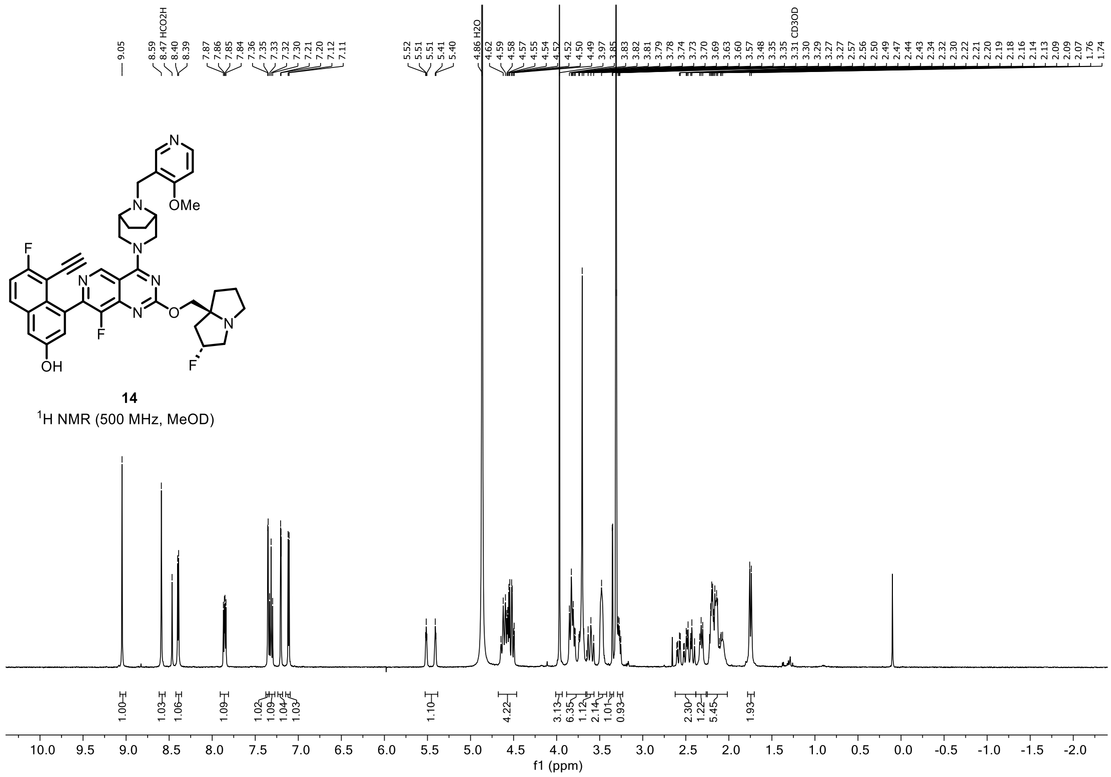


**14**
$^1$H NMR (500 MHz, MeOD)

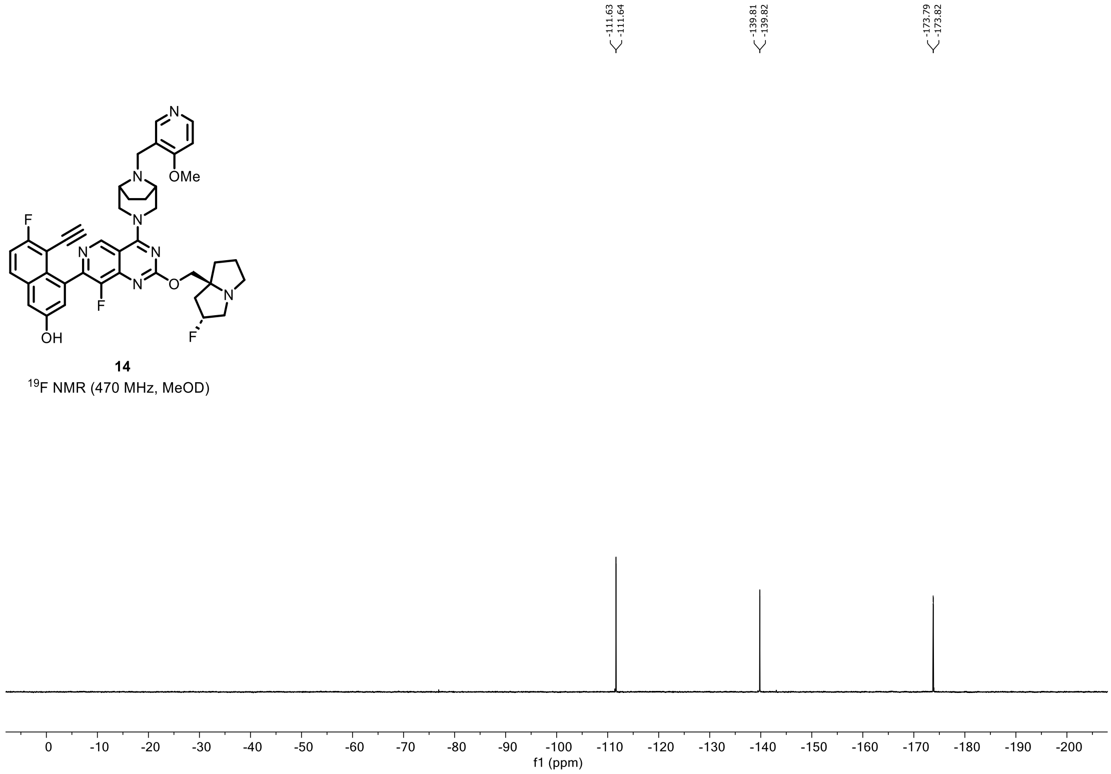


**14**
$^{19}$F NMR (470 MHz, MeOD)

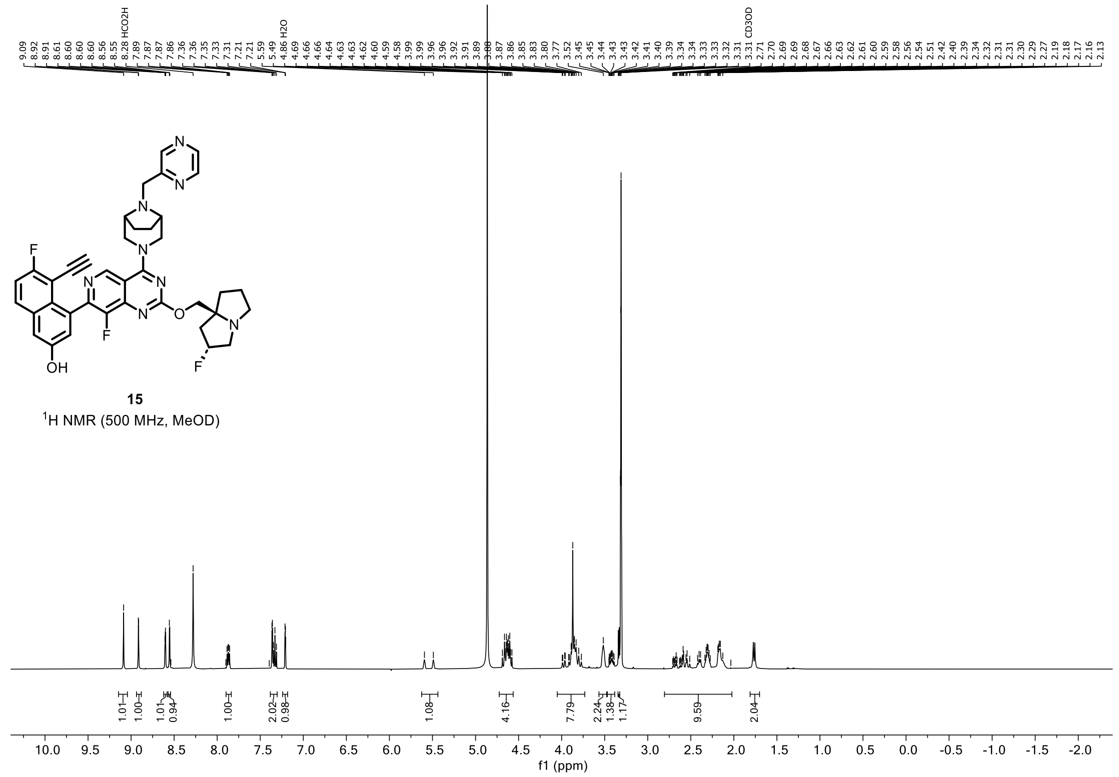
15
$^1$H NMR (500 MHz, MeOD)
f1 (ppm)

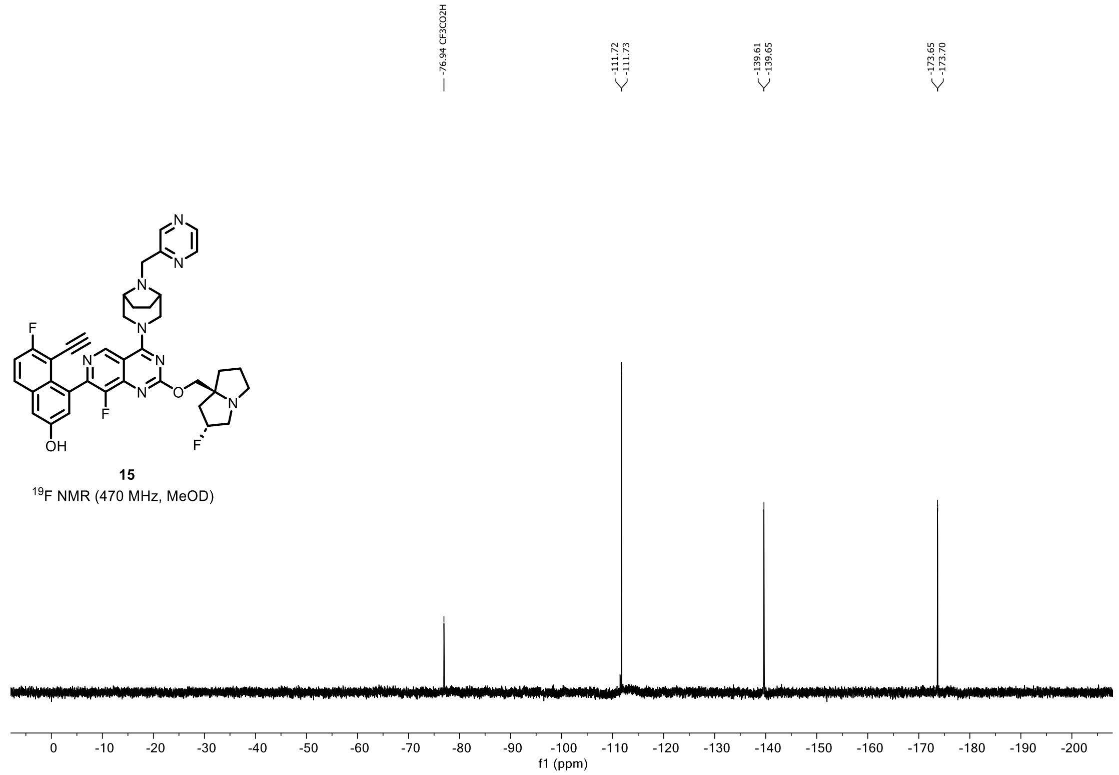
15
$^{19}$F NMR (470 MHz, MeOD)
f1 (ppm)

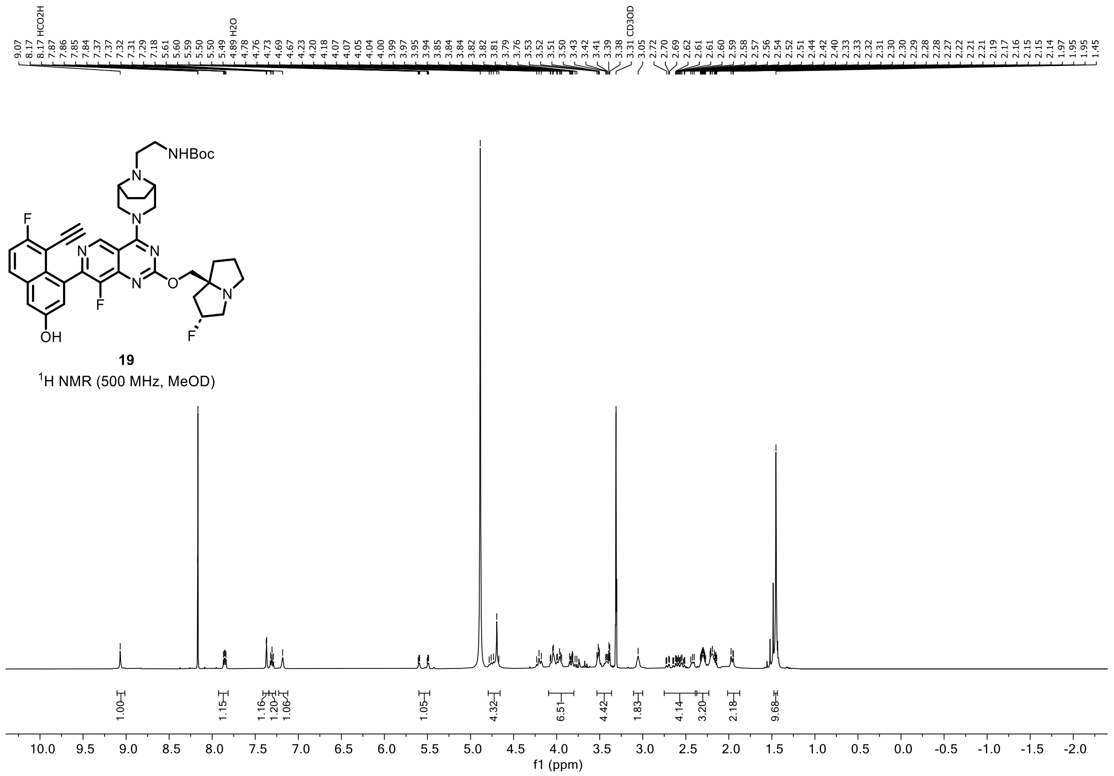


**19**
$^{1}$H NMR (500 MHz, MeOD)

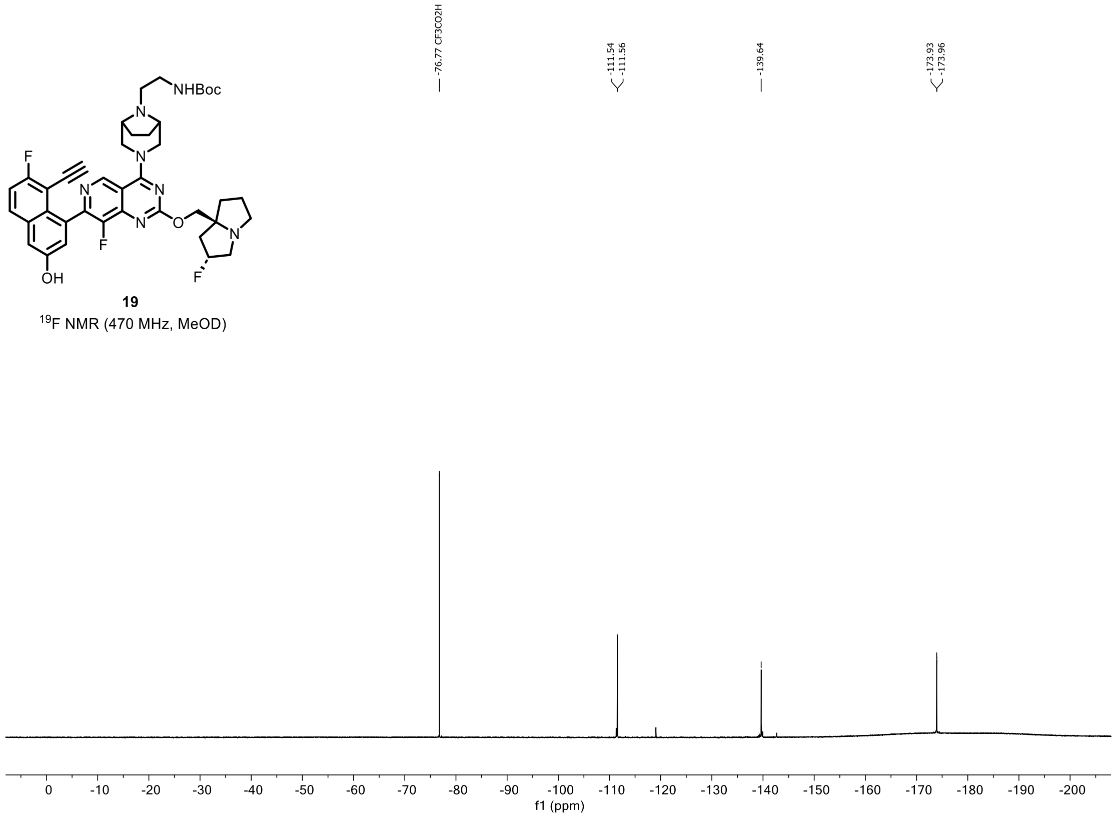


**19**
$^{19}$F NMR (470 MHz, MeOD)

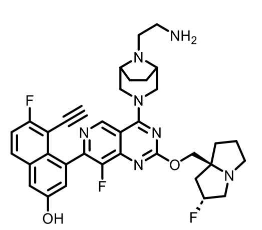


**16**

$^{1}$H NMR (500 MHz, MeOD)

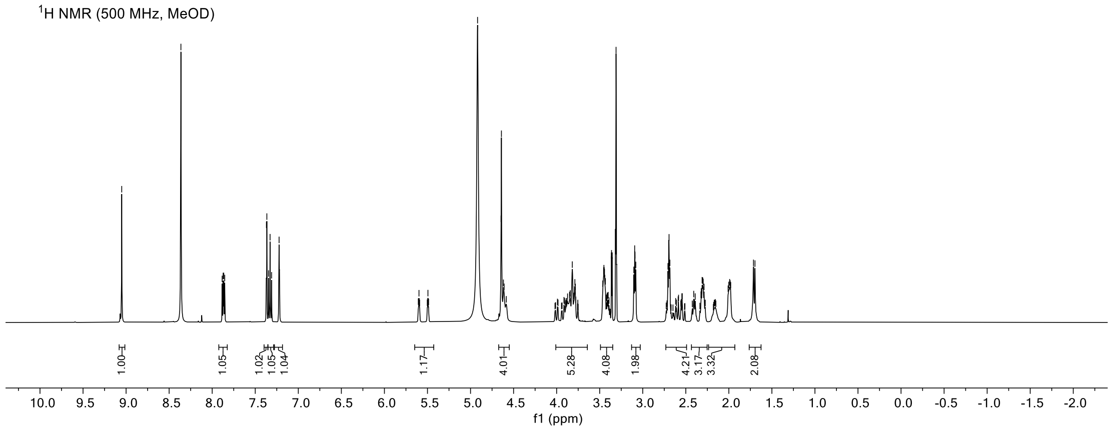


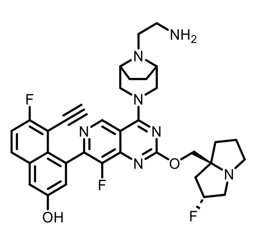


**16**

$^{19}$F NMR (470 MHz, MeOD)

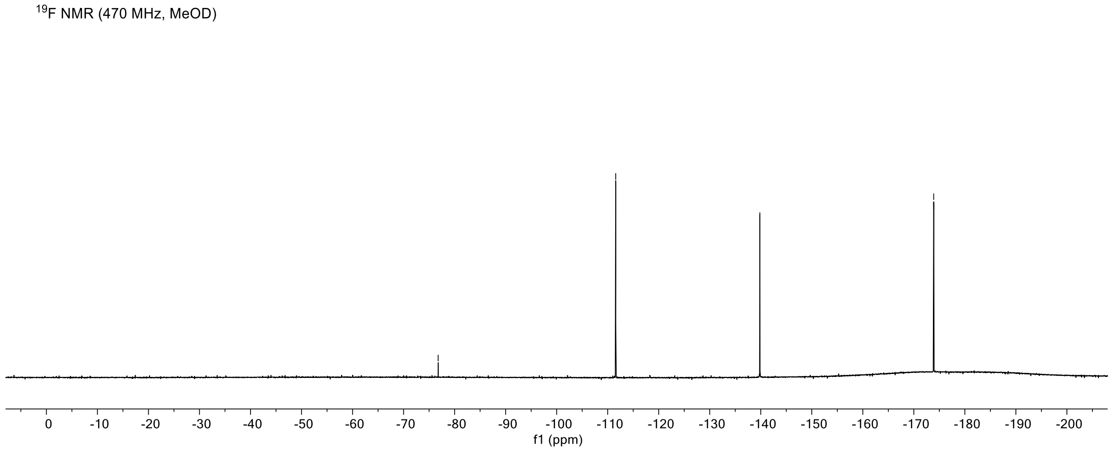

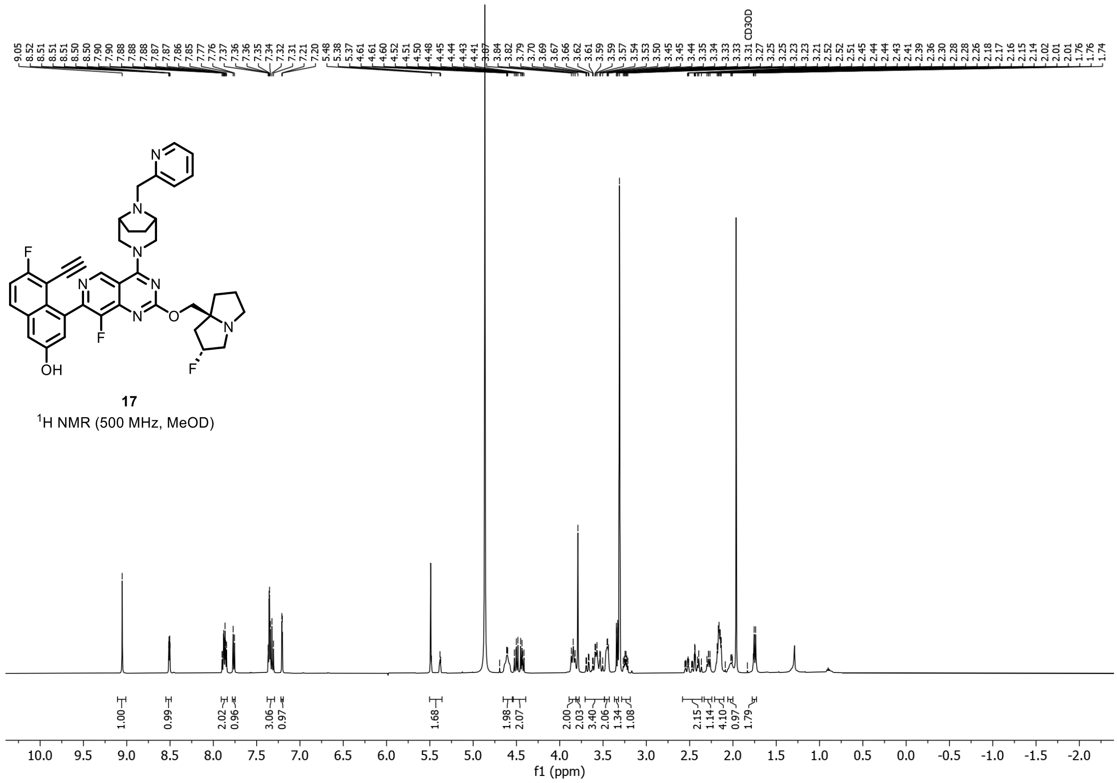


**17**
$^{1}H$ NMR (500 MHz, MeOD)

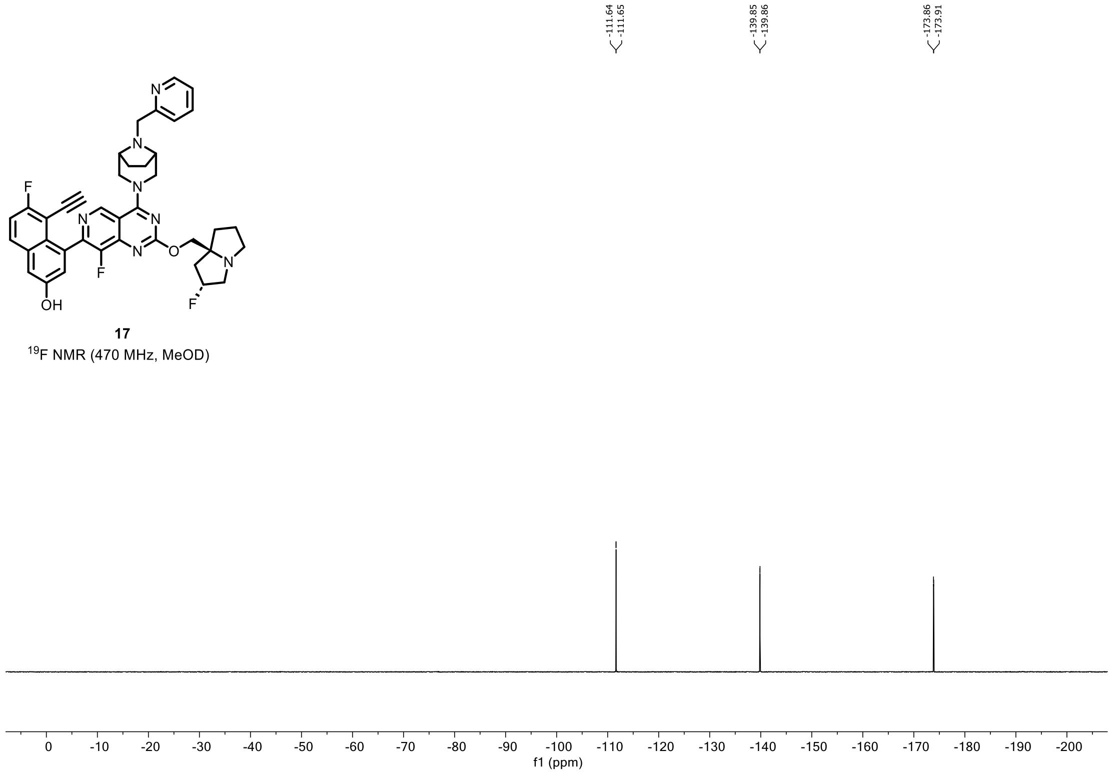


**17**
$^{19}F$ NMR (470 MHz, MeOD)

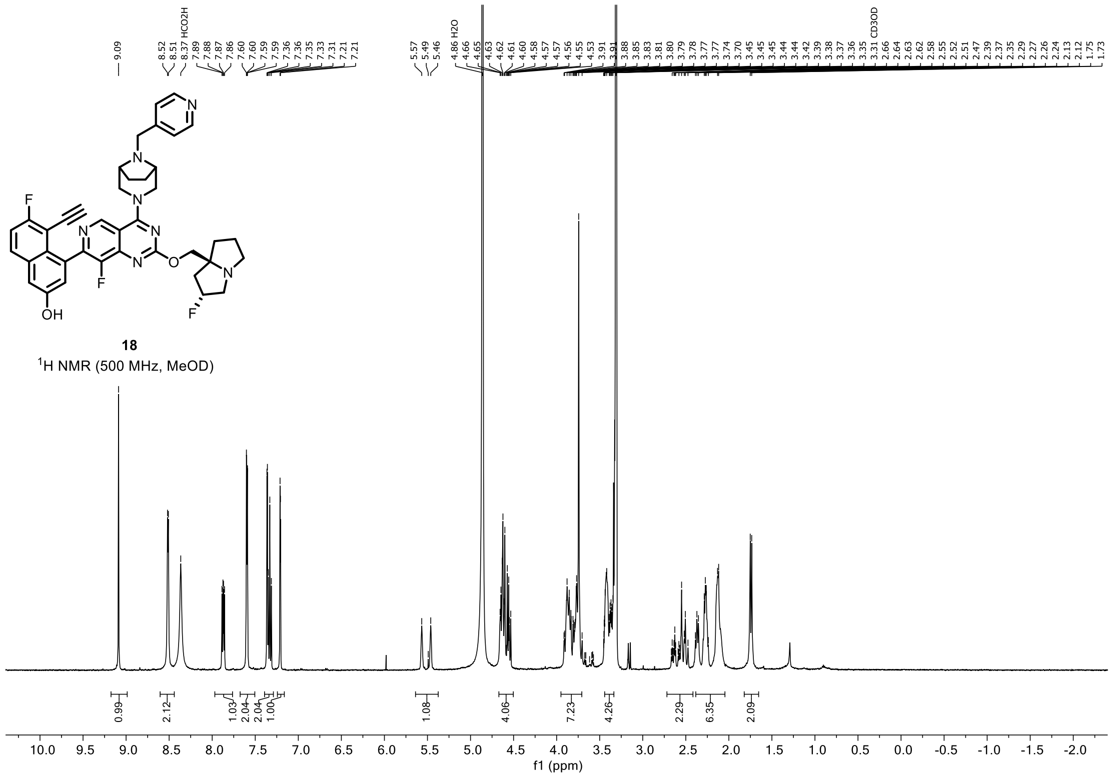

9.09
8.52
8.51
8.37 HCO2H
7.89
7.88
7.87
7.86
7.60
7.60
7.59
7.59
7.36
7.36
7.35
7.33
7.31
7.21
7.21
5.57
5.49
5.46
4.86 H2O
4.66
4.65
4.63
4.62
4.61
4.60
4.58
4.57
4.57
4.56
4.55
4.53
3.91
3.91
3.88
3.85
3.83
3.81
3.80
3.79
3.78
3.77
3.74
3.70
3.45
3.45
3.45
3.44
3.42
3.39
3.38
3.37
3.36
3.35
3.31 CD3OD
2.66
2.64
2.63
2.62
2.58
2.55
2.52
2.51
2.47
2.39
2.37
2.35
2.29
2.27
2.26
2.24
2.13
2.12
1.75
1.73
18
$^{1}$H NMR (500 MHz, MeOD)
0.99
2.12
1.03
2.04
2.04
1.00
1.08
4.06
7.23
4.26
2.29
6.35
2.09
10.0 9.5 9.0 8.5 8.0 7.5 7.0 6.5 6.0 5.5 5.0 4.5 4.0 3.5 3.0 2.5 2.0 1.5 1.0 0.5 0.0 -0.5 -1.0 -1.5 -2.0
f1 (ppm)


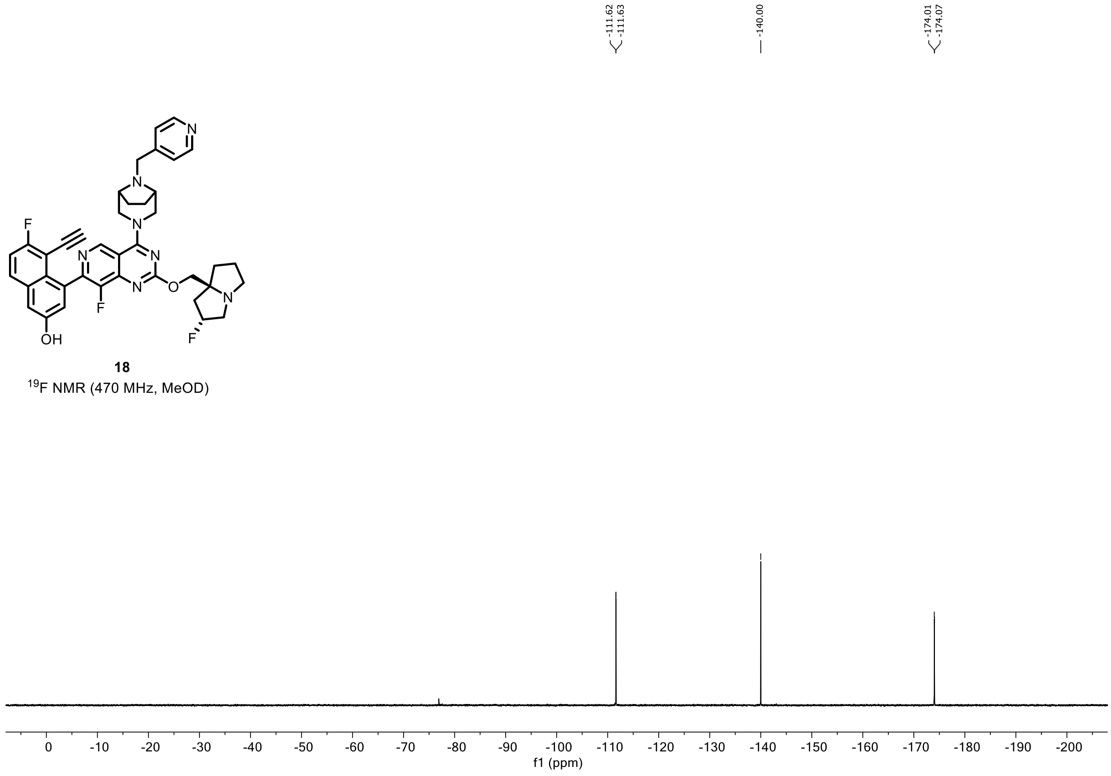

-111.62
-111.63
-140.00
-174.01
-174.07
18
$^{19}$F NMR (470 MHz, MeOD)
0 -10 -20 -30 -40 -50 -60 -70 -80 -90 -100 -110 -120 -130 -140 -150 -160 -170 -180 -190 -200
f1 (ppm)

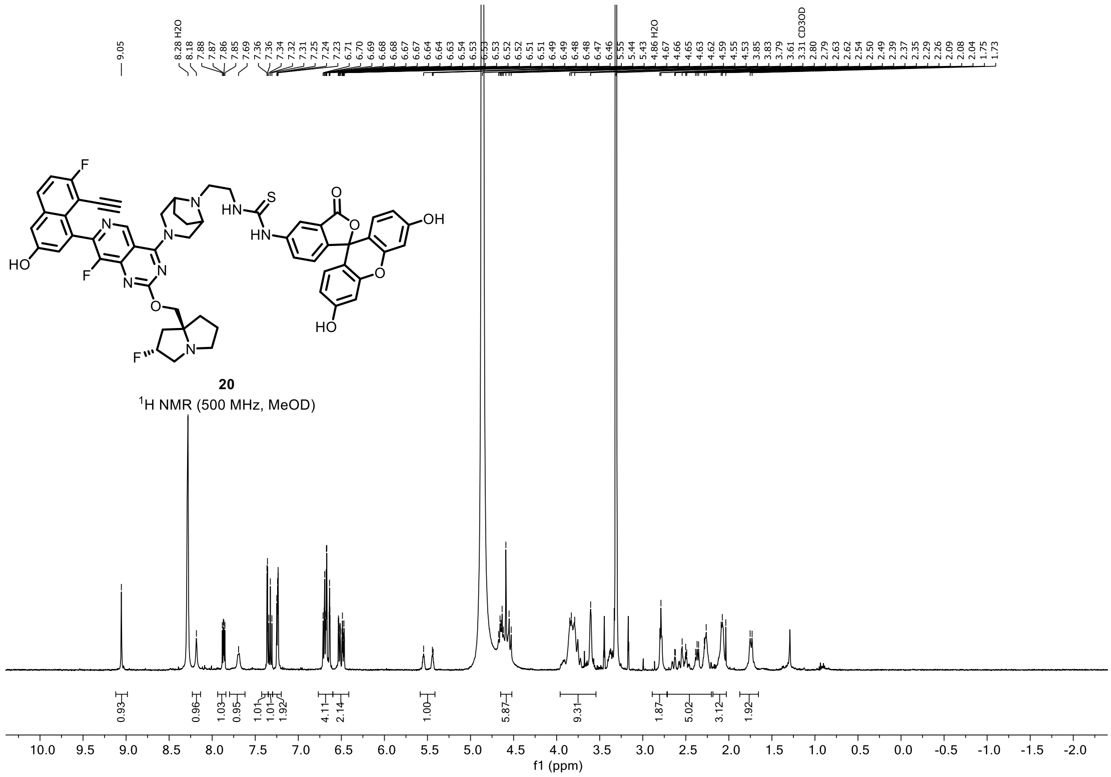


**20**

$^1$H NMR (500 MHz, MeOD)

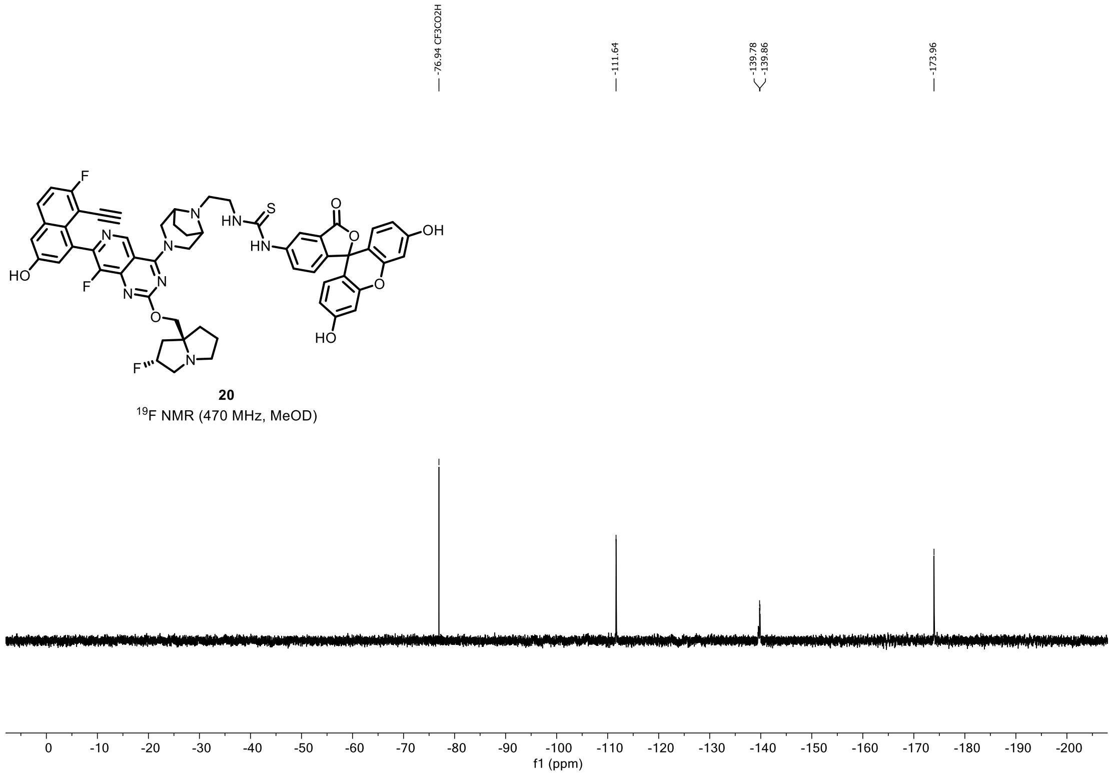


**20**

$^{19}$F NMR (470 MHz, MeOD)

**Part 6. Reference.**

[1] Pan, X.; Wang, H.; Li, C.; Zhang, J. Z. H.; Ji, C. MolGpka: A Web Server for Small Molecule p $K_a$ Prediction Using a Graph-Convolutional Neural Network. *J. Chem. Inf. Model.* **2021**, *61* (7), 3159–3165. https://doi.org/10.1021/acs.jcim.1c00075.
